# Transcriptome and chromatin structure annotation of liver, CD4+ and CD8+ T cells from four livestock species

**DOI:** 10.1101/316091

**Authors:** Sylvain Foissac, Sarah Djebali, Kylie Munyard, Nathalie Vialaneix, Andrea Rau, Kevin Muret, Diane Esquerré, Matthias Zytnicki, Thomas Derrien, Philippe Bardou, Fany Blanc, Cèdric Cabau, Elisa Crisci, Sophie Dhorne-Pollet, Françoise Drouet, Thomas Faraut, Ignacio Gonzalez, Adeline Goubil, Sonia Lacroix-Lamandé, Fabrice Laurent, Sylvain Marthey, Maria Marti-Marimon, Raphaelle Momal-Leisenring, Florence Mompart, Pascale Quéré, David Robelin, Magali San Cristobal, Gwenola Tosser-Klopp, Silvia Vincent-Naulleau, Stéphane Fabre, Marie-Hélène Pinard-Van der Laan, Christophe Klopp, Michelè Tixier-Boichard, Hervé Acloque, Sandrine Lagarrigue, Elisabetta Giuffra

## Abstract

**Background:** Functional annotation of livestock genomes is a critical step to decipher the genotype-to-phenotype relationship underlying complex traits. As part of the Functional Annotation of Animal Genomes (FAANG) action, the FR-AgENCODE project (http://www.fragencode.org) aimed to profile the landscape of transcription (RNA-seq), chromatin accessibility (ATAC-seq) and conformation (Hi-C) in four livestock species representing ruminants (cattle, goat), monogastrics (pig) and birds (chicken), using three target samples related to metabolism (liver) and immunity (CD4+ and CD8+ T cells).

**Results:** RNA-seq assays considerably extended the available catalog of annotated transcripts and identified differentially expressed genes with unknown function, including new syntenic lncRNAs. ATAC-seq highlighted an enrichment for transcription factor binding sites in differentially accessible regions of the chromatin. Comparative analyses revealed a core set of conserved regulatory regions across species. Topologically Associating Domains (TADs) and epigenetic A/B compartments annotated from Hi-C data were consistent with RNA-seq and ATAC-seq data. Multi-species comparisons showed that conserved TAD boundaries had stronger insulation properties than species-specific ones and that the genomic distribution of orthologous genes in A/B compartments was significantly conserved across species.

**Conclusions:** We report the first multi-species and multi-assay genome annotation results obtained by a FAANG project. Beyond the generation of reference annotations and the confirmation of previous findings on model animals, the integrative analysis of data from multiple assays and species sheds a new light on the multi-scale selective pressure shaping genome organization from birds to mammals. Overall, these results emphasize the value of FAANG for research on domesticated animals and reinforces the importance of future meta-analyses of the reference datasets being generated by this community on different species.

## Background

Most complex trait-associated loci lie outside protein-coding regions, and comparative genomics studies have shown that the majority of mammalian-conserved and recently adapted regions consist of non-coding elements [1–3]. This evidence prompted the first large-scale efforts into genome annotation for human and model organisms [4–6]. The genome-wide annotation maps generated by these projects helped to shed light on the main features of genome activity. For example, chromatin conformation or transcription factor occupancy at regulatory elements can often be directly tied to the biology of the specific cell or tissue under study [3, 7, 8]. Moreover, although a subset of core regulatory systems are largely conserved across humans and mice, the underlying regulatory systems often diverge substantially [9–11], implying that understanding the phenotypes of interest requires organism-specific information for any specific physiological phase, tissue and cell.

The Functional Annotation of Animal Genomes (FAANG) initiative (www.faang.org) aims to support and coordinate the community in the endeavor of creating reference functional maps of the genomes of domesticated animals across different species, tissues and developmental stages, with an initial focus on farm and companion animals [12–15]. To achieve high quality functional annotation of these genomes, FAANG carries out activities to standardize core biochemical assay protocols, coordinate and facilitate data sharing. It also establishes an infrastructure for the analysis of genotype-to-phenotype data (www.faang.org, www.faang-europe.org). Substantial efforts are being dedicated to farm animal species, as deciphering the genotype-to-phenotype relationships underlying complex traits such as production efficiency and disease resistance is a prerequisite for exploiting the full potential of livestock [12].

Here we report the main results of a pilot project (FR-AgENCODE, www.fragencode.org) launched at the beginning of the FAANG initiative. The broad aim was to generate standardized FAANG reference datasets from four livestock species (cattle, goat, chicken and pig) through the adaptation and optimization of molecular assays and analysis pipelines. We first collected a panel of samples from more than 40 tissues from two males and two females of four species: *Bos taurus* (cattle, Holstein breed), *Capra hircus* (goat, Alpine breed), *Gallus gallus* (chicken, White Leghorn breed) and *Sus scrofa* (pig, Large White breed), generating a total of 4,115 corresponding entries registered at the EMBL-EBI BioSamples database (see Methods). For molecular characterization, three tissues were chosen to represent a “hub” organ (liver) and two broad immune cell populations (CD4+ and CD8+ T cells). This allowed the acquisition of a partial representation of energy metabolism and immunity functions, as well as the optimization of the protocols for experimental assays for both tissue-dissociated and primary sorted cells. In addition to the transcriptome, we analyzed chromatin accessibility by the assay for transposase-accessible chromatin using sequencing (ATAC-seq, [16]), and we characterized the three-dimensional (3D) genome architecture by coupling proximity-based ligation with massively parallel sequencing (Hi-C, [17]) (Figure 1). Using this combination of tissues/assays, we assessed the expression of a large set of coding and non-coding transcripts in the four species, evaluated their patterns of differential expression in light of chromatin accessibility at promoter regions, and characterized active and inactive topological domains of these genomes. The integrative analysis showed a global consistency across all data, emphasizing the value of a coordinated action to improve the genomic annotation of livestock species, and revealed multiple layers of evolutionary conservation from birds to mammals.

**Figure 1.**
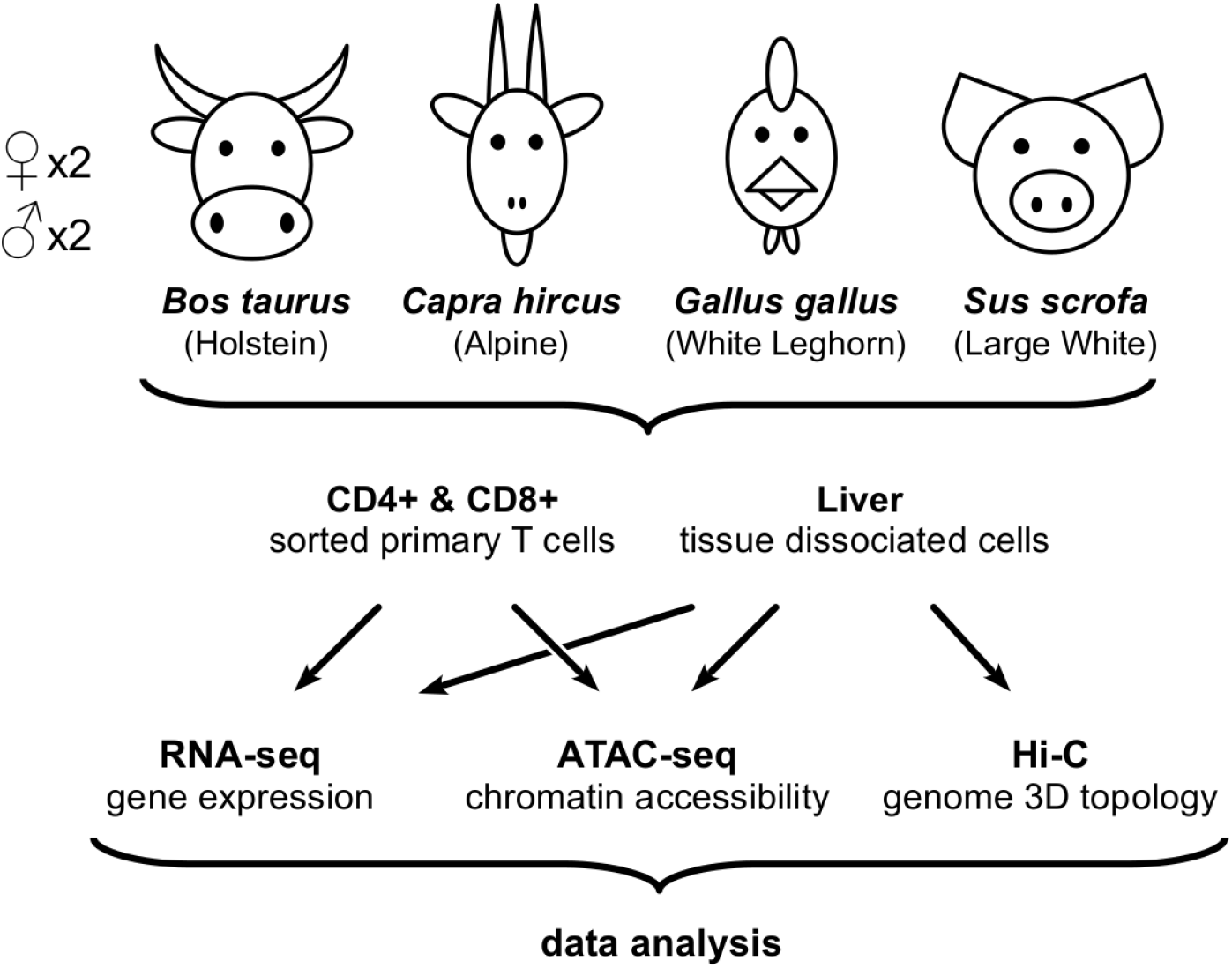
Experimental design overview. For each species, samples from liver and T cells of two males and two females were processed by RNA-seq, ATAC-seq and Hi-C assays. See Table S1 for a complete list of experiments performed and available datasets.

## Results and discussion

### High-depth RNA-seq assays provide gene expression profiles in liver and immune cells from cattle, goat, chicken and pig

For each animal (two males, two females) of the four species, we used RNA-seq to profile the transcriptome of liver, CD4+ and CD8+ T cells (see Methods, Figure 1 and Table S1). We prepared stranded libraries from polyA+ selected RNAs longer than 200bp, and we sequenced them on an Illumina HiSeq3000 (see Methods). Between 250M (chicken) and 515M (goat) read pairs were obtained per tissue on average, of which 94% (chicken) to 98% (pig) mapped to their respective genome using STAR [18, 19] (see Methods, Figure S1 and Tables S2-S4). As an initial quality control step, we processed the mapped reads with RSEM [20] to estimate the expression levels of all genes and transcripts from the Ensembl reference annotation (hereafter called “reference” genes/transcripts/annotation) (Table S2, Figure S1 and supplementary data file 2). As expected, a large majority of the reads (from 62% in cattle to 72% in goat) fell in annotated exons of the reference genes (Figure S2). In spite of the specialized scope of this initial study limited to liver and immune cells, a large share of all reference genes were detected (from 58% in chicken to 65% in goat), even considering only transcript isoforms with an expression level higher than 0.1 TPM in at least two samples (see Methods and Table 1).

**Table 1.**
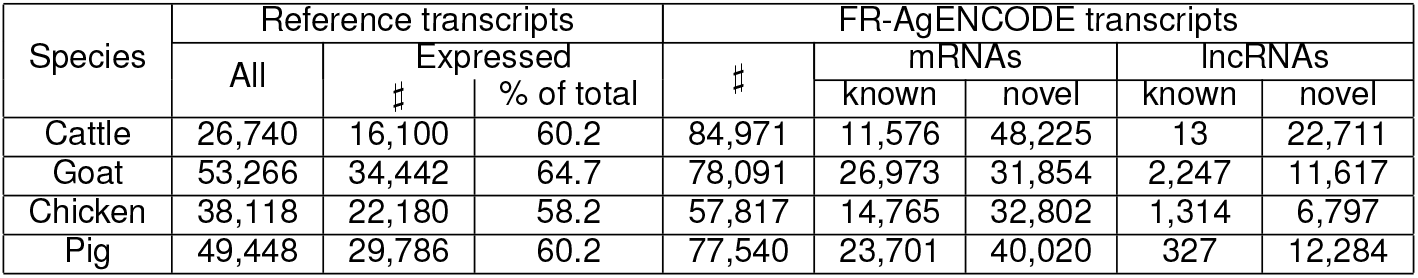
Reference and FR-AgENCODE detected transcripts. This table provides the total number of reference transcripts for each species, the number and percent of those that were detected by RNA-seq (TPM ≥ 0.1 in at least 2 samples), the total number of FR-AgENCODE transcripts, and the subsets of them that were mRNAs (known and novel) and lncRNAs (known and novel). Overall, the transcript repertoire is increased by about 100% in all species but cattle, where it is more than tripled.

| Species | Reference transcripts |  |  | FR-AgENCODE transcripts |  |  |  |  |
| --- | --- | --- | --- | --- | --- | --- | --- | --- |
|  | All | Expressed |  | # | mRNAs |  | lncRNAs |  |
|  |  | # | % of total |  | known | novel | known | novel |
| Cattle | 26,740 | 16,100 | 60.2 | 84,971 | 11,576 | 48,225 | 13 | 22,711 |
| Goat | 53,266 | 34,442 | 64.7 | 78,091 | 26,973 | 31,854 | 2,247 | 11,617 |
| Chicken | 38,118 | 22,180 | 58.2 | 57,817 | 14,765 | 32,802 | 1,314 | 6,797 |
| Pig | 49,448 | 29,786 | 60.2 | 77,540 | 23,701 | 40,020 | 327 | 12,284 |

For each species, we explored the similarity among samples using the expression profiles of the reference genes. Principal component analysis (PCA) revealed quite consistent patterns across species, where the first principal component (explaining 84 to 91% of the variability among samples) clearly separated samples according to their tissue of origin (liver vs. T cells). A more moderate yet systematic separation was observed between CD4+ and CD8+ T cells on the second principal component (Figure S3). The consistency of these patterns across species supports the reliability of our RNA-seq data.

To compare the expression pattern of reference genes between species, we first checked that the male-to-female expression ratio was globally uniform genome-wide with the exception of the sex chromosomes in chicken (Figure S4). Dosage compensation (leading for instance to X chromosome inactivation in mammals) can indeed be observed in all species except chicken, which is consistent with previous reports on dosage compensation [21]. Next, we hierarchically clustered our samples using the expression of 9,461 genes found to be orthologous across the four species (Figure 2, Methods and supplementary data file 3). Regardless of the species, liver and T cell samples clustered separately. For T cells, samples clustered first by species and then by subtypes (i.e. CD4+ versus CD8+). For most species and tissues, samples also clustered by sex, possibly due in part to the physiological state of females in lactation or laying eggs. These results confirm the higher conservation of the liver compared to the immune system gene expression program across vertebrates, and highlight the fact that when RNA-seq is performed on multiple tissues and species, samples could primarily cluster by either factor [22, 23].

**Figure 2.**
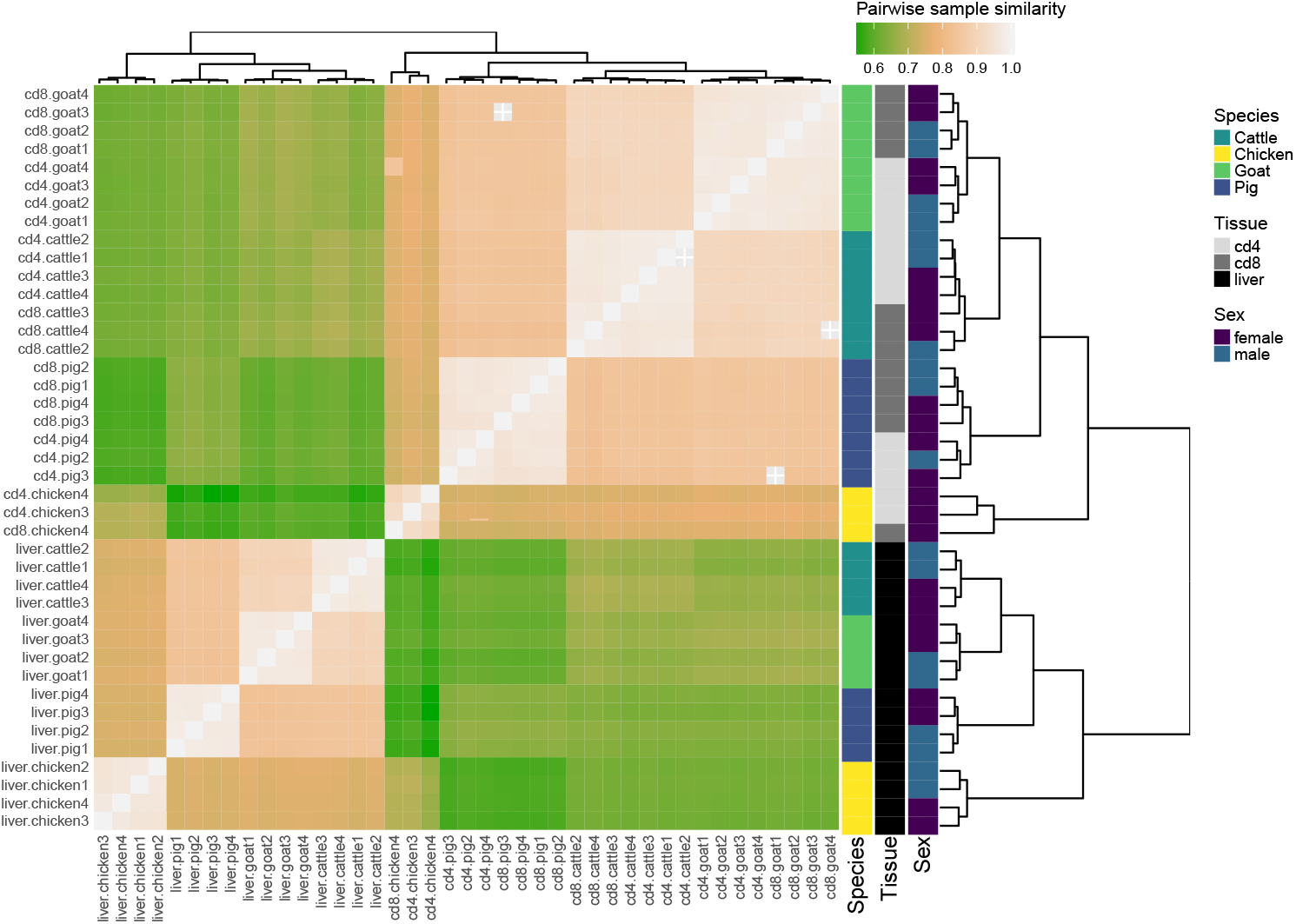
RNA-seq sample heatmap and hierarchical clustering based on the expression of the 9,461 orthologous genes across the four species. Pairwise similarity between samples is computed as the Pearson correlation between the base 10 logarithm of the expression (TPM) of the 9,461 orthologous genes. These similarities are plotted as a heatmap, where samples appear both as rows and columns and are labelled by their species, tissue and the sex of the animal. The color of each heatmap cell also reflects the similarity (Pearson correlation) between each sample pair (the lighter, the higher). Hierarchical clustering is performed using one minus the squared Pearson correlation as a distance and the complete linkage aggregation method.

### Most reference genes are differentially expressed between liver and T cells

To provide functional evidence supporting our RNA-seq data, we performed a differential gene expression analysis across tissues per species for each gene in the reference annotation. The number of exonic reads was first counted for each gene and was then TMM-normalized [24] (see Methods). Taking into account the specificities of our experimental design, in which samples from different tissues come from the same animal, we fitted generalized linear models (GLM) to identify genes with differential expression in either all pairwise comparisons between liver, CD4+ and CD8+ (model 1), or liver versus T cells globally (model 2).

As expected, the liver to T cell comparison yielded the largest number of differentially expressed genes, and relatively few genes were found to be differentially expressed between the two T cell populations (CD4+ and CD8+, see Table S5 and supplementary data file 4). Strikingly, most genes showed significantly different expression between liver and immune cells in each species (from 7,000 genes in chicken to 10,500 genes in goat), reflecting the difference between the physiological functions of these highly specialized cell types in line with findings from the GTEx project [25]. Accordingly, Gene Ontology (GO) analysis provided results in line with the role of liver in metabolism and of T cells in immunity for all species (Figures S5-8 and Methods).

In accordance with the results of the hierarchical clustering (Figure 2), most orthologous genes found to be differentially expressed between CD4+ and CD8+ T cells within species showed high variability of expression levels across species (not shown). This variability is likely caused by the natural heterogeneity in the relative proportions of T cell subtypes among the different species, as already reported between mammals [26, 27]. Nevertheless, 39 orthologous genes could consistently differentiate CD4+ and CD8+ in the four species, including mammals and chicken, which is significantly more than expected by chance (p-value < 10^−3^, permutation test). Among those, 10 and 29 genes showed significant overexpression in CD4+ and CD8+ cells respectively (Table S6). We searched for the human orthologs of theses genes in the baseline expression dataset of human CD4+ and CD8+ *αβ* T cell subsets generated by the Blueprint Epigenome Project [28, 29] and considered their relative enrichment in each cell subset. With one exception (*ACVRL1*), all genes were found to be expressed in human CD4+ and/or CD8+ *αβ* T cells and 25 of them showed a relative enrichment in CD4+ (or CD8+) human cells consistent with our data across the four species. Out of these 25 genes, six and eight genes, respectively, could be associated with CD4+ and CD8+ T cell differentiation, activation and function according to the literature (Table S6).

### Analysis of new transcripts improves and extends gene structure annotation

In order to test if our data could improve the reference gene annotation for each species, we used STAR and Cufflinks to identify all transcripts present in our samples and predict their exon-intron structures. We then quantified their expression in each sample using STAR and RSEM (see Methods and Figure S1) and only retained the transcripts and corresponding genes expressed in at least two samples with TPM ≥ 0.1. We identified between 58,000 and 85,000 transcripts depending on the species (Table 1, supplementary data file 5), hereafter called “FR-AgENCODE transcripts”.

To characterize these FR-AgENCODE transcripts with respect to the reference transcripts, we grouped them into four positional classes (see Methods): (1) *known*: a FR-AgENCODE transcript whose exon-intron structure is strictly identical to a reference transcript *(i.e.* exact same exons and introns); (2) *extension*: same as (1), but the FR-AgENCODE transcript extends a reference transcript by at least 1 bp on at least one side; (3) *alternative*: a FR-AgENCODE transcript that shares at least one intron with a reference transcript but does not belong to the previous categories (only multi-exonic transcripts can be in this class); and (4) *novel*: a FR-AgENCODE transcript that does not belong to any of the above categories. We found that most FR-AgENCODE transcripts (between 37% for goat and 49% for chicken) were of the alternative class, therefore enriching the reference annotation with new splice variants of known genes (Table S7). The proportion of completely novel transcripts was relatively high for cattle, which is likely due to the incompleteness of the UMD3.1 version of the Ensembl annotation used at the time of the study (Figure S9, Tables 1, S2 and S7).

In order to identify interesting new transcripts involved in immunity and metabolism, we first selected the novel FR-AgENCODE coding transcripts that unambiguously project to a single human coding gene. We identified 93 (cattle), 52 (goat), 74 (chicken), 26 (pig) genes, of which 12 are common to at least 2 livestock species (see Methods, Table S8, Figure S10A and supplementary data file 6). Gene set enrichment analyses on these gene lists confirmed their relevance for T cell biology (Figure S10B) and the added value of the FR-AgENCODE novel transcripts in terms of annotation of complex but important loci like TRBV and TRAV (Figure S10C).

In addition, we performed a differential gene expression analysis similar to the one done on reference genes (see above and Methods). Results were globally similar, with more than 88% of correspondence between the differentially expressed genes from the reference and the FR-AgENCODE annotation (Figure S11, Table S5, S9, supplementary data file 7). Among the latter, between 202 (chicken) and 1,032 (goat) genes were coding (at least one coding transcript predicted by FEELnc — see below) and did not overlap any reference gene on the same strand. This highlights the potential to identify novel interesting candidates for further functional characterization.

### Identification, classification and comparative analysis of lncRNAs

Since deep RNA-seq libraries allow the detection of weakly expressed transcripts [30], we sought to identify the proportion of long non-coding RNAs (lncRNAs) among the FR-AgENCODE transcripts. Using the FEELnc classifier [31] trained on the reference annotation (see Methods), we identified from 7,502 (chicken) to 22,724 (cattle) lncRNA transcripts, among which a large majority were not previously reported (Tables S10-11 and supplementary data file 8). The high number of lncRNAs found in cattle is likely due in part to the incomplete genome annotation and genome assembly used at the time of the study (Table S2). Consistent with previous reports in several species including human [32], dog [31], and chicken [33], predicted lncRNA genes had lower expression levels than reference protein-coding genes (Figure S12). The structural features of these predicted lncRNA transcripts were consistent between the four species: lncRNAs are spliced but with fewer exons (1.5 vs. 10) and higher median exon length (660 vs. 130bp) compared to mRNAs (Figure S12). LncRNAs are also smaller than mRNAs (1800 vs. 3600bp). Notably, the lower number of exons and consequent smaller size of lncRNAs compared to mRNAs could also be due to the weaker expression of lncRNAs, which makes their structure more difficult to identify [34].

In addition to the coding/non-coding classification, FEELnc can also categorize lncRNAs as intergenic or intragenic based on their genomic positions with respect to a provided set of reference genes (usually protein coding), and considering their transcription orientation with respect to these reference genes. This analysis revealed an overwhelming majority of intergenic lncRNA genes over intragenic ones (Table S10), which is consistent with results obtained in human [32] and in chicken [33].

We and others previously showed a sharp decrease in lncRNA sequence conservation with increasing phylogenetic distance [32, 33, 35], in particular between chicken and human that diverged 300M years ago. We therefore analyzed lncRNA conservation between the four livestock species using a synteny approach based on the orthology of protein-coding genes surrounding the lncRNA and not on the lncRNA sequence conservation itself [33] (see Methods). We found 73 such conserved, or syntenic, lncRNAs across cattle, goat and pig, 19 across cattle, chicken and pig, and 6 across all four species (supplementary data file 8). All were expressed in these species and located in the same orientation with respect to the flanking orthologous genes. An example of such a conserved lncRNA, hitherto unknown in our four species, is provided in Figure 3. In human, this lncRNA is called *CREMos* for *“CREM* opposite sense” since it is in a divergent position with respect to the neighboring *CREM* protein-coding gene. Interestingly, synteny is conserved across species from fishes to mammals and the *CREMos* lncRNA is overexpressed in T cells while the *CREM* protein-coding gene is overexpressed in liver in goat, cattle and chicken (Figure 3). Additional examples of syntenic lncRNAs are provided in supplementary data file 8, and the ones found to be conserved between the 4 species are represented in Figure S13.

**Figure 3.**
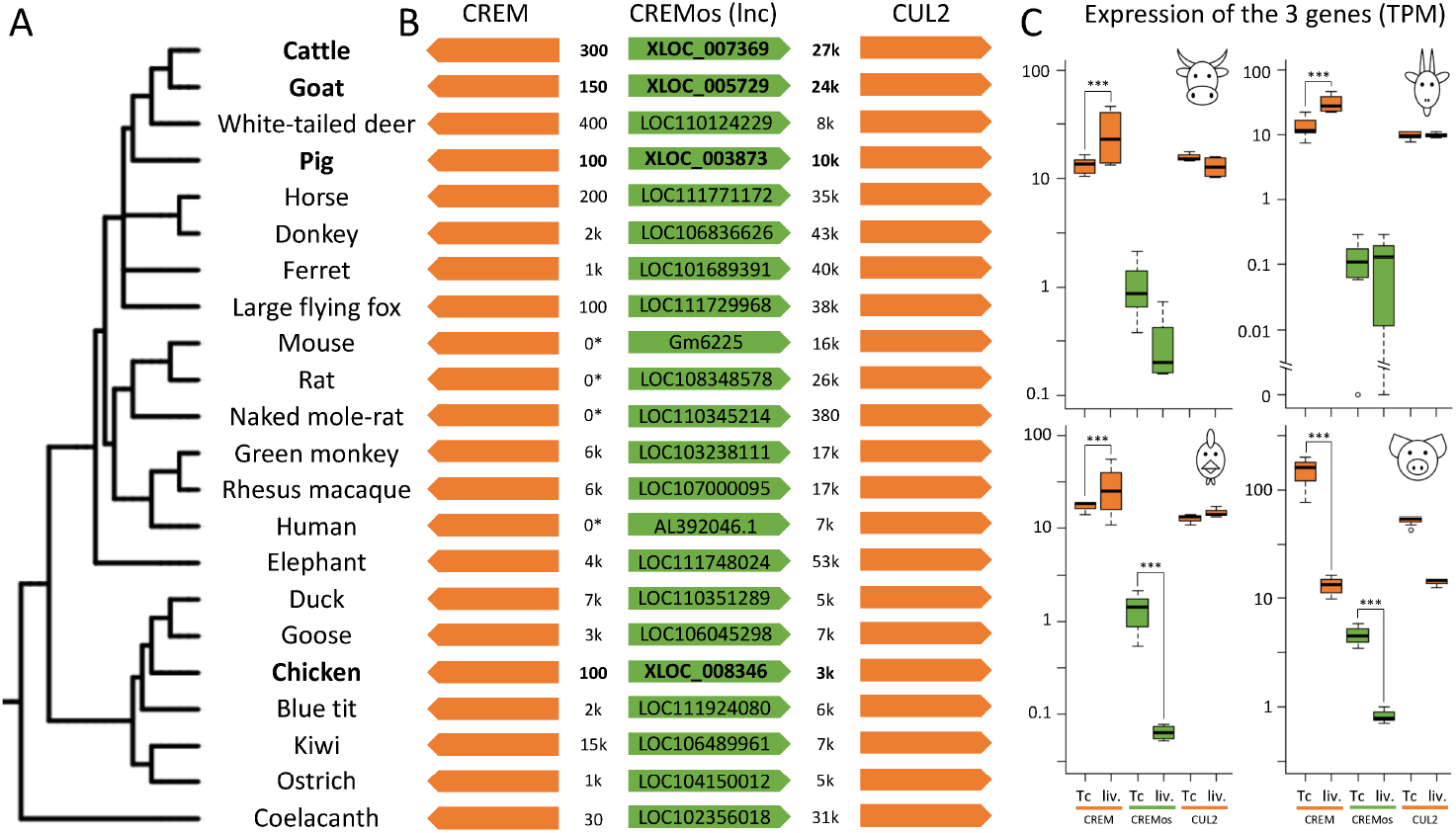
A novel lncRNA conserved across multiple species. Phylogenetic representation based on the NCRI taxonomy of the 22 annotated species from fishes to mammals using iTOL [75] **(A,B)** Three gene syntenic region centered on *CREMos* lncRNA with NCBI IDs for lncRNAs already annotated in reference databases and distance between entities in nucleotides; The cases where *CREM* and *CREMos* genes are overlapping are indicated by the “0*” distance. **(C)** Expression of the 3 genes in cattle, goat, chicken and pig: *CREMos* is generally less expressed in liver than in T cells (in cattle, chicken and pig) whereas *CREM* is generally more expressed in liver than in T cells (in cattle, chicken and goat).

### Landscape of chromatin accessibility in cattle, goat, chicken and pig

We used ATAC-seq to profile the accessible chromatin of liver, CD4+ and CD8+ T cells in animals from the four species. Between 480M (chicken) and 950M (pig) ATAC-seq fragments were sequenced per species, and were processed by a standard pipeline (Methods and Figure S14). Peaks were called in each tissue separately (see Methods), resulting in between 26,000 (pig, liver) and 111,000 (pig, cd8) peaks per tissue (Table S12). Those peaks were further merged into a unique set of peaks per species, resulting in between 75,000 (goat) and 149,000 (pig) peaks (Table S12 and supplementary data file 9), covering 1 to 5% of the genome. The average peak size was around 600 bp for all species, except for chicken where it was less than 500 bp. Merging tissue peaks did not result in much wider peaks (Figure S15).

In comparison to the reference annotation, about 10-15% of the peaks lie at 5Kb or less from a Transcription Start Site (TSS) and can be considered to be promoter peaks. The precise distribution of these promoter peaks showed a clear higher accumulation at the TSS for all species (Figure 4), supporting the quality of both the annotation and our data. Importantly, this signal was also observed around the TSS of novel FR-AgENCODE transcripts (i.e. those not from the known class; Figure S16).

**Figure 4.**
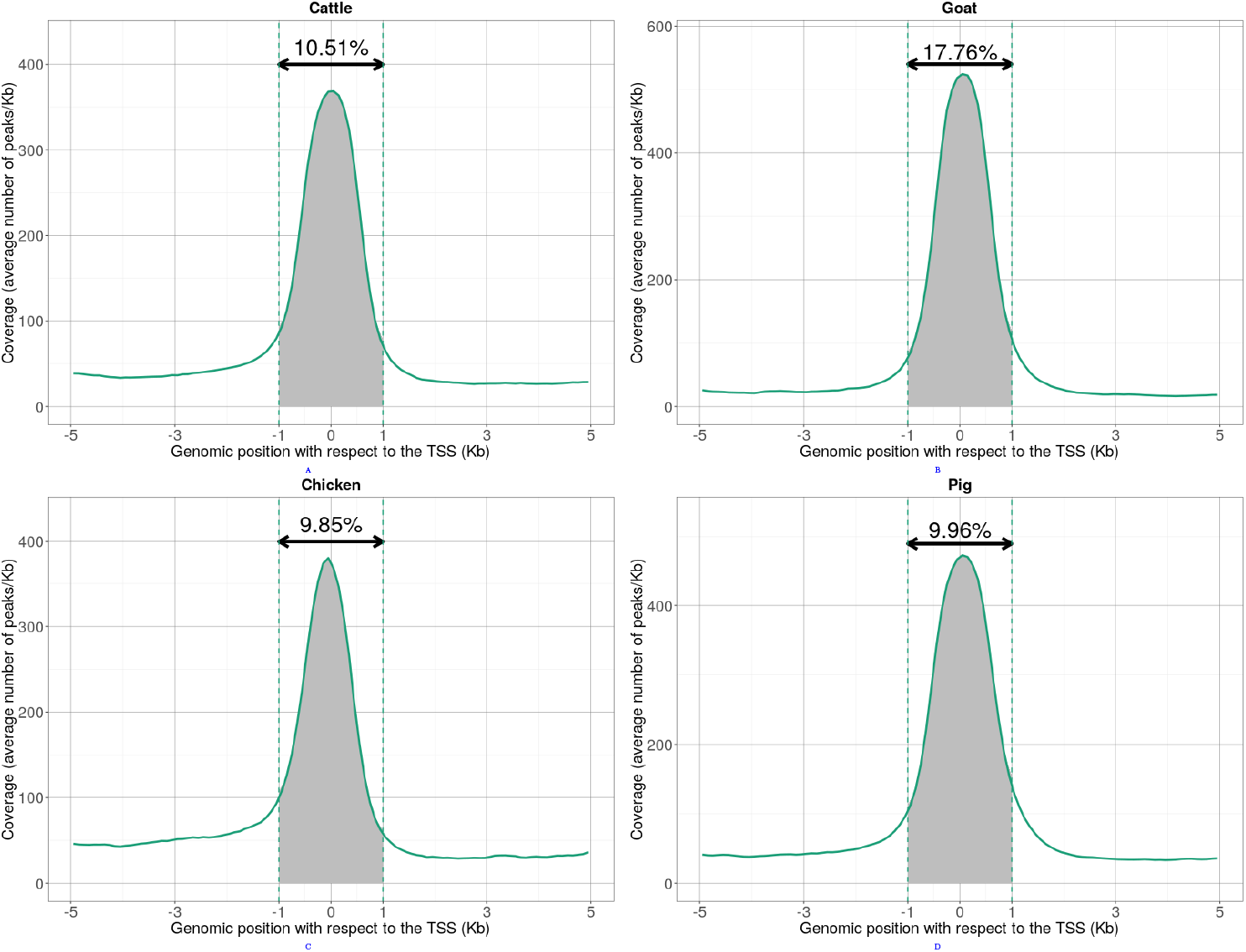
Density of ATAC-seq peaks around Transcription Start Sites (TSS) for cattle (A), goat (B), chicken (C) and pig (D). Mean coverage values of ATAC-seq peaks (*y*-axis) were computed genome-wide relatively to TSS positions (*x*-axis). The proportion of ATAC-seq peaks within the [−1; +1]Kb interval is represented by the shaded area between the dotted lines. The corresponding percentage is indicated above the double arrow, indicating that most of the ATAC-seq signal is distal to TSSs.

The vast majority of the peaks, however, were either intronic or intergenic (Figure 4, Figure S17, supplementary data file 9), similar to GWAS variants associated with human diseases [3]. In particular, from 38% (goat) to 55% (cattle) of the peaks were located at least 5Kb away from any reference gene (Figure S17), indicating that ATAC-seq can detect both proximal and distal regulatory regions.

Since active enhancers are expected to be enriched in chromatin regions that are both accessible and tagged with specific histone modification marks, we compared our ATAC-seq peaks to histone ChIP-seq peaks from another functional genomics study [36]. In that study, two histone modification marks (H3K4me3 and H3K27ac) were profiled in the genome of 20 mammals including pig, for which we have ATAC-seq data in the same tissue (liver). Although only a small proportion of the ChIP-seq peaks overlapped our liver ATAC-seq peaks (2.8% of the 8,538 H3K4me3 peaks and 4.1% of the 25,502 H3K27ac peaks), this proportion was significantly higher than expected by chance as measured by shuffling peak positions (p-value = 0.017 and p-value = 0.026 respectively, permutation test). Interestingly, this subset of overlapping peaks have significantly higher q-value signal scores than their nonoverlapping counterparts (p-value = 8 × 10^−6^, p-value = 1.6 × 10^−5^ and p-value < 2.2 × 10^−16^ for ATAC-seq, H3K4me3 and H3K27ac scores respectively, Wilcoxon tests), which confirms the existence of a common signal between the datasets.

To further characterize functional regulatory sites in our samples, we compared chromatin accessibility between liver and T cells. The ATAC-seq peaks of each species were quantified in each sample and resulting read counts were normalized using a loess correction (see Methods). A differential analysis similar to the one used for RNA-seq genes was then performed on normalized counts (see Methods). We identified from 4,800 (goat) to 13,600 (chicken) differentially accessible (DA) peaks between T cells and liver (Table S13 and supplementary data file 10). To test for the presence of regulatory signals in these regions, we computed the density of transcription factor binding sites (TFBS) in ATAC-seq peaks genome-wide. Interestingly, TFBS density was significantly higher in DA ATAC-seq peaks compared to non-DA ATAC-seq peaks (Model 2; p-value < 7.1 × 10^−4^ for goat and p-value < 2.2 × 10^−16^ for chicken and pig, Wilcoxon tests, see Methods). This enrichment was also observed for distal ATAC-seq peaks, at least 5kb away from promoters (not shown), and suggests that differentially accessible peaks are more likely to have a regulatory role than globally accessible peaks.

### Promoter accessibility is associated with both positive and negative regulation of gene expression

Accessible promoters are commonly associated with gene activation [37, 38]. Given the specific distribution of the ATAC-seq signal, we initially focused on proximal chromatin peaks (i.e. at 1Kb or less from a gene TSS) and used them to assign a promoter accessibility value to each gene. Using normalized read counts (see Methods), we investigated the correlation between ATAC-seq and RNA-seq data either across all genes within each sample, or across all samples for each gene.

Within each sample, genes with highly accessible promoters showed higher expression values globally (Figure S18), as already reported in mouse and human [39]. For pig and goat, the number of available samples further allowed us to compute for each gene the correlation between promoter accessibility and gene expression across all samples (Figure S19 and Methods). Interestingly, while the correlation value distribution appeared to be unimodal for non-differentially expressed genes, it was bimodal for differentially expressed genes, with an accumulation of both positive and negative correlation values (Figures 5 and S19). This pattern supports the existence of different types of molecular mechanisms (i.e. both positive and negative) involved in gene expression regulation.

**Figure 5.**
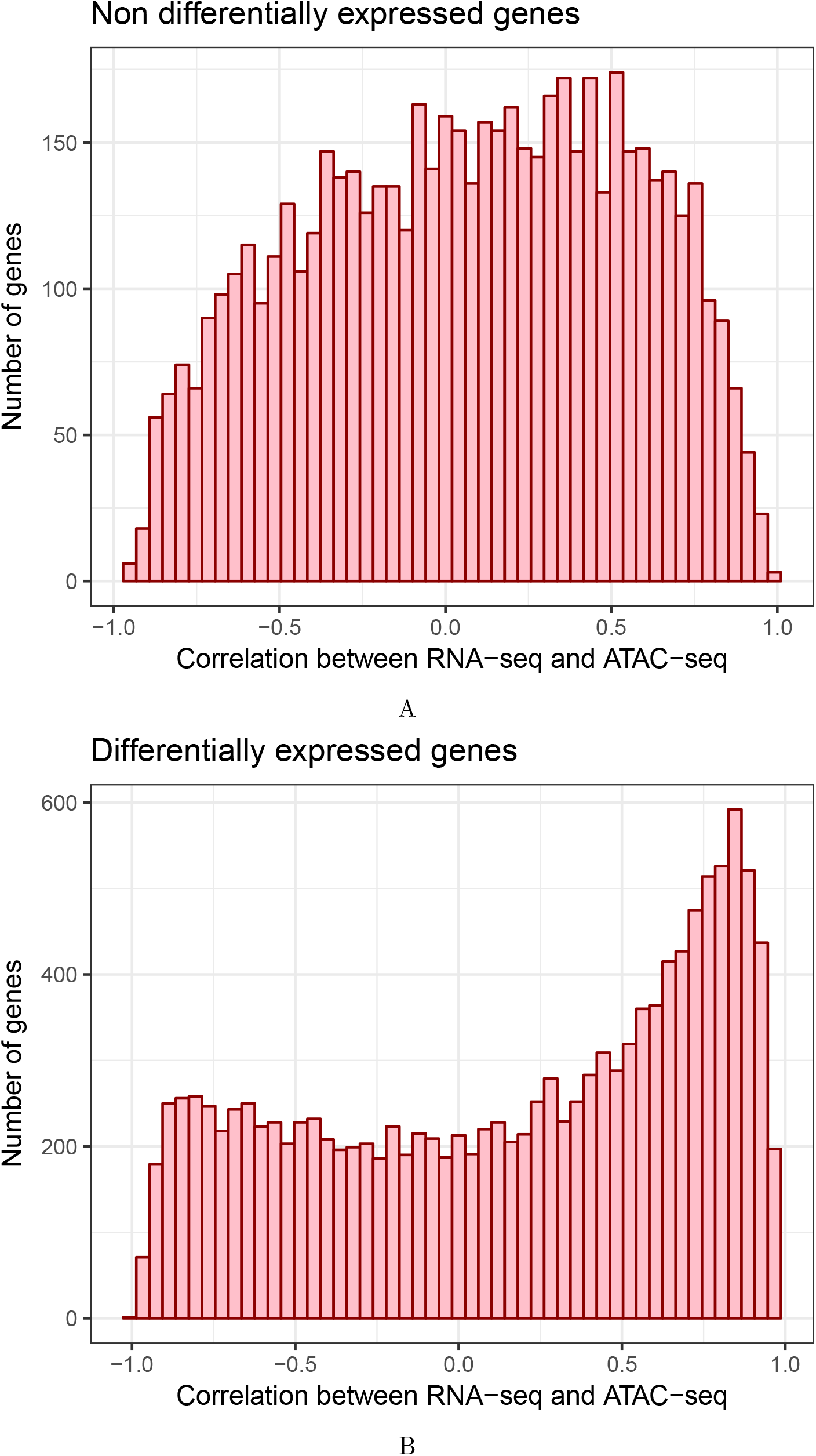
Correlation between gene expression and promoter accessibility in pig. For each expressed FR-AgENCODE gene with an ATAC-seq peak in the promoter region, the Pearson correlation was computed between the base 10 logarithm of the RNA-seq gene expression (TMM) and the base 10 logarithm of the ATAC-seq chromatin accessibility (normalized by csaw). The distribution is represented for genes with no significant differential expression between liver and T cells **(A, top)** and for differentially expressed genes **(B, bottom)**. The distribution obtained for differentially expressed genes showed an accumulation of both positive and negative correlation values, suggesting a mixture of regulatory mechanisms.

### Comparative genomics reveals a core set of conserved chromatin accessible sites

We then investigated the evolution of chromatin accessibility genome-wide by performing a comparative analysis of all (proximal and distal) conserved ATAC-seq peaks across species. We identified conserved peaks by aligning all the sequences that correspond to peaks from each species (both proximal and distal) to the human genome (see Methods). Most peaks could be mapped globally, with an expected strong difference between mammals (72%-80% of the peaks) and chicken (12% of the peaks). After keeping the best sequence hits, merging them on the human genome and retaining the unambiguous ones (see Methods), we obtained a set of 212,021 human projections of livestock accessible chromatin regions, that we call human hits.

A large majority of the human hits (about 86%) originated from a single livestock species, which is consistent with previous reports about the fast evolution of regulatory elements and the species-specific feature of many enhancers [5, 36]. Nevertheless, the remaining 28,292 human hits (14%) had a conserved accessibility across two or more livestock species (Methods and supplementary data file 13). As they share both sequence conservation and experimental evidence between several species, we refer to those human hits as “orthologous peaks” and to the number of species sharing the peak as its “orthology level”. Among them, 1,083 orthologous peaks had an orthology level of 4, *i.e.* shared by all 4 livestock species. Human hits from a single species were assigned an orthology level of 1. As previously done with the orthologous genes using RNA-seq data, we performed a hierarchical clustering of the samples based on the normalized accessibility values of the peaks with an orthology level of 4 (Figure S21). Contrary to what was observed from the expression data, samples here mostly clustered according to species first, with the chicken as a clear outlier. However for the two phylogenetically closest species (goat and cow), we observed that all T cells clustered together, separately from liver. This suggests a stronger divergence and specialization of the regulatory mechanisms compared to the gene expression programs.

In addition, shuffling the peak positions within each species did not drastically change the mapping efficiency on the human genome overall but resulted in a much lower proportion of orthologous peaks (from 14% to 3% human hits with an orthology level > 1, see Methods). Also, the overlap on the human genome between all the 212,021 human hits and ENCODE DNAse I hypersensitive sites from liver and T cell samples [40] was three to four times higher than with the random set of shuffled peaks (25-39% per species vs. 7-9%).

Lastly, human hits that were identified as differentially accessible between liver and T cells in at least one of the species had higher PhastCons conservation scores on the human genome than the non differential peaks of the same orthology level (Figure 6). This difference was significant for three out of the four orthology levels (p-values < 0.01 overall, Wilcoxon tests), supporting a selective pressure on functionally active regulatory regions. Remarkably, this contrast was even stronger after discarding human hits close to a TSS in any of the species (Figure S22, p-values < 10^−6^ overall, Wilcoxon tests, supplementary data file 12), in line with a specific conservation of distal regulatory elements beyond the promoter regions. Altogether, these results highlight a core set of conserved regulatory regions from birds to mammals that include both proximal and distal sites.

**Figure 6.**
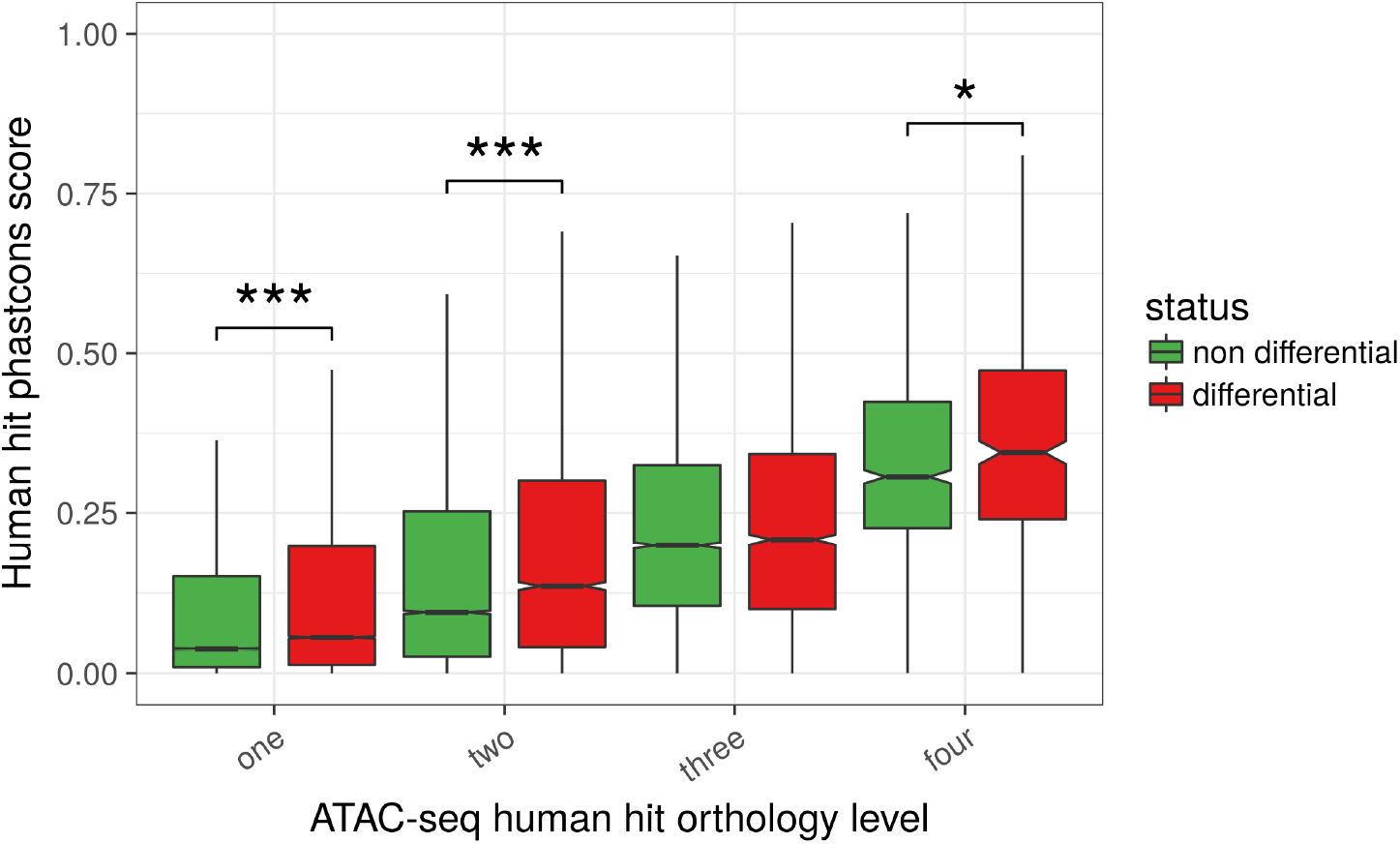
Relationship between chromatin accessibility conservation and differential accessibility. Phastcons scores of ATAC-seq human hits were plotted after dividing the human hits according to both their orthology level (between 1 and 4, *x*-axis) and their differential accessibility (DA) status (DA in at least one species or DA in none of the 4 species, boxplot color). Although the phastcons score obviously increases with the orthology level, it is clear that, for a given orthology level, the phastcons score is higher for DA human hits than for non DA human hits (all orthology levels except 3, p-values < 0.01 overall, Wilcoxon tests) (number of elements in the boxplots from left to right: 163509, 21578, 16329, 4437, 6231, 6231, 2241, 878, 417).

### Comprehensive maps of topological domains and genomic compartments in goat, chicken and pig

In order to profile the structural organization of the genome in the nucleus, we performed *in situ* Hi-C on liver cells from the two male and the two female samples of pig, goat and chicken. The in *situ* Hi-C protocol was applied as previously described [41] with slight modifications (see FAANG protocols online and Methods). Reads were processed using a bioinformatics pipeline based on HiC-Pro [42] (Methods). From 83 to 91% of the reads could be genomically mapped depending on the sample, and after filtering out all inconsistent mapping configurations we obtained a total of 182, 262 and 290M valid read pairs in goat, chicken and pig respectively (Table S14 and Figure S23). These sequencing depths allowed us to build interaction matrices (or Hi-C contact heatmaps) at the resolution of 40 and 500Kb in order to detect Topologically Associating Domains (TADs) and A/B compartments respectively (Figures S24-25).

We identified from ≈ 5,400 (chicken) to 11,000 (pig) TADs of variable sizes (150 to 220Kb on average, Table S15 and supplementary data file 12), with a 90-92% genome-wide coverage. To validate these domains predicted by armatus [43] (see Methods), we computed three metrics along the genome: the Directionality Index (DI), to quantify the degree of upstream or downstream interaction bias for any genomic region [44], the local interaction score to represent the insulation profile along the genome [45, 46], and the density of in *silico* predicted CTCF binding sites, expected to be prevalent at TAD boundaries [47, 48]. For each species, we observed that the distribution of these three metrics was consistent with previous reports on model organisms (Figure 7 and Figure S26), supporting the relevance of our topological annotation.

**Figure 7.**
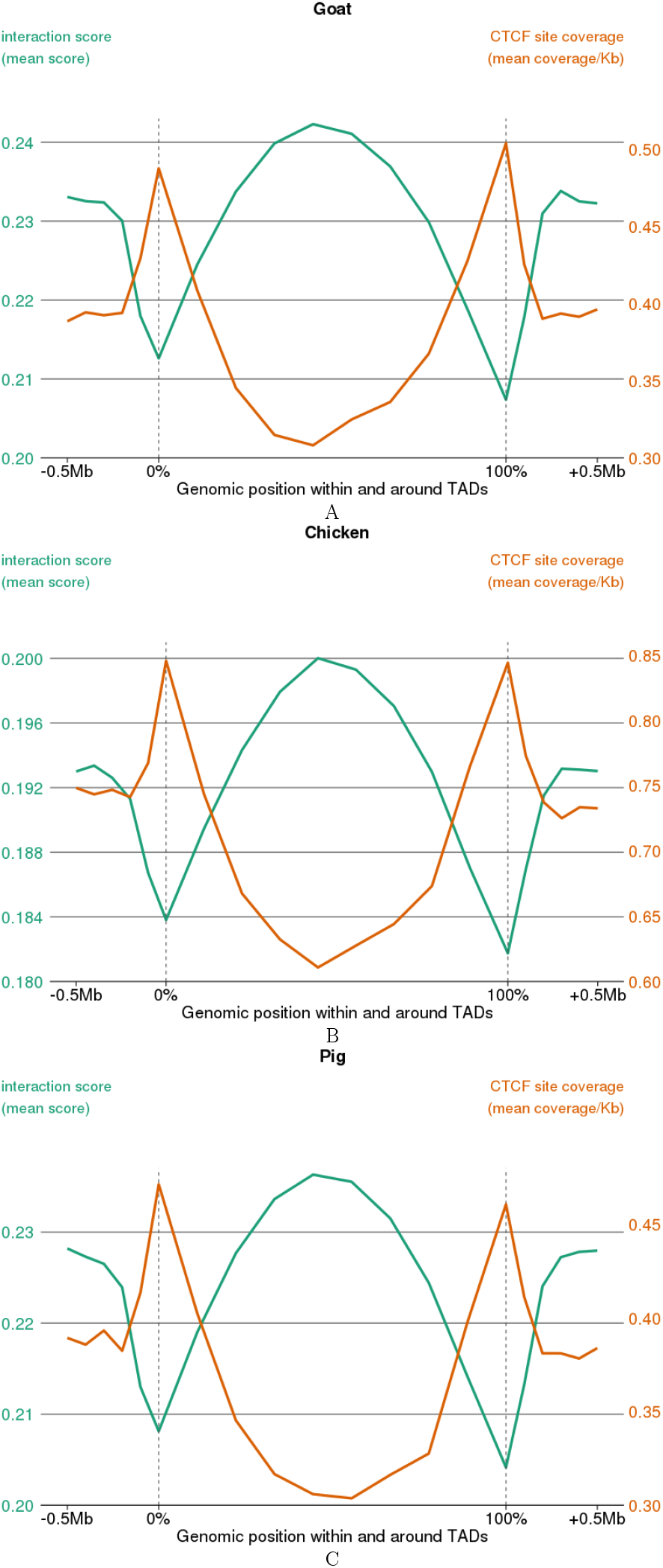
CTCF motif density and local interaction score within and around TADs. Local interaction score across any position measured from Hi-C matrices and represented on the *y*-axis (left). The mean density of predicted CTCF binding sites is also shown on the *y*-axis (right). Mean interaction score and CTCF density are plotted relative to the positions of Hi-C-derived Topologically Associating Domains. Dotted lines indicate TAD boundaries. Absolute scale is used on the *x*-axis up to 0.5Mb on each side of the TADs while relative positions are used inside the domains (from 0 to 100% of the TAD length).

At a higher organizational level, we identified “active” (A) and “inactive” (B) epigenetic compartments as defined by [17] (see Methods and Figure S24). We obtained from ≈ 580 to 700 compartments per genome with a mean size between 1.6Mb (chicken) and 3.4Mb (goat) and covering between 71.9% (goat) and 88.6% (pig) of the genome (see Table S15 and supplementary data file 11). We also observed high consistency of the compartments between replicates (same compartment for 80% of the loci in all 4 animals, see Figure S27). In model organisms, A compartments represent genomic regions enriched for open chromatin and transcription compared to B compartments [44]. By using RNA-seq and ATAC-seq data from the same liver samples as those for which we had Hi-C data, we observed that both the average gene expression and the average chromatin accessibility were significantly higher in A than in B compartments (Figure 8, p-value < 2.2 × 10^−16^ for each comparison, Wilcoxon tests), emphasizing the biological consistency of our results across all molecular assays and species.

**Figure 8.**
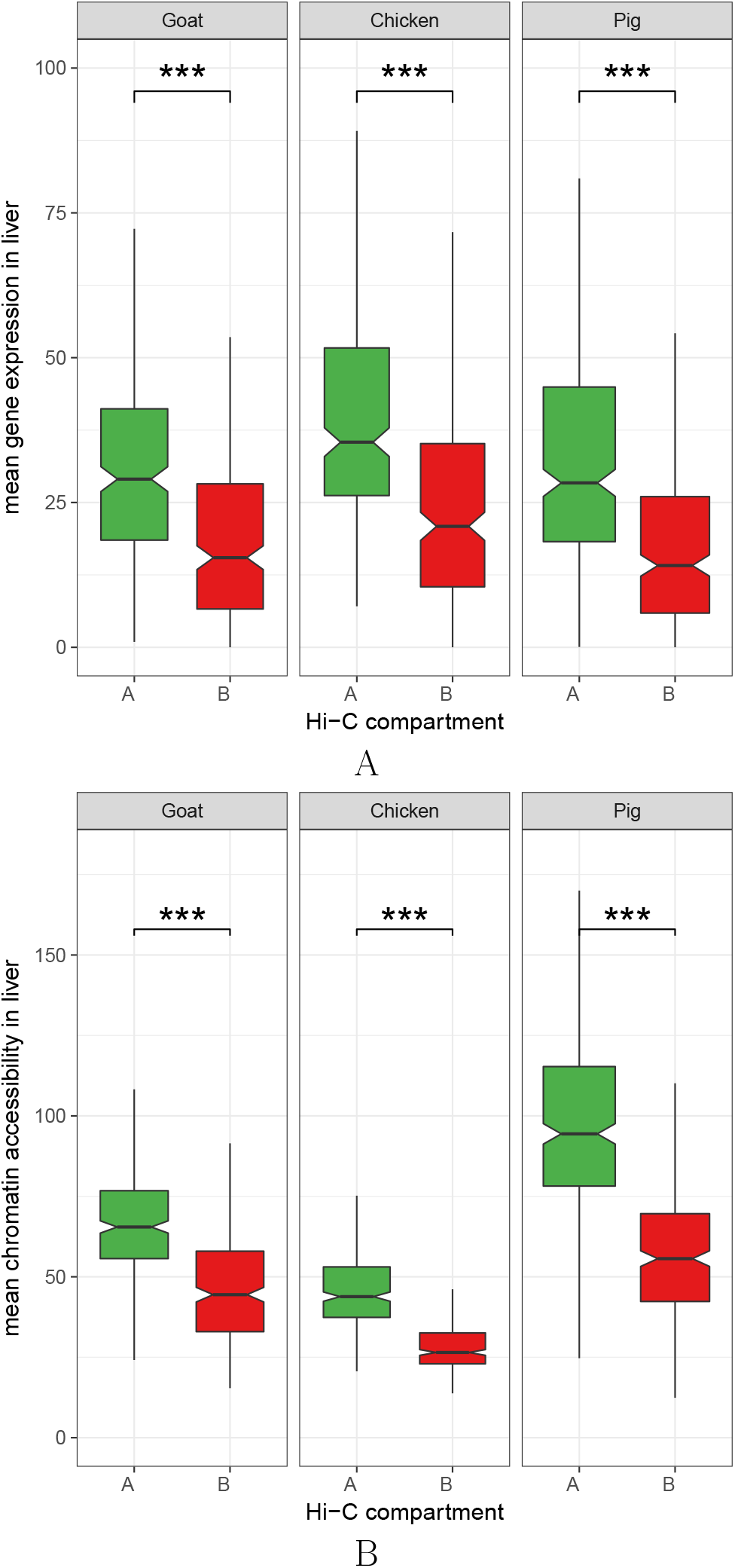
Gene expression and chromatin accessibility in A and B topological compartments. For the three species with Hi-C-derived A and B compartments, the distribution of the RNA-seq gene expression values (normalized read counts, top panel) and ATAC-seq chromatin accessibility values (normalized read counts, bottom panel) is shown per compartment type. A: “active” compartments. B: “repressed” compartments. As Hi-C data was only available from liver, only RNA-seq and ATAC-seq values from the same samples were considered. The significant and systematic difference of gene expression and chromatin accessibility values between A and B compartments (p-values < 2.2 × 10^−16^ overall, Wilcoxon tests) confirms a general consistency between RNA-seq, ATAC-seq and Hi-C data across species.

### Genome structure comparison reveals a multi-scale selective pressure on topological features across evolution

It has been shown that the general organization in TADs tends to be conserved across species [48, 49] and that the presence of specific TAD boundaries can be crucial for biological functions like development [50]. In line with these reports, we wondered if TAD boundaries might play a fine grain regulatory role beyond a binary model of simple absence/presence. Under this assumption, we hypothesized that the insulating capacity of conserved TAD boundaries could be under selective pressure. We therefore assessed the link between their insulation potential and their evolutionary conservation.

As previously done with ATAC-seq peaks (see Methods and above) we first mapped all the TAD boundaries from each species to the human genome to identify the orthologous ones. Pairwise comparisons of their local interaction scores showed a clear correlation between our species (Figure 9A). Since the interaction score here reflects the proportion of cis-contacts across a TAD boundary, such a correlation supports a conservation of the insulation strength between adjacent TADs. Strikingly, similar correlations were obtained between each of our mammalian species and human (GM12878 cell line, see Methods and Figure 9B-C) [41]. Beside confirming a general conservation of the TAD structure throughout evolution, these results emphasize the quantitative nature of this activity, in line with previous findings [48]. Moreover, a conserved insulation level at TAD boundaries suggests various degrees of functional impact and a fine control of their regulatory role, involving complex molecular mechanisms.

To further characterize this link between conservation and TAD strength, we assigned to each boundary an orthology level depending on the number of livestock species with a common hit on the human genome, as we did for ATAC-seq peaks (see above and supplementary data file 13). In all 3 species, we observed that the higher the orthology level of TAD boundaries, the lower the interaction score (Figure 9D). These results revealed that TAD boundaries under stronger selective pressure had higher insulation activities and, expectedly, a more important role in terms of genome architecture and regulatory function.

Unlike TADs, chromosomal A/B compartments have often been reported as highly variable between tissues or developmental stages, involving dynamic mechanisms of epigenetic control [51–53]. Here, we postulated that despite its plasticity, the structural organization in genome compartments for a given tissue could also be under selective pressure across a large phylogenetic spectrum. As active compartments are known to be gene-rich we first confirmed that, although both compartment types roughly have the same size, most genes were found in A compartments in each species (Figure S27). In addition, we observed that the general proportion of genes in A compartments was remarkably stable across species (66.9%, 69.7% and 70.1% of all genes in chicken, goat and pig respectively). The 5,728 orthologous genes with a predicted compartment in all three species were also found to be preferentially localized in active compartments, with slightly higher proportions than for all genes in general (69.5%, 75.9%, and 76.4% for chicken, goat, and pig respectively), probably due to the fact that conserved genes tend to have higher expression levels.

Since all these orthologous genes were assigned a compartment type (i.e. A or B label) in each species separately, we tested whether any significant conservation of compartment type across species could be detected. Among the 5,728 orthologous genes, 3,583 had the same compartment type in all species, which was 49% more than expected by chance assuming independence between species. This cross-species conservation was observed for both A and B compartments (p-value < 2.2× 10^−17^ for both, χ^2^ goodness-of-fit test), suggesting that such conservation was not restricted to regions of higher gene expression.

Altogether, results from the cross-species comparisons of ATAC-seq peaks, TAD boundaries and A/B compartments reveal a general conservation of the genome structure at different organizational levels from birds to mammals, and sheds a new light on the complex interplay between genome structure and function.

## Conclusion

We report the first multi-species and multi-assay genome annotation results obtained by a FAANG project. The main outcomes were consistent with our expectations and provide new evolutionary insights about regulatory and structural aspects of the genome:

- Despite only three different tissues being used, a majority of the reference transcripts could be detected. Moreover, the newly identified transcripts considerably enrich the reference annotations of these species.
- Differential analyses of gene expression in liver and T cells yielded results consistent with known metabolism and immunity functions and identified novel interesting candidates for functional analyses, including conserved syntenic lncRNAs.
- ATAC-seq data allowed an abundance of potential regulatory regions to be mapped, and, upon integration with RNA-seq data, suggested complex mechanisms of gene expression regulation. Comparative genomics analyses revealed evolutionary conservation both for proximal and distal regulators.
- Hi-C experiments provided the first set of genome-wide 3D interaction maps of the same tissue from three livestock species. Beyond the chromosome topology annotation, the analysis showed high consistency with gene expression and chromatin accessibility. Multi-species analyses revealed a global selective pressure on organizational features of the genome structure at different scales, beyond the TAD level.

Therefore, the FR-AgENCODE group has delivered a strong proof of concept of a successful collaborative approach at a national scale to apply FAANG guidelines to various experimental procedures and animal models. This notably includes the set up of a combination of sequencing assays on primary cells and tissue-dissociated cells, as well as a large collection of documented tissue samples available for further molecular characterization in future projects. It also confirmed, in line with several studies in model species [4–6, 8] the value of combining molecular assays on the same samples to simultaneously identify the transcriptomes and investigate underlying regulatory mechanisms.

In the context of the global domesticated animal genome annotation effort, lessons learned from this pilot project confirm conclusions drawn by the FAANG community regarding the challenges to be addressed in the future [12]. Furthermore, the mosaic nature of a global annotation effort that gathers contributions from various partners worldwide emphasizes the challenge of translating recent advances from the field of data science into efficient methods for the integrative analysis of ‘omics data and the importance of future meta-analyses of several datasets [15].

Altogether, these annotation results will be useful for future studies aiming to determine which subsets of putative regulatory elements are conserved, or diverge, across animal genomes representing different phylogenetic taxa. This will be beneficial for devising efficient annotation strategies for the genomes of emerging domesticated species.

## Methods

### Animals, sampling and tissue collections

#### Animals and breeds

Well-characterized breeds were chosen in order to obtain well-documented samples. Holstein is the most widely used breed for dairy cattle. For goats, the Alpine breed is one of the two most commonly used dairy breeds, and for pigs, the Large white breed is widely used as a dam line. For chickens, the White Leghorn breed was chosen as it provides the genetic basis for numerous experimental lines and is widely used for egg production.

Four animals were sampled for each species, two males and two females. They all had a known pedigree. Animals were sampled at an adult stage, so that they were sexually mature and had performance records, obtained in known environmental conditions. Females were either lactating or laying eggs.

All animals were fasted at least 12 hours before slaughter. No chemicals were injected before slaughtering, animals were stunned and bled in a licensed slaughter facility at the INRA research center in Nouzilly.

#### Samples

Liver samples of 0.5 cm^3^ were taken from the edge of the organ, avoiding proximity with the gallbladder and avoiding blood vessels. Time from slaughter to sampling varied from 5 minutes for chickens to 30 minutes for goats and pigs and 45 minutes for cattle. For the purpose of RNA-seq, samples were immediately snapfrozen in liquid nitrogen, stored in 2ml cryotubes and temporarily kept in dry ice until final storage at −80°C.

For mammals, whole blood was sampled into EDTA tubes before slaughter; at least one sampling took place well before slaughter (at least one month) and another just before slaughter, in order to obtain at least 50 ml of whole blood for separation of lymphocytes (PBMC). PBMC were re-suspended in a medium containing 10% FCS, counted, conditioned with 10% DMSO and stored in liquid nitrogen prior to the sorting of specific cell types: CD3+CD4+ (“CD4”) and CD3+CD8+ (“CD8”).

For chicken, spleen was sampled after exsanguination. Spleen leucocytes were purified by density-gradient separation to remove nucleated erythrocytes contamination and stored in liquid nitrogen prior to CD4+ and CD8+ T cell sorting.

All protocols for liver sampling, PBMC separation, splenocyte purification and T cell sorting can be found at ftp://ftp.faang.ebi.ac.uk/ftp/protocols/samples/

### Experimental assays and protocols

All assays were performed according to FAANG guidelines and recommendations, available at http://www.faang.org. All detailed protocols used for RNA extraction and libraries production for RNA-seq, ATAC-seq and Hi-C are available at http://ftp.faang.ebi.ac.uk/ftp/protocols/assays/.

#### RNA extraction

Cells and tissues were homogenized in TRIzol reagent (Thermo) using an ULTRA-TURRAX (IKA-Werke) and total RNAs were extracted from the aqueous phase. They were then treated with TURBO DNase (Ambion) to remove remaining genomic DNA and then processed to separate long and small RNAs using the mirVana miRNA Isolation kit. Small and long RNA quality was assessed using an Agilent 2100 Bioanalyzer and RNA 6000 nano kits (Agilent) and quantified on a Nanodrop spectrophotometer.

#### RNA-seq

Stranded mRNA libraries were prepared using the TruSeq Stranded mRNA Sample Prep Kit -V2 (Illumina) on 200 ng to 1*μ*g of total long RNA with a RNA Integrity Number (RIN) over 8 following the manufacturer’s instructions. Libraries were PCR amplified for 11 cycles and library quality was assessed using the High Sensitivity NGS Fragment Analysis Kit DNF-474 and the Fragment Analyser system (AATI). Libraries were loaded onto a High-seq 3000 (Illumina) to reach a minimum read numbers of 100M paired reads for each library.

#### Hi-C

*In situ* Hi-C libraries were made according to [41] with a few modifications. For all species, fresh liver biopsies were dissociated using Accutase, and each resulting cell suspension was filtered using a 70 μm cell strainer. Cells were then fixed with 1% formaldehyde for 10 minutes at 37°C and fixation was stopped by adding Glycine to a final concentration of 0.125M. After two washes with PBS, cells were pelleted and kept at −80°C for long term storage. Subsequently, cells were thawed on ice and 5 million cells were processed for each Hi-C library. Cell membranes were disrupted using a potter-Elvehjem PTFE pestle and nuclei were then permeabilized using 0.5% SDS with digestion overnight with HindIII endonuclease. HindIII-cut restriction sites were then end-filled in the presence of biotin-dCTP using the Klenow large fragment and were religated overnight at 4°C. Nucleus integrity was checked using DAPI labelling and fluorescence microscopy. Nuclei were then lysed and DNA was precipitated and purified using Agencourt AMPure XP beads (Beckman Coulter) and quantified using the Qubit fluorimetric quantification system (Thermo). Hi-C efficiency was controlled by PCR using specific primers for each species and, if this step was successful, DNA was used for library production. DNA was first treated with T4 DNA polymerase to remove unligated biotinylated ends and sheared by sonication using a M220 Covaris ultra-sonicator with the DNA 550pb SnapCap microtube program (Program length: 45s; Picpower 50; DutyF 20; Cycle 200; Temperature 20°C).

Sheared DNA was then size-selected using magnetic beads, and biotinylated fragments were purified using M280 Streptavidin Dynabeads (Thermo) and reagents from the Nextera_Mate_Pair Sample preparation kit (Illumina). Purified biotinylated DNA was then processed using the TrueSeq nano DNA kit (Illumina) following the manufacturer’s instructions. Libraries were amplified for 10 cycles and then purified using Agencourt AMPure XP beads. Library quality was assessed on a Fragment Analyser (AATI) and by endonuclease digestion using NheI endonuclease. Once validated, each library was sequenced on an Illumina Hi-Seq 3000 to reach a minimum number of 150M paired reads per library. Libraries from the cattle samples failed the Quality Control steps (proportion of mapped reads, number of valid interactions) and were not included in the analysis.

#### ATAC-seq

ATAC-seq libraries were prepared according to Buenrostro et al. (2013) with a few modifications. For liver, cells were dissociated from the fresh tissue to obtain a single cell suspension. Cells were counted and 50,000 cells were processed for each assay. Transposition reactions were performed using the Tn5 Transposase and TD reaction buffer from the Nextera DNA library preparation kit (Illumina) for 30 minutes at 37°C. DNA was then purified using the Qiagen MinElute PCR purification kit. Libraries were first amplified for 5 cycles using custom-synthesized index primers (see supplementary methods) and then a second amplification was performed. The appropriate number of additional PCR cycles was determined using real-time PCR, permitting the cessation of amplification prior to saturation. The additional number of cycles needed was determined by plotting the Rn versus Cycle and then selecting the cycle number corresponding to one-third of the maximum fluorescent intensity. After PCR amplification, libraries were purified using a Qiagen MinElute PCR purification kit followed by an additional clean-up and sizing step using AMPure XP beads (160 *μl* of bead stock solution was added to 100 *μl* of DNA in EB buffer) following the manufacturer’s instructions. Library quality was assessed on a BioAnalyser (Agilent) using Agilent High Sensitivity DNA kit (Agilent), and libraries were quantified using a Qubit Fluorometer (Thermo). Considering that the Hi-C protocol was not successful on the liver samples from cattle, ATAC-seq was not attempted on these samples either.

### Bioinformatics and Data Analysis

All software used in this project along with the corresponding versions are listed in Table S3. The reference gene annotation was obtained from the Ensembl v90 release (pig: Sscrofa11.1, chicken: GalGal5, cattle: UMD3.1, goat: ARS1). Since *Capra hircus* was not part of the Ensembl release, we used the NCBI CHIR_1.0.102 annotation (see Table S2).

#### RNA-seq

##### RNA-seq pipeline

Prior to any processing, all RNA-seq reads were trimmed using cutadapt version 1.8.3. Reads were then mapped twice using STAR v2.5.1.b [18, 19]: first on the genome indexed with the reference gene annotation to quantify expression of reference transcripts, and secondly on the same genome indexed with the newly generated gene annotation (FR-AgENCODE transcripts) (see below and Figure S1) [54]. The STAR `--quantMode TranscriptomeSAM` option was used in both cases in order to additionally generate a transcriptome alignment (bam) file. After read mapping and CIGAR-based softclip removal, each sample alignment file (bam file) was processed with Cufflinks 2.2.1 [55, 56] with the `max-intron-length` (100000) and `overlap-radius` (5) options, guided by the reference gene annotation (`--GTF-guide` option) ([54], Figure S1). All cufflinks models were then merged into a single gene annotation using Cuffmerge 2.2.1 [55, 56] with the `--ref-gtf` option. The transcript and gene expressions on both the reference and the newly generated gene annotation were quantified as TPM (transcripts per million) using RSEM 1.3.0 [20] on the corresponding transcriptome alignment files ([54], Figure S1). The newly generated transcripts were then processed with FEELnc version 0.1.0 [31] in order to classify them into “lncRNA”, “mRNA” and “otherRNA” (Figure S1, Tables S9-10, Figure S9). The newly generated transcripts with a TPM value of at least 0.1 in at least 2 samples were called FR-AgENCODE transcripts and kept as part of the new annotation. The 0.1 threshold was chosen knowing that the expression values of polyadenylated transcripts usually go from 0.01 to 10,000 [30] and that we wanted to simultaneously capture long non coding RNAs that are generally lowly expressed and reduce the risk of calling artefactual transcripts.

##### PCA based on gene expression

Principal Component Analysis (PCA) was performed using the **mixOmics R** package [57] on the RNA-seq sample quantifications of each species. This was done using the expression (TPM) of two different sets of genes: reference genes with TPM 0.1 in at least two samples (Figure S3) and FR-AgENCODE genes with TPM 0.1 in at least two samples (Figure S11).

##### Annotated gene orthologs

We used Ensembl Biomart [58] to define the set of orthologous genes across cattle, chicken and pig. Only “1 to 1” orthology relationships were kept (11,001 genes). Since goat was not part of the Ensembl annotation, goat gene IDs were added to this list using gene name as a correspondence term. The resulting 4-species orthologous set contained 9,461 genes (supplementary data file 3).

##### RNA-seq sample hierarchical clustering

Based on the expression of the 9,461 orthologous genes in the 39 RNA-seq samples from the four species, the sample-by-sample correlation matrix was computed using the Pearson correlation of the log_10_ gene TPM values (after adding a pseudocount of 10^−3^). We then represented this sample by sample correlation matrix as a heatmap where the samples were also clustered using a complete linkage hierarchical clustering (Figure 2).

##### RNA-seq normalization and differential analysis

To perform the differential analysis of gene expression, we used the expected read counts provided by RSEM [20]. RNA-seq library size normalization factors were calculated using the weighted Trimmed Mean of M-values (TMM) approach of [59] as implemented in the R/Bioconductor package **edgeR** [24]. The same package was used to fit three different per-gene negative binomial (NB) generalized log-linear models [60].

- In **Model 1**, the expression of each gene was explained by a tissue effect; because all three tissues (liver, CD4, CD8) were collected from each animal, an animal effect was also included to account for these repeated measures:

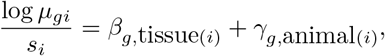

where *μ_gi_* represents the mean expression of gene *g* in sample *i, S_i_* the TMM normalization factor for sample *i*, tissue(*i*) ∈ {liver, CD4, CD8} and animal(*i*) ∈ {1, 2, 3, 4} the tissue and animal corresponding to sample *i*, and *β*_*g*, tissue(*i*)_ and *γ*_*g*,animal(*i*)_ the fixed tissue and animal effects, respectively, of gene *g* in sample *i*. Hypothesis tests were performed to identify significantly differentially expressed genes among each pair of tissues, *e.g.*

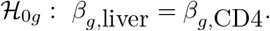
- **Model 2** is identical to the previous model, where gene expression was modeled using both a tissue and an animal effect, with the exception that the CD4 and CD8 tissues were collapsed into a single group. In this model, the only hypothesis of interest is thus between the liver and global CD cell group:

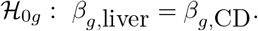

All hypothesis tests were performed using likelihood-ratio tests and were corrected for multiple testing with the Benjamini-Hochberg (FDR, [61]) procedure. Genes with an FDR smaller than 5% and an absolute log-fold change larger than 2 were declared differentially expressed.

##### GO analysis of differentially expressed genes

For each species, GO term enrichment analysis was performed on the genes found to be over-or under-expressed in liver versus T cells. This analysis was done separately for each species (Figures S5-S7) and subsequently for genes identified for all species (Figure S8), using the three following ontologies: biological process (BP), molecular function (MF) and cell compartment (CC), and using the **GOstat R**/Bioconductor package [62]) for only those genes with a human ortholog.

##### FR-AgENCODE transcript positional classification

The FR-AgENCODE transcript models were first classified according to their position with respect to reference transcripts:

known the FR-AgENCODE transcript is strictly identical to a reference transcript (same intron chain and same initial and terminal exons)
extension extension the FR-AgENCODE transcript extends a reference transcript (same intron chain but one of its two most extreme exons extends the reference transcript by at least one base pair)
alternative the FR-AgENCODE transcript corresponds to a new isoform (or variant) of a reference gene, i.e. the FR-AgENCODE transcript shares at least one intron with a reference transcript but does not belong to the above categories
novel the FR-AgENCODE transcript is in none of the above classes

##### FR-AgENCODE transcript coding classification

The FR-AgENCODE transcript models were also classified according to their coding potential. For this, the FEELnc program (release v0.1.0) was used to discriminate long non-coding RNAs from protein-coding RNAs. FEELnc includes three consecutive modules: FEELnc_filter_, FEELnc_codpot_ and FEELnc_classifier_. The first module, FEELnc_filter_, filters out non-lncRNA transcripts from the assembled models, such as transcripts smaller than 200 nucleotides or those with exons strandedly overlapping exons from the reference annotation. This module was used with default parameters except `-b transcript_biotype=protein_coding, pseudogene` to remove novel transcripts overlapping protein-coding and pseudogene exons from the reference. The FEELnc_codpot_ module then calculates a coding potential score (CPS) for the remaining transcripts based on several predictors (such as multi k-mer frequencies and ORF coverage) incorporated into a random forest algorithm [63]. In order to increase the robustness of the final set of novel lncRNAs and mRNAs, the options `--mode=shuffle` and `--spethres=0.98,0.98` were set. Finally, the FEELnc_classifier_ classifies the resulting lncRNAs according to their positions and transcriptional orientations with respect to the closest annotated reference transcripts (sense or antisense, genic or intergenic) in a 1Mb window (`--maxwindow=1000000`).

It is worth noting that between 83 and 2,718 lncRNA transcripts were not classified because of their localization on the numerous unassembled contigs in livestock species with no annotated genes.

##### FR-AgENCODE gene conservation between species

In order to obtain gene orthology relationships, we projected FR-AgENCODE transcripts from the four livestock species to the human GRCh38 genome using the UCSC pslMap program (https://github.com/ENCODE-DCC/kentUtils/tree/master/src/hg/utils/pslMap, v302). More precisely, we used the UCSC `hg38To[species.assembly].over.chain.gz` chain file for each species (created in-house for goat following UCSC instructions) and retained only the best hit for each transcript (according to the pslMap score). We further required each FRAgENCODE gene to project to a single human gene that did not strandedly overlap any other projected FR-AgENCODE gene.

##### Syntenic conservation of lncRNAs

Briefly, a lncRNA was considered as “syntenically” conserved between two species if (1) the lncRNA was located between two orthologous protein-coding genes, (2) the lncRNA was the only one in each species between the two protein-coding genes and (3) the relative gene order and orientation of the resulting triplet was identical between species. Using these criteria, we found six triplets shared between the four species, 73 triplets shared between cattle, goat and pig, and 19 triplets shared between cattle, chicken and pig.

#### ATAC-seq

##### ATAC-seq data analysis pipeline

ATAC-seq reads were trimmed with trimgalore 0.4.0 using the `--stringency 3, -q 20, —paired` and `--nextera` options (Table S3). The trimmed reads were then mapped to the genome using bowtie 2 2.3.3.1 with the `-X 2000` and `-S` options [64]. The resulting sam file was then converted to a bam file with samtools 1.3.1, and this bam file was sorted and indexed with samtools 1.3.1 [65]. The reads for which the mate was also mapped and with a MAPQ ≥ 10 were retained using samtools 1. 3.1 (`-F` 12 and `-q` 10 options, [65]), and finally only the fragments where both reads had a MAPQ ≥ 10 and which were on the same chromosome were retained.

Mitochondrial reads were then filtered out, as well as duplicate reads (with picard tools, MarkDuplicates subtool). The highest proportion of filtering was due to the MAPQ 10 and PCR duplicate filters (Figure S14). The peaks were called using MACS2 2.1.1.20160309 [66] in each tissue separately using all the mapped reads from the given tissue (`-t` option) and with the `--nomodel, -f BAMPE` and `--keep-dup all` options. To get a single set of peaks per species, the tissue peaks were merged using mergeBed version 2.26.0 [67]. These peaks were also quantified in each sample by simply counting the number of mapped reads overlapping the peak.

ATAC-seq peaks were also classified with respect to the reference gene annotation using these eight genomic domains and allowing a peak to be in several genomic domains:

exonic the ATAC-seq peak overlaps an annotated exon by at least one bp
intronic the ATAC-seq peak is totally included in an annotated intron
tss the ATAC-seq peak includes an annotated TSS
tss1Kb the ATAC-seq peak overlaps an annotated TSS extended 1Kb both 5’ and 3’
tss5Kb the ATAC-seq peak overlaps an annotated TSS extended 5Kb both 5’ and 3’
tts the ATAC-seq peak includes an annotated TTS
tts1Kb the ATAC-seq peak overlaps an annotated TTS extended 1Kb both 5’ and 3’
tts5Kb the ATAC-seq peak overlaps an annotated TTS extended 5Kb both 5’ and 3’
intergenic the ATAC-seq peak does not overlap any gene extended by 5KB on each side

##### ATAC-seq differential analysis: normalization and model

Contrary to RNA-seq counts, ATAC-seq counts exhibited trended biases visible in log ratio-mean average (MA) plots between pairwise samples after normalization using the TMM approach, suggesting that an alternative normalization strategy was needed. In particular, trended biases are problematic as they can potentially inflate variance estimates or log fold-changes for some peaks. To address this issue, a fast loess approach [68] implemented in the normOffsets function of the **R**/Bioconductor package **csaw** [69] was used to correct differences in log-counts vs log-average counts observed between pairs of samples.

As for for RNA-seq, we used two different differential models: Model 1 for tissue pair comparisons, Model 2 for T cell versus liver comparisons (see corresponding RNA-seq section for more details).

##### ATAC-seq peak TFBS density

In order to identify Transcription Factor Binding Sites (TFBS) genome-wide, we used the FIMO [70] software (Table S3) to look for genomic occurrences of the 519 TFs catalogued and defined in the Vertebrate 2016 JASPAR database [71]. We then intersected these occurrences with the ATAC-seq peaks of each species and computed the TFBS density in differential vs non differential ATAC-seq peaks. Among the predicted TFBSs, those obtained from the CTCF motif were used to profile the resulting density with respect to Topological Associating Domains from Hi-C data (Figure 7).

##### Comparison between ATAC-seq peaks and ChIP-seq histone mark peaks

Pig liver H3K4me3 and H3K27ac ChIP-seq peaks from the Villar et al. study [36] were downloaded from ArrayExpress (experiment number E-MTAB-2633). As these peaks were provided on the 10.2 pig genome assembly, they were first lifted over to the 11.1 pig genome assembly using the UCSC liftover program (https://genome.sph.umich.edu/wiki/LiftOver). About 76.8% (8,538 out of 11, 114) of the H3K4me3 peaks and 75.2% (25,502 out of 33,930) of the H3K27ac peaks could be lifted over to the 11.1 genome assembly. The median peak size was 1900 bp for H3K4me3 and 2701 bp for H3K27ac, and the peak size distribution was very similar for the initial 10.2 and the lifted over 11.1 peaks. As for genome coverage, the H3K4me3 and H3K27ac peaks covered 0.7% and 3.9% of the 11.1 pig genome, respectively. In comparison, there were 25,885 pig liver ATAC-seq peaks with a median size of 360 bp and covering 0.5% of the pig genome. Consistent with what was expected from the two histone marks, the vast majority (94.9%) of the H3K4me3 peaks (known to be associated to promoter regions) overlapped (bedtools intersect program) with the H3K27ac peaks (known to be associated to both promoter and enhancer regions), and about 30% of the H3K27ac peaks overlapped with the H3K4me3 peaks. Comparing our pig liver ATAC-seq peaks to the histone mark peaks, we found that 1.0% (250 out of 25,885) and 4.3% (1109 out of 25,885) of our ATAC-seq peaks overlapped with the H3K4me3 and H3K27ac peaks, respectively. Reciprocally, 2.8% (242 out of 8,538) and 4.1% (1,036 out of 25,502) of the H3K4me3 and H3K27ac peaks respectively overlapped with our ATAC-seq peaks, which is rather small. It was noted that, for a given kind of peak, the peaks common to the two techniques (ATAC-seq and ChIP-seq) were not longer than the other ones (not shown).

Although a rather small percentage of the ChIP-seq peaks (2.8% and 4.1%) overlapped with the ATAC-seq peaks, we wanted to know whether these numbers were higher than expected by chance. We therefore shuffled (bedtools shuffle program) the 25,885 pig liver ATAC-seq peaks 1,000 times on the pig genome and recomputed their intersection with the two sets of histone mark peaks (H3K4me3 and H3K27ac). After doing so, only 18 and 30 times did we get percentages of H3K4me3 and H3K27ac peaks, respectively, overlapping the shuffled ATAC-seq peaks that were equal or higher than the ones obtained with the real ATAC-seq peaks. This means that indeed, 2.8% and 4.1% of the histone mark peaks overlapping our ATAC-seq peaks are percentages that are significantly higher than expected by chance (p-value = 0.018 and p-value = 0.030).

We also compared the ATAC-seq, H3K4me3 and H3K27ac peak scores (fold enrichment against random Poisson distribution with local lambda for ATAC-seq peaks and fold-enrichment over background for ChIP-seq peaks) of the common peaks versus the other peaks. In doing so, we found that common peaks had significantly higher scores than non common peaks (median 49 versus 41, p-value = 8 × 10^−6^ for ATAC-seq peaks, median 59 versus 44, p-value = 1:6 ×10^−5^ for H3K4me3 peaks and median 20 versus 14, p-value < 2:2 × 10^−16^ for H3K27ac peaks, Wilcoxon tests), highlighting a common signal between the two techniques.

##### Chromatin accessibility conservation across species

In order to investigate the conservation of chromatin accessibility across our 4 livestock species, we used the human GRCh38 genome as a reference. After indexing the softmasked GRCh38 genome (main chromosomes) using lastdb (last version 956, -uMAM4 and -cR11 options, http://last.cbrc.jp/), we used the lastal program followed by the last-split program (-m1 and –no-split options) (last version 956, http://last.cbrc.jp/) to project the 104,985 cattle, 74,805 goat, 119,894 chicken and 149,333 pig ATAC-seq peaks onto the human genome. In doing so and consistent with the phylogenetic distance between our species and human, we were able to project 72.6% (76,253) cattle, 73.7% (55,113) goat, 12.3% (14,792) chicken and 80.1% (119,680) pig peaks to the human genome. The percentage of bases of the initial peaks that could be aligned was around 40% for mammals and 14% for chicken. Then, for each peak that could be projected onto the human genome, we retained its best hit (as provided by lastal) and then merged all these best hits (i.e. from the 4 species) on the human genome (using bedtools merge). A total of 215,620 human regions were obtained, from which we kept the 212,021 that came from a maximum of 1 peak from each species. Those 212,021 regions were called human hits.

Based on the 1,083 four-species orthologous peaks in the 38 ATAC-seq samples, the sample-by-sample correlation matrix was computed using the Pearson correlation of the log_10_ normalized ATAC-seq values (after adding a pseudo-count of 10^−3^ to the values). We then represented this sample-by-sample correlation matrix as a heatmap where the samples were also clustered using a complete linkage hierarchical clustering (Figure S21). Chicken ATAC-seq samples clustered completely separately from mammal ATAC-seq samples. T cell samples from cattle and goat were also closer to each other than to liver samples.

To shuffle the 104,985 cattle, 74,805 goat, 119,894 chicken and 149,333 pig ATAC-seq peaks we used the bedtools shuffle program on their respective genomes and projected these shuffled peaks to the human genome as was done for the real peaks.

We also compared the human hits to the combined set of 519,616 ENCODE human DNAse I peaks from two CD4+, two CD8+ and one “right lobe of liver” samples (experiment accessions ENCSR683QJJ, ENCSR167JFX, ENCSR020LUD, ENCSR316UDN, ENCSR555QAY from the encode portal https://www.encodeproject.org/, by merging the peaks from the 5 samples into a single set of peaks using bedtools merge). We found that 23.1% (48,893 out of 212,021) of the human hits obtained from the real ATAC-seq peaks overlapped human DNAse I peaks, whereas only 8.5% (21,159 out of 249,943) of the human hits obtained from shuffled ATAC-seq peaks overlapped human DNAse I peaks. This further supports the biological signal present in these data.

Finally we used the phastcons measure of vertebrate sequence conservation obtained from the multiple alignment of 100 vertebrate species genomes including human (hg38.phastCons100way.bw bigwig file from the UCSC web site https://genome.ucsc.edu/). For each human hit, we computed its phastcons score using the bigWigAverageOverBed utility from UCSC (https://github.com/ENCODE-DCC/kentUtils).

#### Hi-C

##### Hi-C data analysis pipeline

Our Hi-C analysis pipeline includes HiC-Pro v2.9.0 [72] (Table S3) for the read cleaning, trimming, mapping (this part is internally delegated to Bowtie 2 v2.3.3.1), matrix construction, and matrix balancing ICE normalization [73]. HiC-Pro parameters: BOWTIE2_GLOBAL_OPTIONS = --very-sensitive -L 30 --score-min L,-1,-0.1 --end-to-end --reorder, BOWTIE2_LOCAL_OPTIONS = --very-sensitive -L 20 --score-min L,-0.6,-0.2 --end-to-end --reorder, LIGATION_SITE = AAGCTAGCTT, MIN_INSERT_SIZE = 20, MAX_INSERT_SIZE= 1000, GET_ALL_INTERACTION_CLASSES = 1, GET_PROCESS_SAM = 1, RM_SINGLETON = 1, RM_MULTI = 0, RM_DUP = 1, MAX_ITER = 100,FILTER_LOW_COUNT_PERC = 0.02, FILTER_HIGH_COUNT_PERC = 0, EPS =0.1. TAD finding was done using Armatus V2.1 [43] with default parameters and `gamma=0.5` (Table S3). Graphical visualization of the matrices was produced with the **HiTC R**/Bioconductor package v1.18.1 [42] (Table S3). Export to JuiceBox [74] was done through Juicer Tools v0.7.5 (Table S3). These tools were called through a pipeline implemented in Python. Because of the high number of unassembled scaffolds (e.g. for goat) and/or micro-chromosomes (e.g. for chicken) in our reference genomes, only the longest 25 chromosomes were considered for TAD and A/B compartment calling. For these processes, each chromosome was considered separately.

The Directionality Index (DI) was computed using the original definition introduced by [44] to indicate the upstream vs. downstream interaction bias of each genomic region. Interaction matrices of each chromosome were merged across replicates and the score was computed for each bin of 40Kb. CTCF sites were predicted along the genomes by running FIMO with the JASPAR TFBS catalogue (see section “ATAC-seq peak TFBS density”).

A/B compartments were obtained using the method described in [17] as illustrated in Figure S2: first, ICE-normalized counts, *K_ij_*, were corrected for a distance effect with:

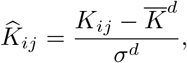

in which 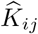 is the distance-corrected count for the bins *i* and *j*, 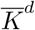 is the average count over all pairs of bins at distance *d* = *d*(*i, j*) and *σ^d^* is the standard deviation of the counts over all pairs of bins at distance *d*. Within-chromosome Pearson correlation matrices were then computed for all pairs of bins based on their distance-corrected counts and a PCA was performed on this matrix. The overall process was performed similarly to the method implemented in the **R**/Bioconductor package **HiTC** [42]. Boundaries between A and B compartments were identified according to the sign of the first PC (eigenvector). Since PCAs had to be performed on each chromosome separately, the average counts on the diagonal of the normalized matrix were used to identify which PC sign (+/−) should be assigned to A and B compartments for each chromosome. This allowed a homonegenous assignment across chromosomes to be obtained, without relying on the reference annotation. In line with what was originally observed in humans, where the first PC was the best criterion for separating A from B compartments (apart from a few exceptions like chromosome 14 for instance [17]), we also observed a good agreement between the plaid patterns of the normalized correlation matrices and the sign of the first PC (Figure S24).

To estimate the robustness of A/B compartment calling, the method was tested on each replicate separately (four animals). Since the HiTC filtering method can discard a few bins in some matrices, resulting in missing A/B labels, the proportion of bins with no conflicting labels across replicates was computed among the bins that had at least two informative replicates (Figure S27).

##### Chromatin structure conservation across species

To get insight into chromatin structure conservation across species, similar to what was done with chromatin accessibility data (see above), we projected the 11,711 goat, 6,866 chicken and 14,130 pig 40kb TAD boundaries to the human GRCh38 genome using lastal followed by last-split (`-m1` and `--no-split` options, last version 956, http://last.cbrc.jp/, using the same indexed GRCh38 softmasked genome as was used for ATAC-seq, see above). As expected from their length, TAD boundary projections were highly fragmented (median 16, 2 and 19 blocks per projection representing 3%, 0.6% and 3% of the initial segment, for the best hit of goat, chicken and pig, respectively). In order to recover conserved segments, we chained the alignments using a python in-house script (program available on demand, used with stranded mode, `coverage=0.4`, `score=3000` and `length_cutoff=5000`). Doing so, we managed to project 90% of the mammalian and 5% of the chicken TAD boundaries onto the human genome. Similar to what was done for ATAC-seq, for each projected TAD boundary, its best hit (according to the chaining score) was retained. The median length of those best hits represented 95% and 78% of the initial query size for mammals and chicken respectively. Merging these best hits on the human genome (using bedtools merge), we obtained 16,870 human regions with a median length of 44.6kb (similar to the initial TAD boundary size of 40kb). Out of those, 16,468 were considered non ambiguous (i.e. not coming from several TAD boundaries from the same species) and were retained for further analyses. As was found for the ATAC-seq peaks, the majority (65.6%) of the hits were single species (orthology level 1), a substantial percentage of them (34%) were 2 species hits (orthology level 2), and seventy one of them (0.4%) were 3 species hits (orthology level 3).

To estimate the structural impact of each TAD boundary, we used the local interaction score as used by [45] and [46] and sometimes referred to as “interaction ratio” or “insulation profile”. Within a sliding window of 500kb along the genome (step=10Kb), the insulation score ratio is defined as the proportion of read pairs that span across the middle of the window. The score ratio is reported at the middle position of the window and represents the local density of the chromatin contacts around this point. This proportion is expected to be maximal in regions with many local interactions (typically TADs) and minimal over insulators (typically TAD boundaries). Intuitively, a TAD boundary with a low interaction score (which indicates strong insulation properties) has a good capacity to prevent interactions that cross it while a TAD with a relatively high interaction score has a “weak” insulation strength. Here, only valid interactions (“valid pairs”) in *in cis* (inter-chromosomal contacts were not considered in the ratio) were considered after applying all HiC-Pro QC and filters. Computing a ratio among all read pairs that have both reads within the sliding window reduces the impact of potential biases (read coverage, restriction site density, GC content, etc). Consequently, the interaction profiles from the 4 replicates along the genome of each species were highly similar (not shown), allowing to merge them in order to assign each TAD boundary a single score per species. For orthologous TAD boundaries, the scores from different species could be used to compute pairwise correlations. Human data were obtained from http://aidenlab.org/data.html [41] for the GM12878 cell line (ENCODE batch 1, HIC048.hic file from https://bcm.app.box.com/v/aidenlab/file/95512487145). The.hic file was parsed by the juicer tool (“dump” mode with options “observed KR”) to compute the corresponding interaction score as described above. The LiftOver tool was used to convert the genomic positions of the human TAD boundaries (version hg19 vs. hg38) before comparing the interaction scores with livestock species.

The number and proportion of genes (all or only the orthologous ones) in each compartment type was computed using bedtools map (`-distinct` option on the gene ID field). Orthologous genes were taken from Ensembl as previously described. Under the independence assumption of compartment assignment between species, the expected proportion of orthologous genes with “triple A” (resp. with “triple B”) assignments between species is equal to the product of the observed frequencies for A (resp. for B) compartments in the three species. The observed frequencies of “triple A” and “triple B” assignments in orthologous genes was compared to this expected proportion using a *χ*^2^ goodness-of-fit test.

#### Multi-assay integration

##### ATAC-seq vs. RNA-seq correlation: intra- and inter-sample analysis

For each ATAC-seq peak that overlapped a promoter region (1Kb upstream of the TSS, as suggested in Figure 4) its loess-normalized read count value (see differential analysis) was associated with the TMM-normalized expression of the corresponding gene from the reference annotation. Intra- and inter-sample correlations were then investigated: within each sample, genes were ranked according to their expressions and the distribution of the corresponding ATAC-seq values was computed for each quartile (Figure S17). Across samples, the Pearson correlation coefficient was computed for each gene using only the samples for which both the ATAC-seq and the RNA-seq normalized values were available (e.g. *n* =10 for pig, Figures S18-19). Similar results were obtained with Spearman correlations (not shown).

##### Chromatin accessibility and gene expression in A/B compartments

To compute the general chromatin accessibility in A and B compartments, we first computed the average of the normalized read count values across all liver samples for each ATAC-seq peak. For each compartment, the mean value of all contained peaks was then reported and the resulting distributions for all A and B compartments were reported (Figure 8).

**Figure 9.**
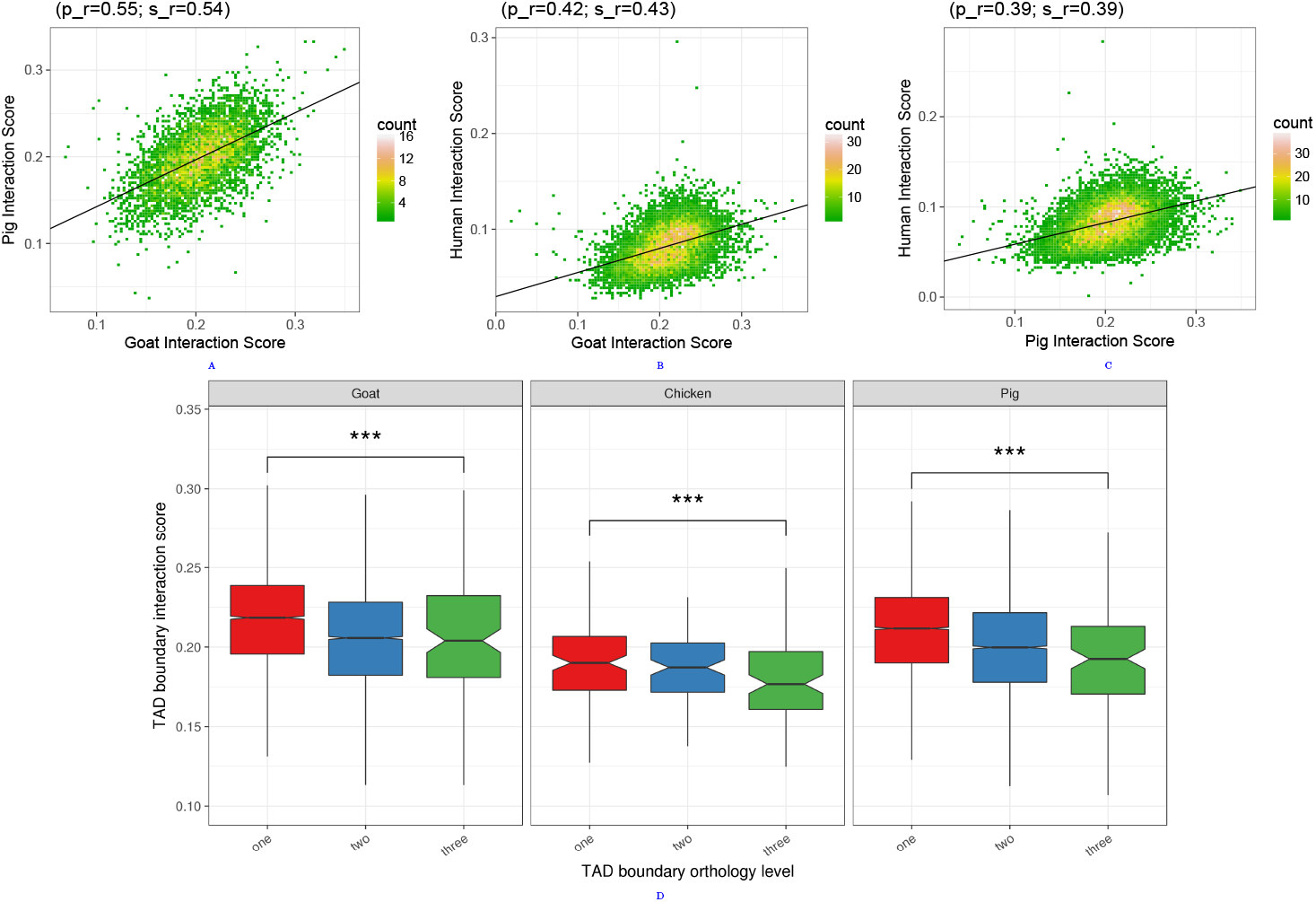
Relationship between chromatin structure conservation and functionality. Interaction scores of orthologous TAD boundaries between Goat and Pig (A), Goat and Human (B) and Pig and Human (C). (D) For each species with Hi-C data, TAD boundaries were divided according to their orthology level (1, 2 and 3, *x*-axis and boxplot colours) and their interaction scores were plotted (*y*-axis). There is a clear decrease of the interactions core with the TAD boundary orthology level, indicating a stronger insulation for more evolutionary conserved TAD boundaries.

The same approach was used to assess the general expression of genes in A and B compartments, using the average of the normalized expression values from the liver samples. Difference between A and B distributions was tested for statistical significance using a Wilcoxon test.

## List of abbreviations

DE: Differentially Expressed
FAANG: Functional Annotation of Animal Genomes
lncRNA: long non-coding RNA
mRNA: messenger RNA
PE: Paired-End
polyA: polyAdenylation
RT-PCR: Reverse Transcriptase Polymerase Chain Reaction
TAD: Topological Associating Domain
TF: Transcription Factor
TFBS: Transcription Factor Biding Site
TPM: Transcript Per Million
TSS: Transcription Start Site
TTS: Transcription Termination Site

## Declarations

### Ethics approval and consent to participate

All animal handling and sampling was realized in conformity with the French legislation on animal experimentation. For mammals, blood samples were collected in the context of the following approvals (APAFIS/project#): 334-2015031615255004_v4 and 333-2015031613482601_v4 (pigs), 3066-201511301610897_v2 (cattle), 03936.02 and 8613-2017012013585646_v4 (goats). Chicken immune cells were obtained from spleen sampled after slaughter (no need for animal experiment authorization).

### Consent for publication

Not applicable.

### Availability of data and material

Experimental protocols for tissue sampling and molecular assays are available at the FAANG portal of the EMBL-EBI ftp website: ftp://ftp.faang.ebi.ac.uk/ftp/protocols/samples/. Sample records are available at the BioSamples database using the keyword “FR-AgENCODE” (submission codes GSB-99 and GSB-721). Biological material from the 16 animals (tissue samples and aliquots) are available at the INRA CRB-Anim BioBanking facility on request. Raw sequencing reads are available in fastq format at the EMBL-EBI’s European Nucleotide Archive ENA using the keyword “FR-AgENCODE” or under the accessions PRJEB27455 (RNA-seq), PRJEB27111 (ATAC-seq) and PRJEB27364 (Hi-C). All the corresponding metadata information has been provided to the FAANG Data Coordination Center. Additional data and results (including annotation files, list of differentially expressed genes with normalized expression values, annotated ATAC-seq peaks with raw read counts, differentially accessible peaks with normalized read counts, Hi-C matrices, TADs and A/B compartments) are available at INRA’s FR-AgENCODE website: www.fragencode.org.

### Authors’ Contributions

Animal and sampling: MTB and SFa (coordination), AG, EG, FB, FD, FL, GTK, HA, MTB, PQ, SDP, SFa, SLL, SVN (sampling and cell sorting). Molecular Assays: DE and HA (coordination), DE, SDP (RNA-seq libraries), AG, EG, KMun (ATAC-seq libraries), FM, HA, MM (Hi-C libraries). Bioinformatics and Data Analysis: CK, SD, SFo (coordination), CC, SD (RNA-seq), KMun, SD, SFo (ATAC-seq), KMur, SL, TD (lncRNAs), AR, NV, RML (differential analyses), DR, IG, MM, MSC, MZ, NV, SFo (Hi-C pipeline), EC, EG, SD, SL, TD (functional analyses), SD, SFo (metadata), AR, NV, SD, SFo, TF (integrative analyses), HA, SD, SFo, SM, PB (data submission). Manuscript writing: AR, CK, EG, HA, KMun, KMur, MTB, MZ, NV, SD, SFo, SL, TD. Management committee: EG, MHP, SFo, SL. Project coordination: EG, SFo. All authors have read and approved the manuscript.

## Acknowledgements

We would like to thank the FAANG community for the general support and in particular A. Archibald (Roslin Institute, UK), L. Clarke, P. Harrison (EMBL-EBI, UK) and M. Groenen (WUR, Netherlands) for their collaboration in the organization of the project, and B. Rosen (ARS, USDA) for providing information about the goat sexual chromosomes. We wish to thank all field operators at INRA experimental animal facilities, units and platforms in France for the access and the assistance in animal handling and sampling, including: Y. Gallard, UE Le Pin (Gouffern en Auge), F. Bouvier and T. Fassier, UE Bourges (Osmoy), S. Ferchaud, UE GenESI (Magneraud/Rouille), Y. Baumard, UE PEAT (Nouzilly), C. Berri and J. Gautron, UR BOA (Nouzilly), G. Gomot, J.P. Dubois and A. Arnould, UR PRC CIRE (Nouzilly), E. Guettier, D. Capo and J. Savoie, UE PAO (Nouzilly).

We are grateful to A. Breschi (Stanford, USA) and J. Lagarde (CRG, Spain) for sharing scripts and to N. Servant (Institut Curie, France) for assistance on HiC-PRO. Additional acknowledgements go to C. Donnadieu and O. Bouchez from the Get-Plage sequencing platform (INRA Toulouse) and to C. Gaspin and her staff at the GenoToul bioinformatics platform (INRA Toulouse), especially D. Laborie and M.S. Trotard for IT support.

## Supplementary data files

Supplementary data files are available in the supplementary data file section and on the FR-AgENCODE website www.fragencode.org.

### Supplementary data file 1 — SF1.pdf

Supplementary figures (S1-S26) and tables (S1-S15).

### Supplementary data file 2 – refgn.tar.gz

Reference genes and transcripts (structure, expression) of the 4 species. Archive content:

- `bos_taurus.gtf`
- `bos_taurus.refgn.tpm.tsv`
- `capra_hircus.gtf`
- `capra_hircus.refgn.tpm.tsv`
- `gallus_gallus.gtf`
- `gallus_gallus.refgn.tpm.tsv`
- `sus_scrofa.gtf`
- `sus_scrofa.refgn.tpm.tsv`

### Supplementary data file 3 – refgn.orth.tar.gz

Orthologs between the 4 livestock species. We used Biomart to retrieve the 1 to 1 orthology relationships between chicken, pig and cattle and added goat via gene name. The human gene id is given for reference.

### Supplementary data file 4 – de.refgn.tar.gz

Reference DE genes (all combinations): the archive contains four folders, one for each species (`bos_taurus`, `capra_hircus`, `gallus_gallus`, `sus_scrofa`). Each folder contains itself two subfolders, one for each model: diffcounts.nominsum (Model 1) and `diffcounts.cdvsliver` (Model 2).

Results of Model 1 are given in:

- `refgenes.counts.min2tpm0.1.normcounts.diff.readme.idx`
- `refgenes.counts.min2tpm0.1.normcounts.diff.cd4.cd8.bed`
- `refgenes.counts.min2tpm0.1.normcounts.diff.cd4.liver.bed`
- `refgenes.counts.min2tpm0.1.normcounts.diff.cd8.liver.bed`

Results of Model 2 are given in:

- `refgenes.counts.min2tpm0.1.normcounts.diff.readme.idx`
- `refgenes.counts.min2tpm0.1.normcounts.diff.cd.liver.bed`

All bed files contain the coordinates and id of the genes found to be differentially expressed between the two conditions. The file also contains the normalized read counts of those genes in the different samples as well as the adjusted pvalue, logFC and normLogFC (see readme.idx file for more details)

### Supplementary data file 5 – fraggn.tar.gz

FR-AgENCODE genes and transcripts (structure, expression, positional and coding classes).

- `bos_taurus_cuff_tpm0.1_2sample_complete.gff`
- `bos_taurus_cuff_tpm0.1_2sample_trid_4posclasses_3codingclasses_booleans.tsv`
- `bos_taurus.frag.gnid.posclasslist.codclasslist.tsv`
- `bos_taurus.fraggn.tpm.tsv`
- `capra_hircus_cuff_tpm0.1_2sample_complete.gff`
- `capra_hircus_cuff_tpm0.1_2sample_trid_4posclasses_3codingclasses_booleans.tsv`
- `capra_hircus.frag.gnid.posclasslist.codclasslist.tsv`
- `capra_hircus.fraggn.tpm.tsv`
- `gallus_gallus_cuff_tpm0.1_2sample_complete.gff`
- `gallus_gallus_cuff_tpm0.1_2sample_trid_4posclasses_3codingclasses_booleans.tsv`
- `gallus_gallus.frag.gnid.posclasslist.codclasslist.tsv`
- `gallus_gallus.fraggn.tpm.tsv`
- `sus_scrofa_cuff_tpm0.1_2sample_complete.gff`
- `sus_scrofa_cuff_tpm0.1_2sample_trid_4posclasses_3codingclasses_booleans.tsv`
- `sus_scrofa.frag.gnid.posclasslist.codclasslist.tsv`
- `sus_scrofa.fraggn.tpm.tsv`

### Supplementary data file 6 – fraggn.orth.tar.gz

Four livestock species FR-AgENCODE gene orthology

### Supplementary data file 7 – de.fraggn.tar.gz

FR-AgENCODE DE genes (all combinations). The archive has the same structure than `de.refgn.tar.gz` with names starting with cuffgenes instead of `refgenes`.

### Supplementary data file 8 – lncRNAs.tar.gz

lncRNAs (information from FEELnc, orthology, structure, etc). Archive content:

- `bos_taurus.lncrna.TPM0.1in2samples.classif.tsv`
- `capra_hircus.lncrna.TPM0.1in2samples.classif.tsv`
- `ConservedLncRNABySynteny_73_19_6.xlsx`
- `gallus_gallus.lncrna.TPM0.1in2samples.classif.tsv`
- `sus_scrofa.lncrna.TPM0.1in2samples.classif.tsv`

### Supplementary data file 9 – atac.peaks.tar.gz

ATAC-seq peaks (coordinates, quantification, positional classification): the archive contains four folders, one for each species (`bos_taurus`, `capra_hircus`, `gallus_gallus`, `sus_scrofa`). Each folder contains the following six files:

- `mergedpeaks_allinfo_gn_frag.tsv`
- `mergedpeaks_allinfo_tr_frag.tsv`
- `mergedpeaks_allinfo_tr_ref.tsv`
- `mergedpeaks_allinfo_gn_ref.tsv`
- `mergedpeaks.peaknb.allexp.readnb.bed.readme.idx`
- `mergedpeaks.peaknb.allexp.readnb.bed`

### Supplementary data file 10 – da.atacpeaks.tar.gz

DA ATAC-seq peaks (all combinations). The archive has the same structure as `de.refgn.tar.gz` with names starting with `mergedpeaks.peaknb.allexp.readnb` instead of `refgenes.counts.min2tpm0.1`.

### Supplementary data file 11 – atac.peaks.orth.tar.gz

Four livestock species ATAC-seq peak orthology

### Supplementary data file 12 – hic.tad.ab.tar.gz

Hi-C TADs and A/B compartments: the archive contains three folders, one for each species (`capra_hircus`, `gallus_gallus`, `sus_scrofa`). Each folder contains the following two files:

- `compartments.bed`
- `mat.40000.longest25chr.tad.consensus.bed`

### Supplementary data file 13 – hic.bound.orth.tar.gz

Three livestock species TAD boundary orthology

