## Supplementary data file 1 for "Transcriptome and chromatin structure annotation of liver, CD4+ and CD8+ T cells from four livestock species"

### Supplementary file 1: Supplementary figures and tables

### List of Figures


S1 **RNA-seq processing pipeline overview.** RNA-seq reads were processed with STAR for mapping and Cufflinks for transcript modeling using the reference gene annotation as input and for each sample individually. Modelled transcripts were then merged with Cuffmerge to produce a single gene annotation per species, and the transcripts from this novel gene set were further processed with FEELnc to produce a reliable set of lncRNAs. Reference and novel transcripts and gene expressions were quantified in each sample using the STAR/RSEM pipeline. Differential expression analysis was performed on reference and novel genes with TPM $\geq$ 0.1 in at least 2 samples using the R/Bioconductor package **edgeR**. . . . . . . . . . . 6

S2 **RNA-seq mapped read classification into genomic domains.** For each sample, RNA-seq mapped reads were classified into exonic (1_exon), intronic (2_intr), intergenic (3_ig) and other (4_other) reads. . . . . . . . . . . . . . . . . . . . . . . . . . . . . . . . . . . . . . . . 7

S3 **RNA-seq sample PCA based on the expression of reference genes.** Only reference genes with a TPM $\geq$ 0.1 in at least 2 samples were used. . . . . . . . . . . . . . . . . . . . . . 8

S4 **Gene expression ratio in males versus females on chromosomes X or Z.** Gene expression was compared in males vs. females using the base 10 logarithm of the TPM values. $y$-axis: the $\log_2$ of the male/female expression ratio for the genes located on the chromosome of interest. For each plot, the median of these expression ratios is indicated in the title. Left: genes located on chromosome 1 used as a control. Right: genes located on chromosome X for mammals and Z for chicken. $x$-axis: the position of the gene on the chromosome of interest. 9

S5 **Biological Process GO term enrichment analysis for reference genes differentially expressed between liver and T cells.** This analysis was performed for each species individually (in columns: cattle, goat, chicken, pig in that order) and for genes over-expressed in liver (top) and over-expressed in T cells (bottom). . . . . . . . . . . . . . . . . . . . . . . . . 10

S6 **Molecular Function GO term enrichment analysis for reference genes differentially expressed between liver and T cells.** This analysis was performed for each species individually (in columns: cattle, goat, chicken, pig in that order) and for genes over-expressed in liver (top) and over-expressed in T cells (bottom). . . . . . . . . . . . . . . . . . . . . . . . . 11

S7 **Cellular Compartment GO term enrichment analysis for reference genes differentially expressed between liver and T cells.** This analysis was performed for each species individually (in columns: cattle, goat, chicken, pig in that order) and for genes over-expressed in liver (top) and over-expressed in T cells (bottom). . . . . . . . . . . . . . . . . . . . . . . . . 12

S8 **GO term enrichment analysis for reference genes differentially expressed between liver and T cells in all species.** This analysis was performed for each GO (in column: Biological Process, Molecular Function, Cellular Compartment in that order) and for genes over-expressed in liver (top) and over-expressed in T cells (bottom). . . . . . . . . . . . . . . . . . 13

S9 **Distribution of FR-AgENCODE transcripts into four positional (known, extension, alternative, novel) and three coding (mRNA, lncRNA, otherRNA) classes. (A)** Distribution of all FR-AgENCODE transcripts into positional and coding classes; **(B)** and **(C)** Distribution of FR-AgENCODE transcripts of each coding class into positional classes, and of each positional class into coding classes respectively. . . . . . . . . . . . . . . . . . . . . . . . . . . 14

S10 **Novel coding FR-AgENCODE genes enrich the set of blood and T cell annotated genes.** (A) Venn diagrams of novel coding genes for each triplet of livestock species (two livestock genes are defined as orthologous if they project to the same human gene). (B) Gene set enrichment analysis on the human orthologs of the 93 cattle, 52 goat, 74 chicken and 26 pig genes, using EnrichR (Kuleshov et al. 2016; `http://amp.pharm.mssm.edu/Enrichr/`) on the ARCS4 Tissue dataset (`https://amp.pharm.mssm.edu/archs4/`). (C) Gene names of the 12 genes that are common to two livestock species. Out of these 12, 8 are coding for T cell Receptor Alpha or Beta Variable genes. . . . . . . . . . . . . . . . . . . . . . . . . . . . 15

S11 **RNA-seq sample PCA based on FR-AgENCODE gene expression.** . . . . . . . . . . . 16

S12 **LncRNA gene features.** Genomic structure features (left) and expression (right) are indicated for protein coding genes (coding) and lncRNA genes (lnc). The expression (TPM) is given for the three tissues and the four species; lncRNA genes are between 8- and 43-fold less expressed than protein coding genes (expression median: 0.4 vs. 10 TPM respectively). . . . 17

S13 **Syntenic lncRNAs conserved between the 4 livestock species and human.** The 6 syntenic lncRNAs conserved in the 4 livestock species and human are represented in green and their surrounding protein coding genes in orange. Distances between the genes are indicated either in base pair or in kilo base pair (k). A distance of 0 means the genes are overlapping. A distance in red means a lower confidence in the orthology relationship for this species, after inspection of the distances found in the other species. When the lncRNA was not known before, a new gene name is proposed (asterisk). . . . . . . . . . . . . . . . . . . . . . . . . . . 18

S14 **ATAC-seq read pair summary statistics.** For each species and sample (labelled by its tissue and animal number), the number of initial read pairs (1_init), of read pairs obtained after trimming (2_trim), mapping (3_map), proper pairing (4_pair), q10 filtering (5_q10), mitochondrial read removal (6_nonmt), and PCR duplicate read removal (7_nondup) are shown. . . . . . . . 19

S15 **ATAC-seq peak size distribution.** For each species, the size distribution of tissue and merged (i.e. across all tissues) ATAC-seq peaks are provided. . . . . . . . . . . . . . . . . . 20

S16 **Density of ATAC-seq peaks around starts of novel (i.e. not known) transcripts for cattle (A), goat (B), chicken (C) and pig (D)** . . . . . . . . . . . . . . . . . . . . . . . . . . . . . . . 21

S17 **ATAC-seq peak distribution into genomic domains.** For each species, are provided: the percentage of ATAC-seq peaks that are exonic, intronic, overlapping the TSS of a reference gene extended by 0, 1 or 5 Kb on each side, overlapping the TTS of a reference gene extended by 0, 1 or 5 Kb on each side and intergenic. . . . . . . . . . . . . . . . . . . . . . . . . . . 22

S18 **Per sample promoter accessibility for four reference gene expression quartiles in pig.** 23

S19 **Schematic illustration of the per-gene and cross-sample RNA-seq versus ATAC-seq correlation analysis.** . . . . . . . . . . . . . . . . . . . . . . . . . . . . . . . . . . . . . . . . 24

S20 **Correlation between reference gene expression and chromatin accessibility at the TSS for goat.** For each pair of reference gene and ATAC-seq peak overlapping the reference gene TSS extended by 1 Kb on each side, the Pearson correlation was computed between the base 10 logarithm of the gene TMM and the base 10 logarithm of the normalized ATAC-seq signal at the peak. The distribution was then plotted for non DE genes **(A, top)** and for DE genes **(B, bottom)**. . . . . . . . . . . . . . . . . . . . . . . . . . . . . . . . . . . . . . . . . . . . 25

S21 **ATAC-seq sample heatmap and hierarchical clustering based on the 1083 level 4 ATAC-seq peaks.** Pairwise similarity between samples is computed as the Pearson correlation between the base 10 logarithm of the normalized reads of the 1083 ATAC-seq peaks common to 4 species. These similarities are plotted as a heatmap, where samples appear both as rows and columns and are labelled by their species, tissue and the sex of the animal. The color of each heatmap cell also reflects the similarity (Pearson correlation) between each sample pair (the lighter, the higher). Hierarchical clustering is performed using one minus the squared Pearson correlation as a distance and the complete linkage aggregation method. . . . . . . . 26

S22 **Relation between non promoter proximal chromatin accessibility conservation and differential accessibility.** This figure is the same as main Figure 6 but is done restricting to ATAC-seq peaks not overlapping the TSS+-1kb in any of the species where it is presen. Phastcons scores of these ATAC-seq peaks were plotted after dividing the human hits according to both their orthology level (between 1 and 3, x-axis, level 4 removed because there were less than 50 peaks in a category) and their differential accessibility (DA) status (DA in at least one species or DA in none of the 4 species, boxplot color). Although the phastcons score obviously increases with the orthology level, it is clear that, for a given orthology level, the phastcons score is higher for DA human hits than for non DA human hits (all orthology levels, p-values ¡ 1.2e-6 overall, Wilcoxon tests) (number of elements the boxplots from left to right: 151,783, 19,732, 11,922, 3,317, 2,100, 1,082). . . . . . . . . . . . . . . . . . . . . . . . . . . . . . . 27

S23 **Hi-C read summary statistics.** For each species and animal, are plotted the number of: initial read pairs after sequencing (1_initial), pairs with both reads mapped on the genome (2_reported), pairs in a valid configuration, that is when the sum of the distances from the reads to their next HindIII restriction sites downstream is comprised between 20bp and 1Kb (3_valid), pairs after removing PCR duplicates (4_valid_rmdup) and the number of read pairs supporting proximity between two different chromosomes (5_trans). . . . . . . . . . . . . . 28

S24 **Method for predicting Hi-C A and B compartments.** Illustration of the A/B compartment calling workflow using the interaction matrices of the first chromosome in pig. Upper panels: interaction matrices at different steps of the workflow. Lower panel: the first three eigenvectors are shown along the chromosome to illustrate the relevance of PC#1 as the discriminative value to segregate bins between A and B compartments. . . . . . . . . . . . . . . . . . . . . 29
S25 **Juicebox Hi-C maps of three livestock species' chromosome 1.** Each column represents a different species (goat, chicken, pig). For a given column, the first map represents the whole chromosome 1 Hi-C heatmap at 500Kb resolution. On this map the blue square represents a zone that is further enlargeg on the second map, itself representing part of chromosome 1 Hi-C heatmap at 40Kb resolution. The third map represents the chromosome 1 bin pair Pearson correlation matrix computed from the 500Kb resolution Hi-C matrix, and that allows to see the A and B compartments. . . . . . . . . . . . . . . . . . . . . . . . . . . . . . . . . 30
S26 **Directionality Index (DI) around Hi-C TADs.** . . . . . . . . . . . . . . . . . . . . . . . . . 31
S27 **Distribution of Hi-C A and B compartments along each chromosome and for each animal.** Genome-wide overview of compartment labels per 500Kb bin in pig for each animal. A general coherence can be observed across replicates. White regions are devoid of any called compartment. . . . . . . . . . . . . . . . . . . . . . . . . . . . . . . . . . . . . . . . . . . . 32

### List of Tables

S1 **Completed experiments.** Available data per animal, sample and experimental assay. Green: completed experiment. Red (NA): not available. . . . . . . . . . . . . . . . . . . . . . . . . . 34
S2 **Genome and reference gene annotation used for each species.** . . . . . . . . . . . . . . 35
S3 **Software used in the FR-AgENCODE project.** . . . . . . . . . . . . . . . . . . . . . . . . . 36
S4 **RNA-seq read mapping statistics.** Number and proportion of mapped and uniquely mapped RNA-seq read pairs for each species, replicate and tissue. . . . . . . . . . . . . . . . . . . 38
S5 **Differentially Expressed (DE) reference genes.** Number of differentially expressed reference genes obtained by the two statistical models (see main text and Methods) . . . . . . . 39
S6 **Genes consistently over-expressed in CD4+ compared to CD8+ or reciprocally, in four livestock species.** For the 39 genes consistently seen as over-expressed in CD4+ with respect to CD8+ (10 genes) or reciprocally (29 genes), are indicated: the human gene ID, the gene name in a column that indicates the cell type in which the gene is over-expressed, the TPM in human CD4+ and CD8+ cells from the Blueprint project (CD4-positive, alpha-beta T cell and CD8-positive, alpha-beta T cell respectively), whether or not these expression levels are consistent with the differential behavior observed in livestock, the expression in livestock CD4+ and CD8+ (average across samples) and a reference describing the role of the gene in blood cells when available. . . . . . . . . . . . . . . . . . . . . . . . . . . . . . . . . . . 40
S7 **FR-AgENCODE transcript positional classification.** . . . . . . . . . . . . . . . . . . . . . 41
S8 **FR-AgENCODE novel coding genes and their orthology with human** . . . . . . . . . . . 42
S9 **Differentially Expressed (DE) FR-AgENCODE genes.** Number of differentially expressed FR-AgENCODE genes obtained by the two statistical models (see main text and Methods). . 43
S10 **lncRNA classification.** Number of classified expressed lncRNAs per species. This table includes monoexonic lncRNAs that represent 57-68% of these transcripts. Loci are bracketed. Fields with an asterisk (*) indicate the existence of lncRNAs unclassified by FEELnc because they are on unassembled contigs; they represent 217, 717, 2,718 and 83 lncRNAs in cattle, goat, chicken and pig respectively and are not listed here. . . . . . . . . . . . . . . . . . 44
S11 **FR-AgENCODE transcript coding classification.** . . . . . . . . . . . . . . . . . . . . . . . 45
S12 **Number of ATAC-seq peaks per species.** . . . . . . . . . . . . . . . . . . . . . . . . . . . 46
S13 **Number of differentially accessible (DA) ATAC-seq peaks per species.** . . . . . . . . . . 47
S14 **Hi-C read pair mapping statistics.** Number of read pairs of different categories. *Initial*: total number of sequenced read pairs. *Reported*: pairs with both reads mapped on the genome. *Valid*: uniquely mapped pairs with an estimated insert size (sum of the distances from the reads to their next downstream HindIII genomic sites) between 20bp and 1Kb. *Valid.rmdup*: valid read pairs after duplication removal that were used to build the interaction matrices. *Trans*: pairs with reads on different chromosomes. . . . . . . . . . . . . . . . . . . . . . . . 48

S15 **Statistics of Hi-C TADs and A/B compartments.** . . . . . . . . . . . . . . . . . . . . . . . . . 49

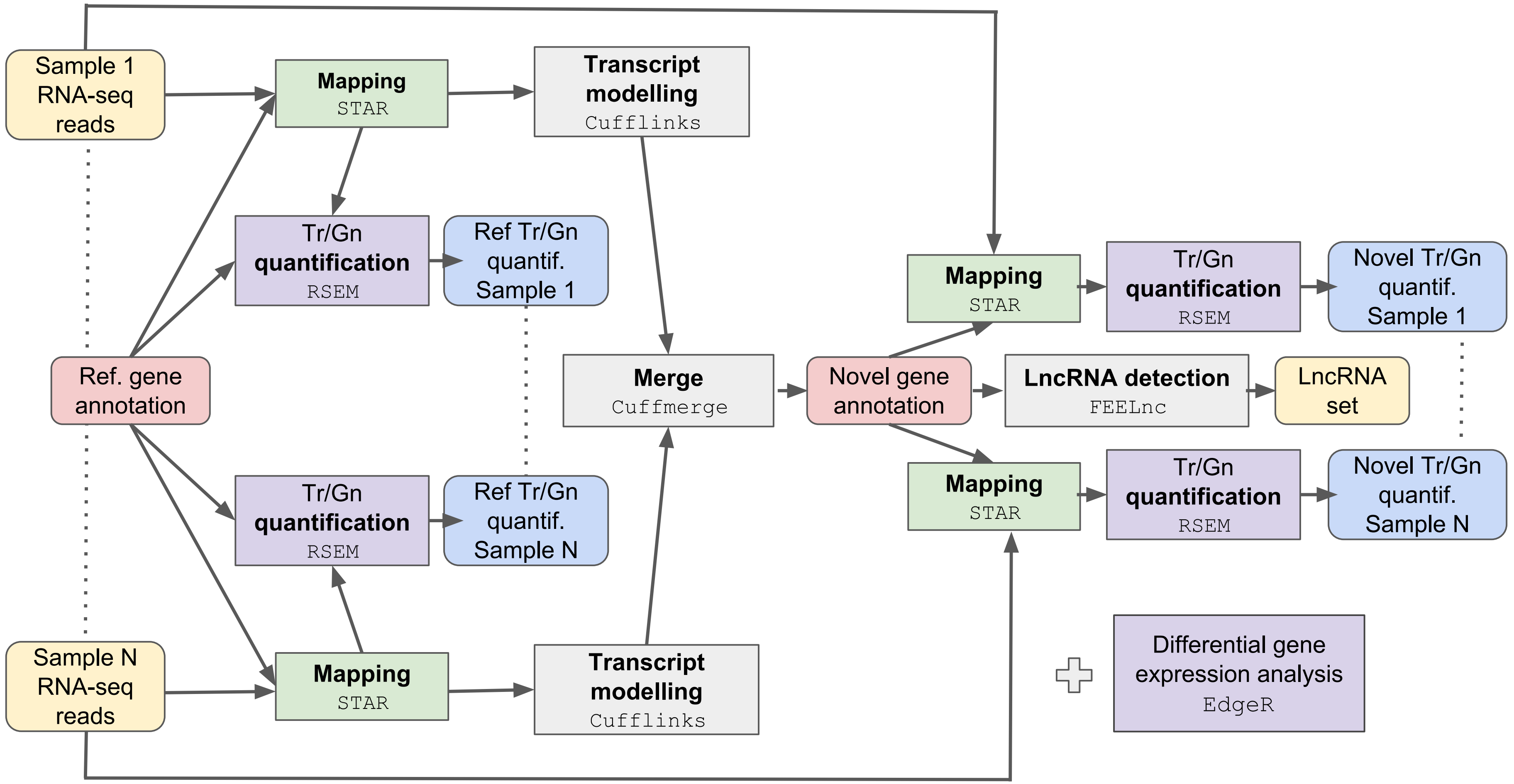


Figure S1: **RNA-seq processing pipeline overview.** RNA-seq reads were processed with STAR for mapping and Cufflinks for transcript modeling using the reference gene annotation as input and for each sample individually. Modelled transcripts were then merged with Cuffmerge to produce a single gene annotation per species, and the transcripts from this novel gene set were further processed with FEELnc to produce a reliable set of lncRNAs. Reference and novel transcripts and gene expressions were quantified in each sample using the STAR/RSEM pipeline. Differential expression analysis was performed on reference and novel genes with TPM $\geq$ 0.1 in at least 2 samples using the R/Bioconductor package **edgeR**.

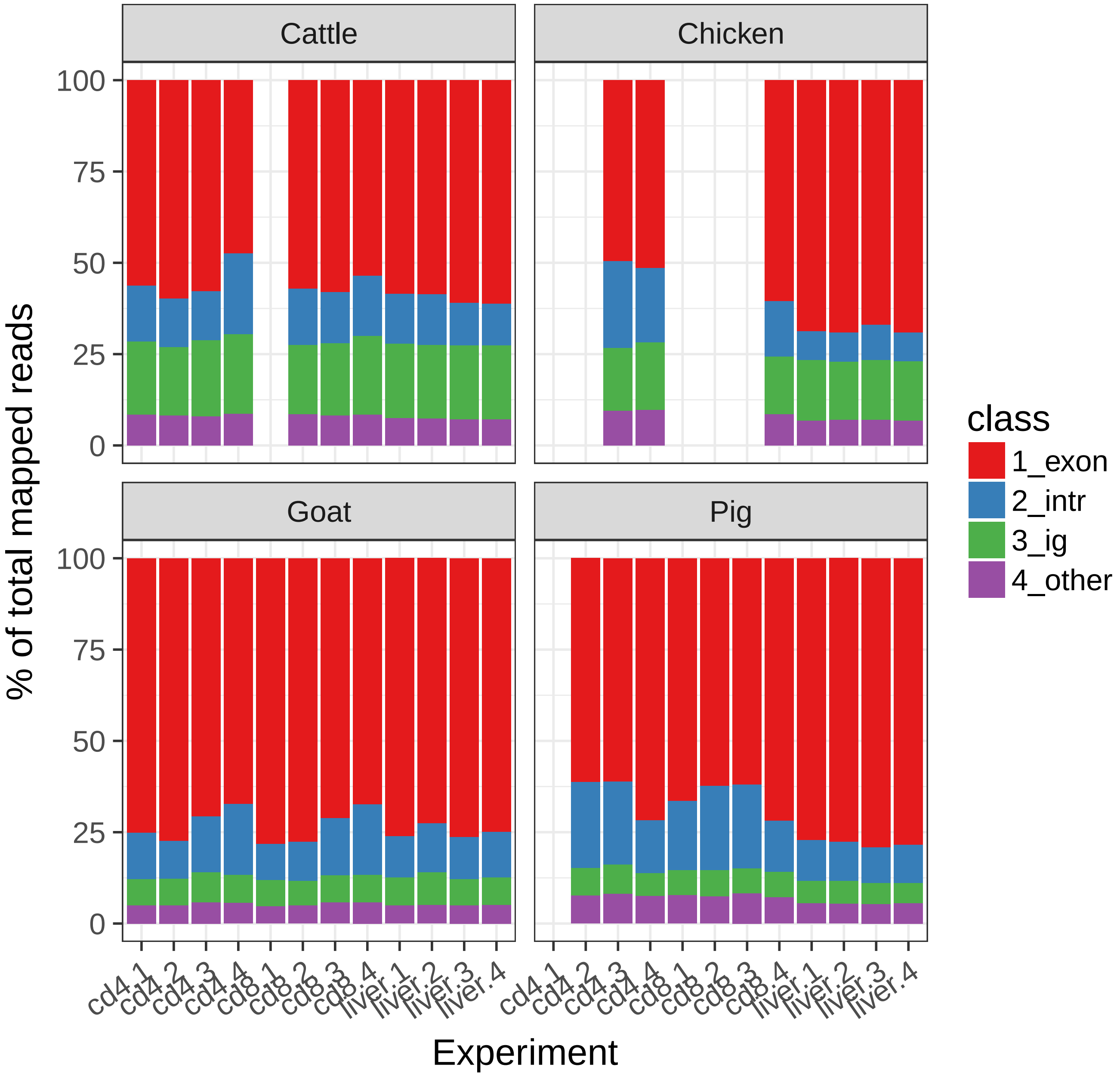


Figure S2: **RNA-seq mapped read classification into genomic domains.** For each sample, RNA-seq mapped reads were classified into exonic (1_exon), intronic (2_intr), intergenic (3_ig) and other (4_other) reads.

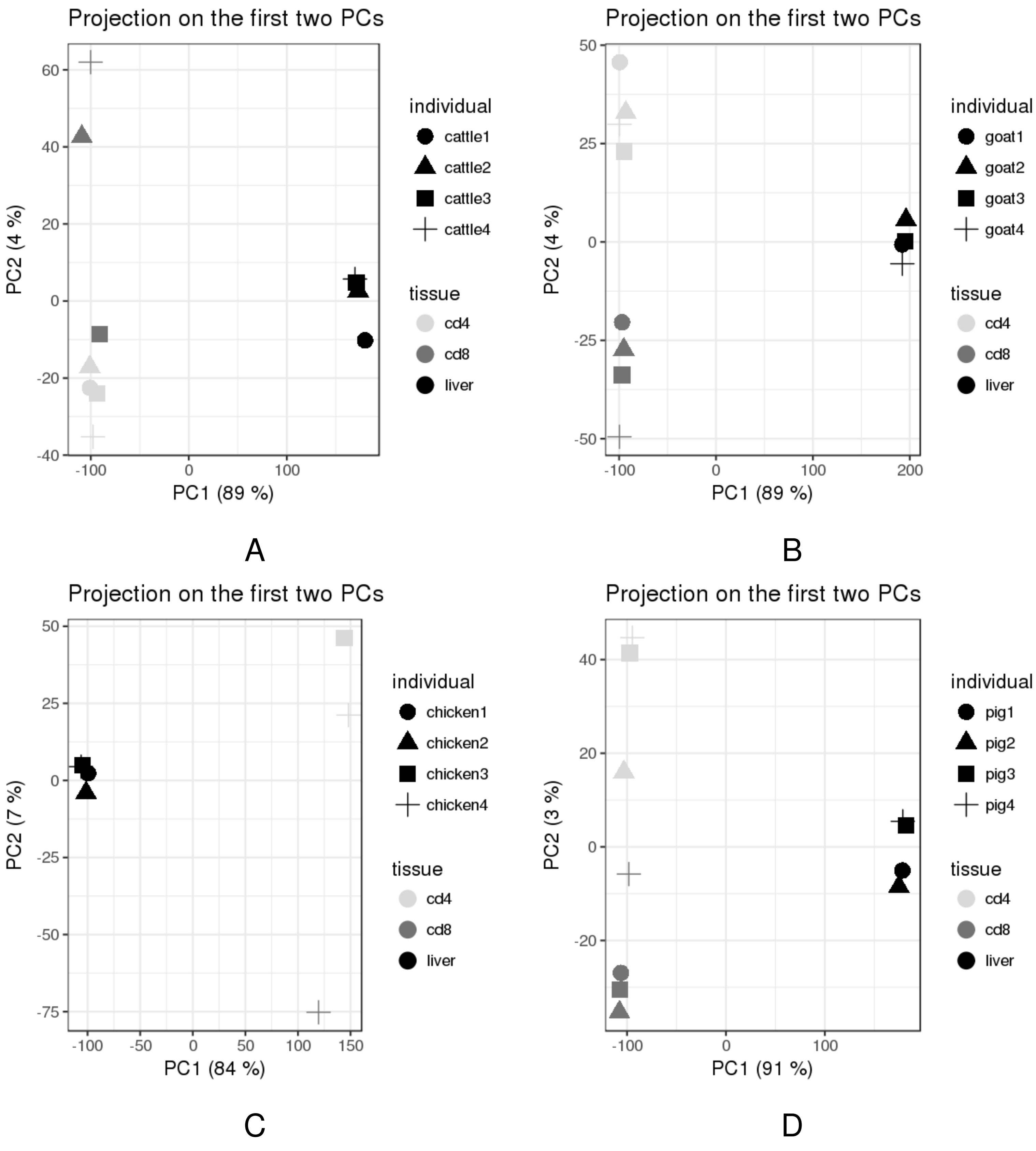


Figure S3: **RNA-seq sample PCA based on the expression of reference genes.** Only reference genes with a TPM $\geq 0.1$ in at least 2 samples were used.

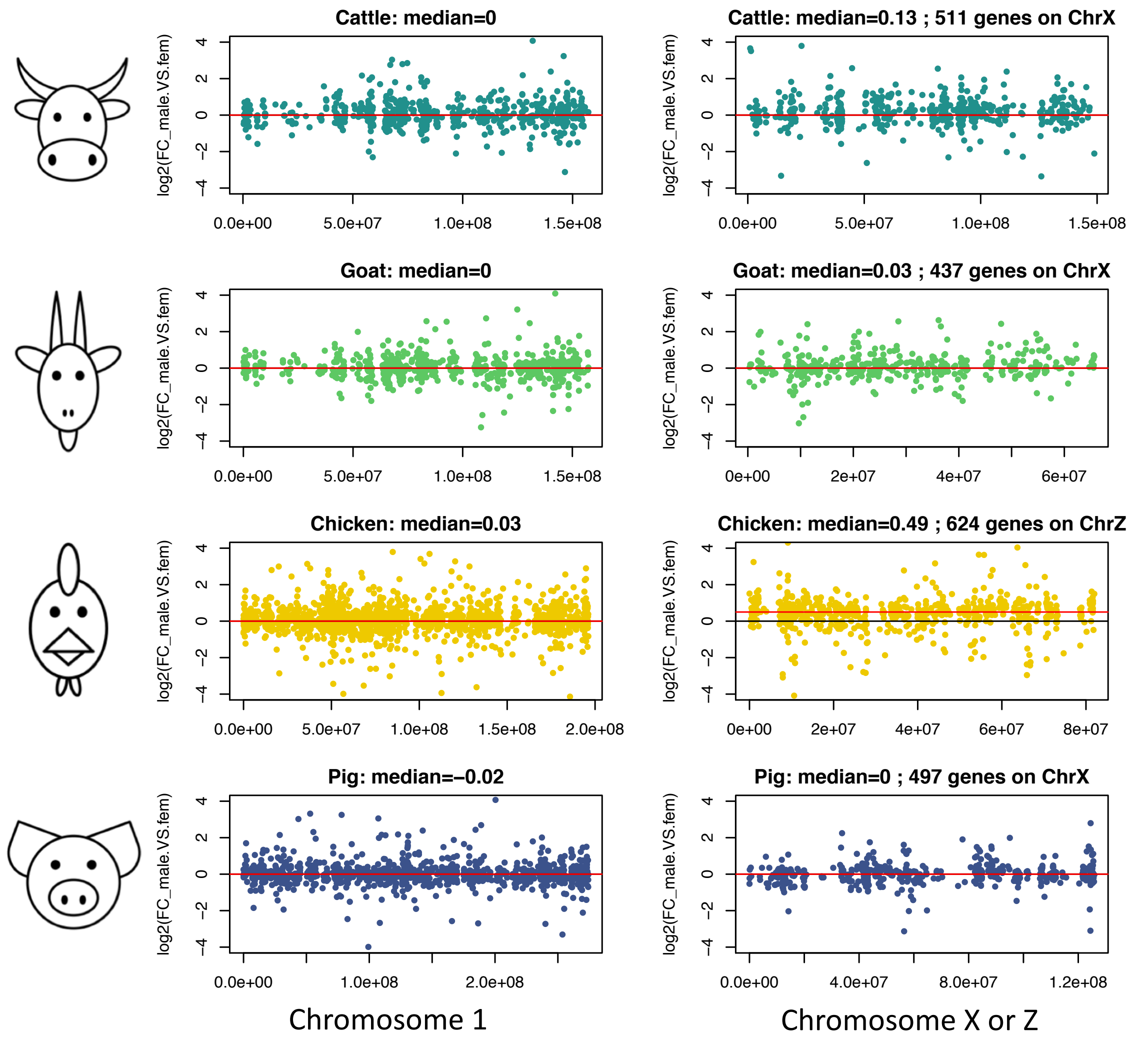


Figure S4: **Gene expression ratio in males versus females on chromosomes X or Z.** Gene expression was compared in males vs. females using the base 10 logarithm of the TPM values. $y$-axis: the $\log_2$ of the male/female expression ratio for the genes located on the chromosome of interest. For each plot, the median of these expression ratios is indicated in the title. Left: genes located on chromosome 1 used as a control. Right: genes located on chromosome X for mammals and Z for chicken. $x$-axis: the position of the gene on the chromosome of interest.

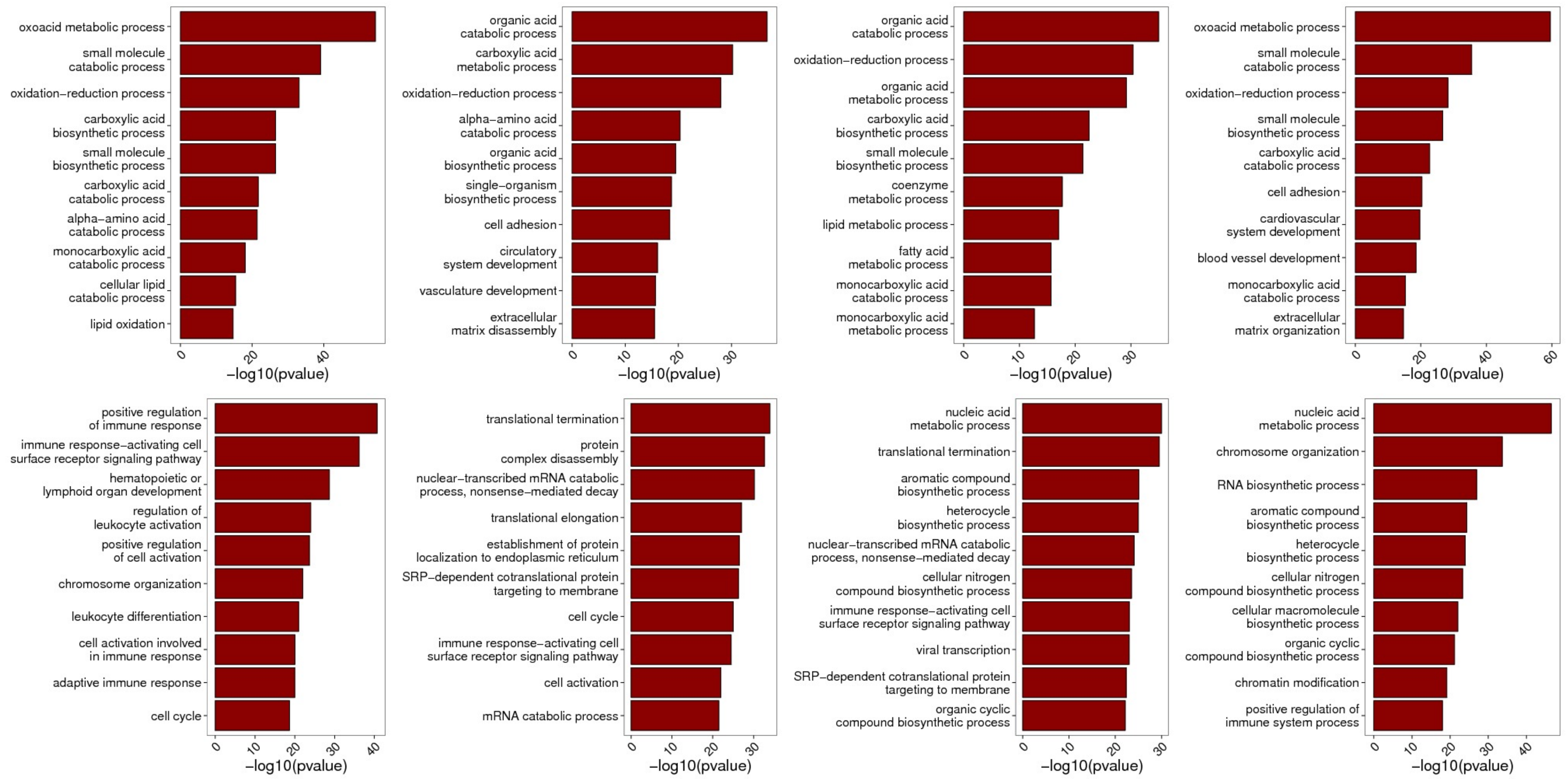


Figure S5: **Biological Process GO term enrichment analysis for reference genes differentially expressed between liver and T cells.** This analysis was performed for each species individually (in columns: cattle, goat, chicken, pig in that order) and for genes over-expressed in liver (top) and over-expressed in T cells (bottom).

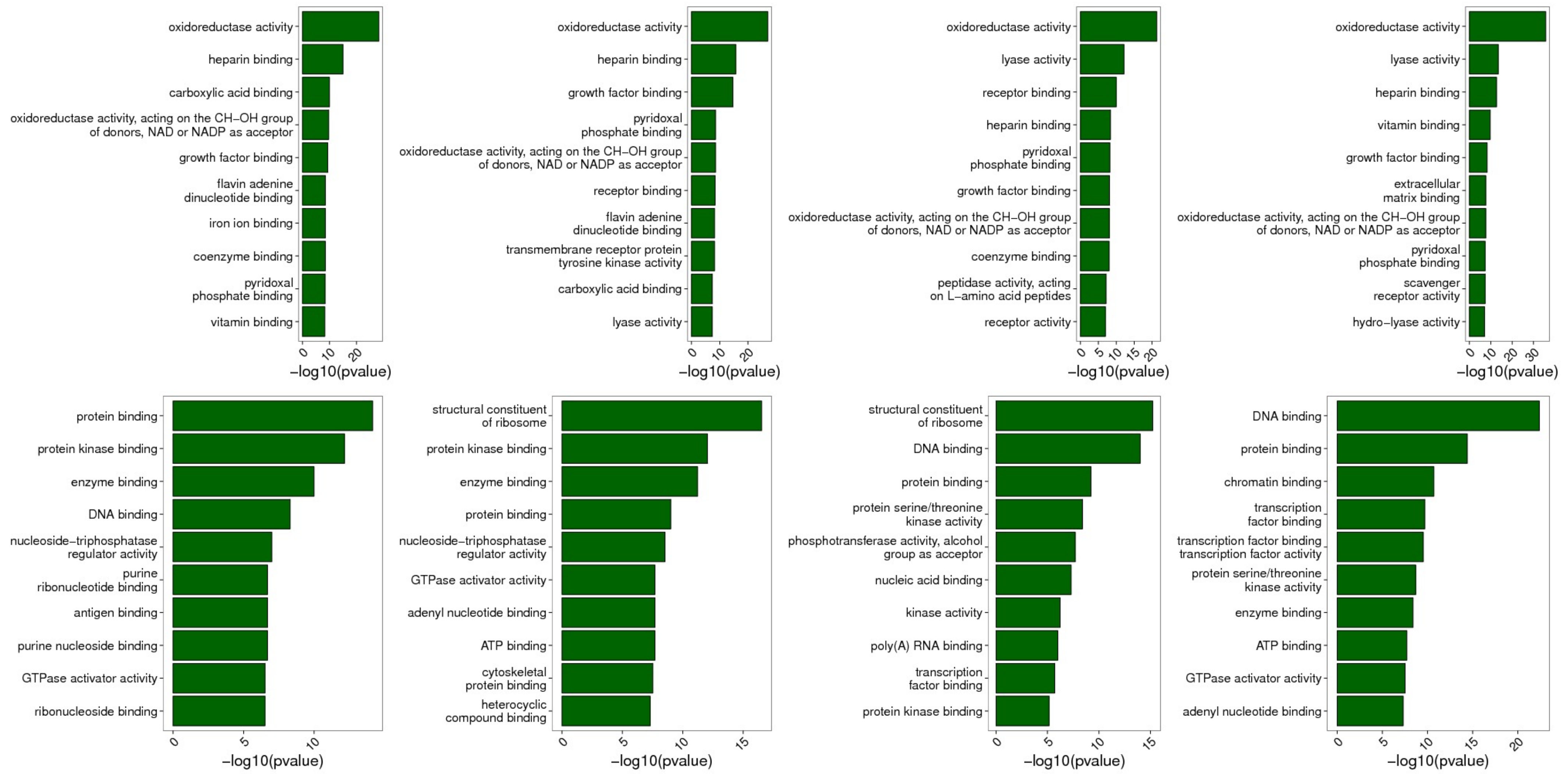


Figure S6: **Molecular Function GO term enrichment analysis for reference genes differentially expressed between liver and T cells.** This analysis was performed for each species individually (in columns: cattle, goat, chicken, pig in that order) and for genes over-expressed in liver (top) and over-expressed in T cells (bottom).

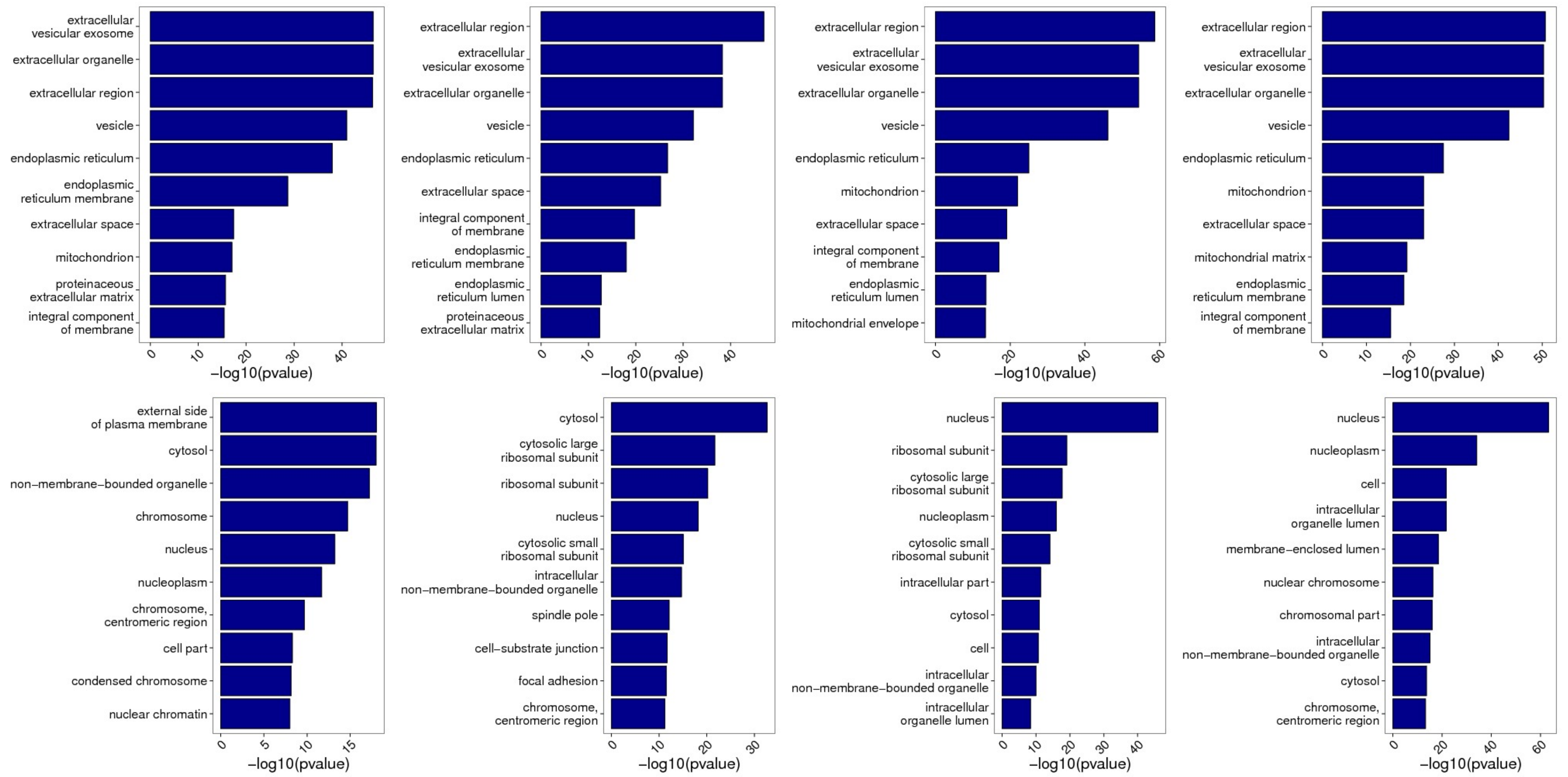


Figure S7: **Cellular Compartment GO term enrichment analysis for reference genes differentially expressed between liver and T cells.** This analysis was performed for each species individually (in columns: cattle, goat, chicken, pig in that order) and for genes over-expressed in liver (top) and over-expressed in T cells (bottom).

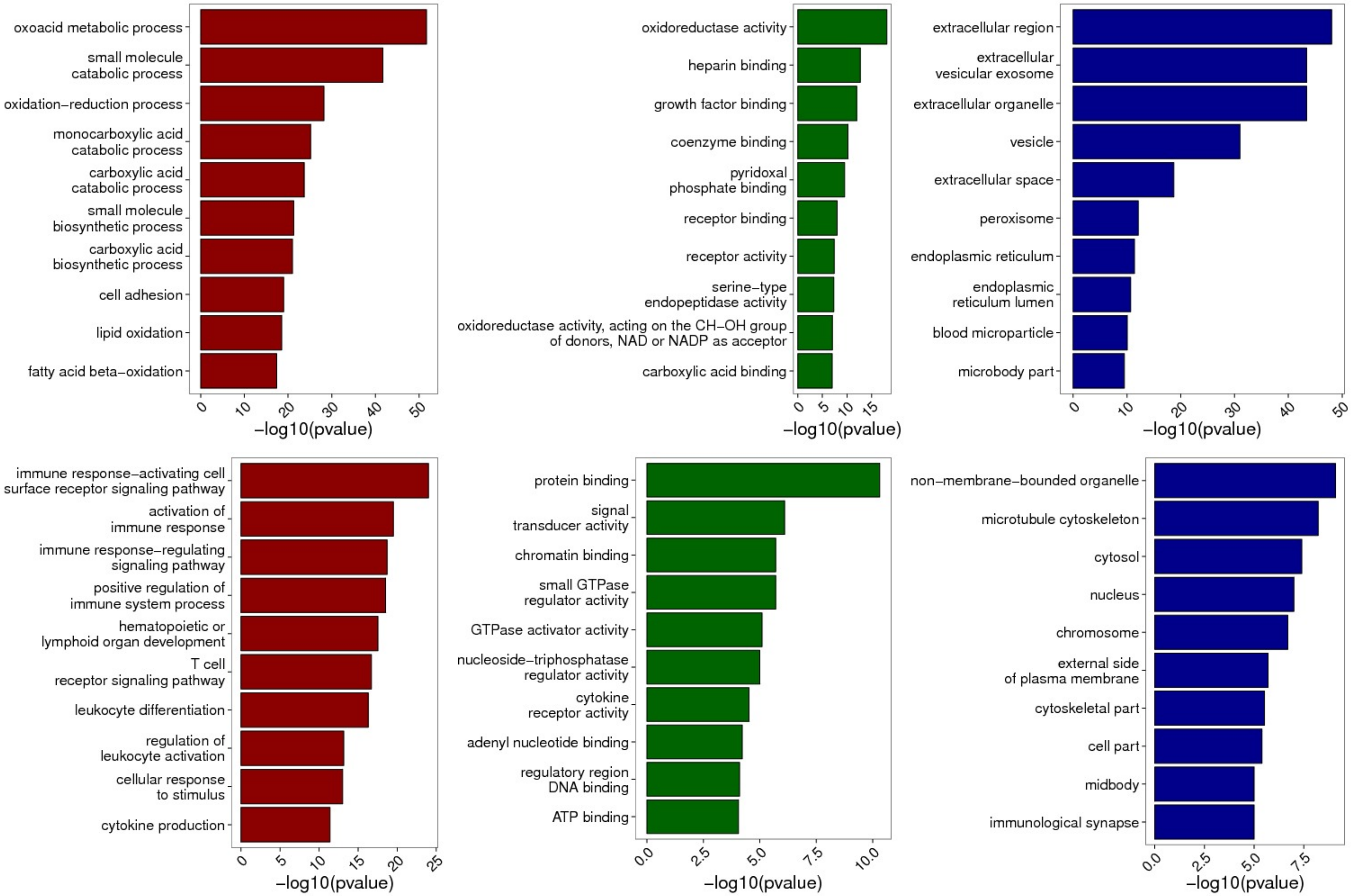


Figure S8: **GO term enrichment analysis for reference genes differentially expressed between liver and T cells in all species.** This analysis was performed for each GO (in column: Biological Process, Molecular Function, Cellular Compartment in that order) and for genes over-expressed in liver (top) and over-expressed in T cells (bottom).

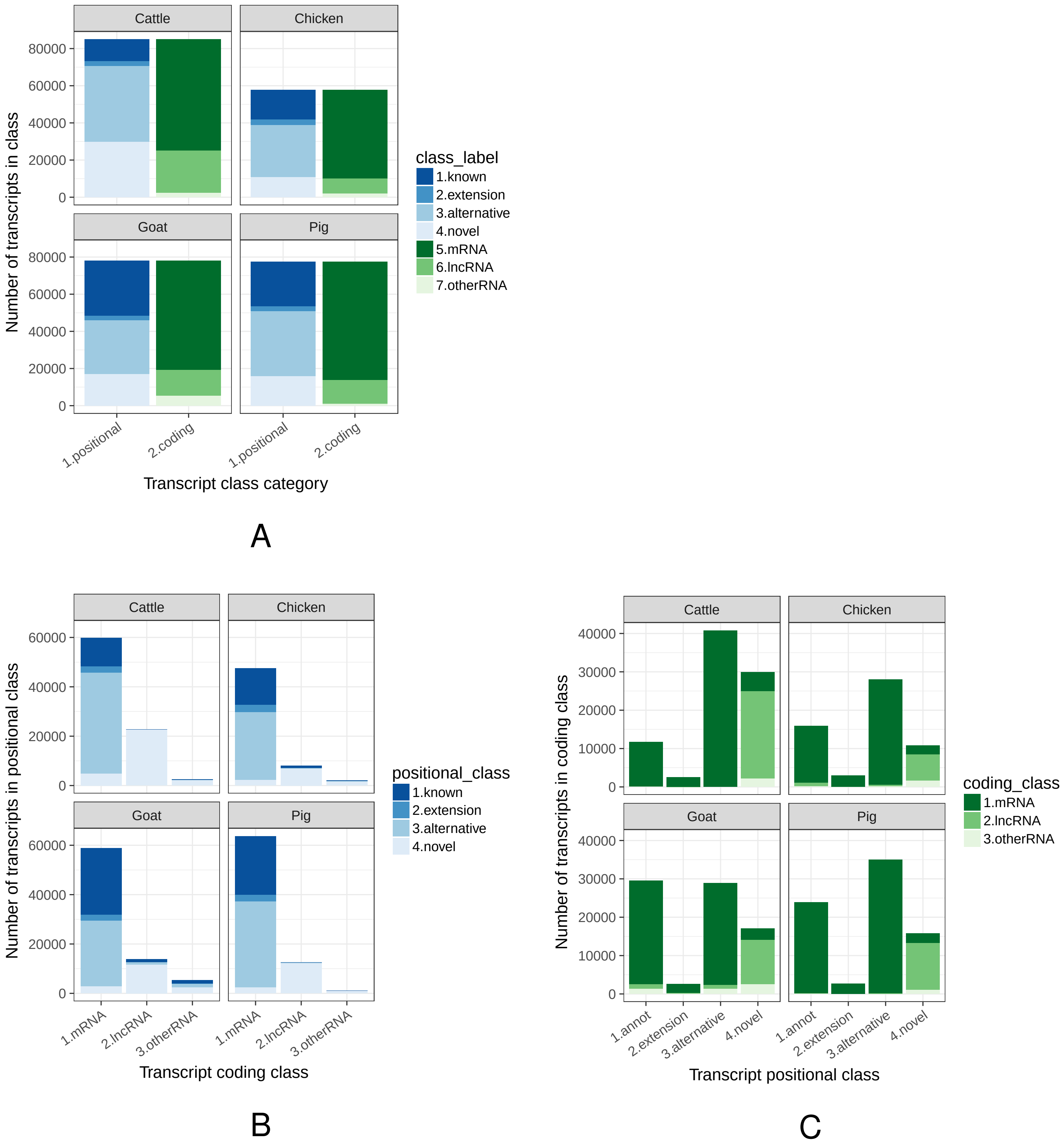


Figure S9: **Distribution of FR-AgENCODE transcripts into four positional (known, extension, alternative, novel) and three coding (mRNA, lncRNA, otherRNA) classes. (A)** Distribution of all FR-AgENCODE transcripts into positional and coding classes; **(B)** and **(C)** Distribution of FR-AgENCODE transcripts of each coding class into positional classes, and of each positional class into coding classes respectively.

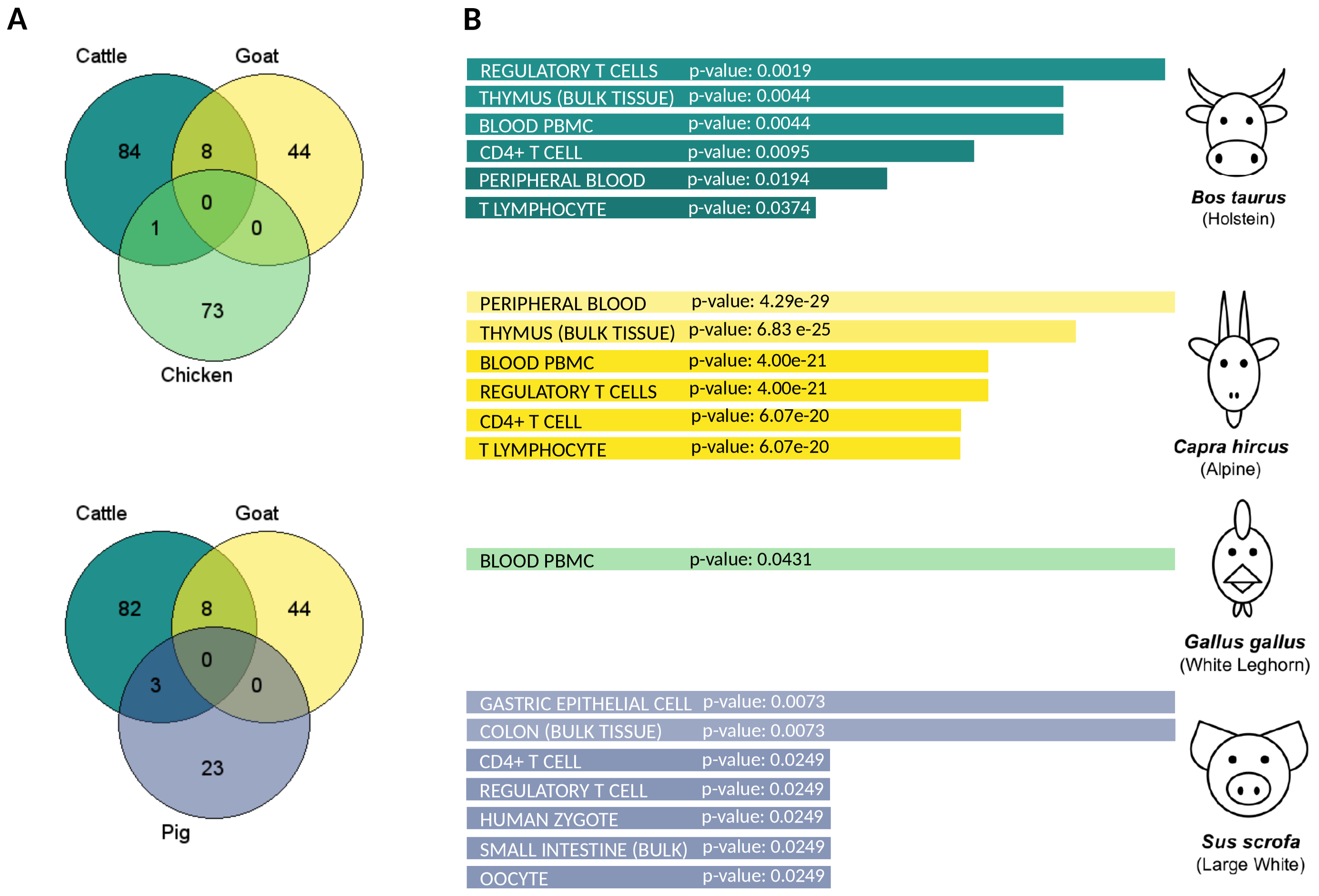


C

| Novel coding FR-AgENCODE genes orthologous to human and shared by at least 2 livestock species |
|---|
| BIK |
| GCNA |
| TRBV20OR9-2 |
| TRBV4-1 |
| TRBV27 |
| TRAV14DV4 |
| TRAV12-3 |
| TRAV17 |
| TRBV3-1 |
| TRBV25-1 |
| INAFM2 |
| FAM200B |

Figure S10: **Novel coding FR-AgENCODE genes enrich the set of blood and T cell annotated genes.** (A) Venn diagrams of novel coding genes for each triplet of livestock species (two livestock genes are defined as orthologous if they project to the same human gene). (B) Gene set enrichment analysis on the human orthologs of the 93 cattle, 52 goat, 74 chicken and 26 pig genes, using EnrichR (Kuleshov et al. 2016; `http://amp.pharm.mssm.edu/Enrichr/`) on the ARCS4 Tissue dataset (`https://amp.pharm.mssm.edu/archs4/`). (C) Gene names of the 12 genes that are common to two livestock species. Out of these 12, 8 are coding for T cell Receptor Alpha or Beta Variable genes.

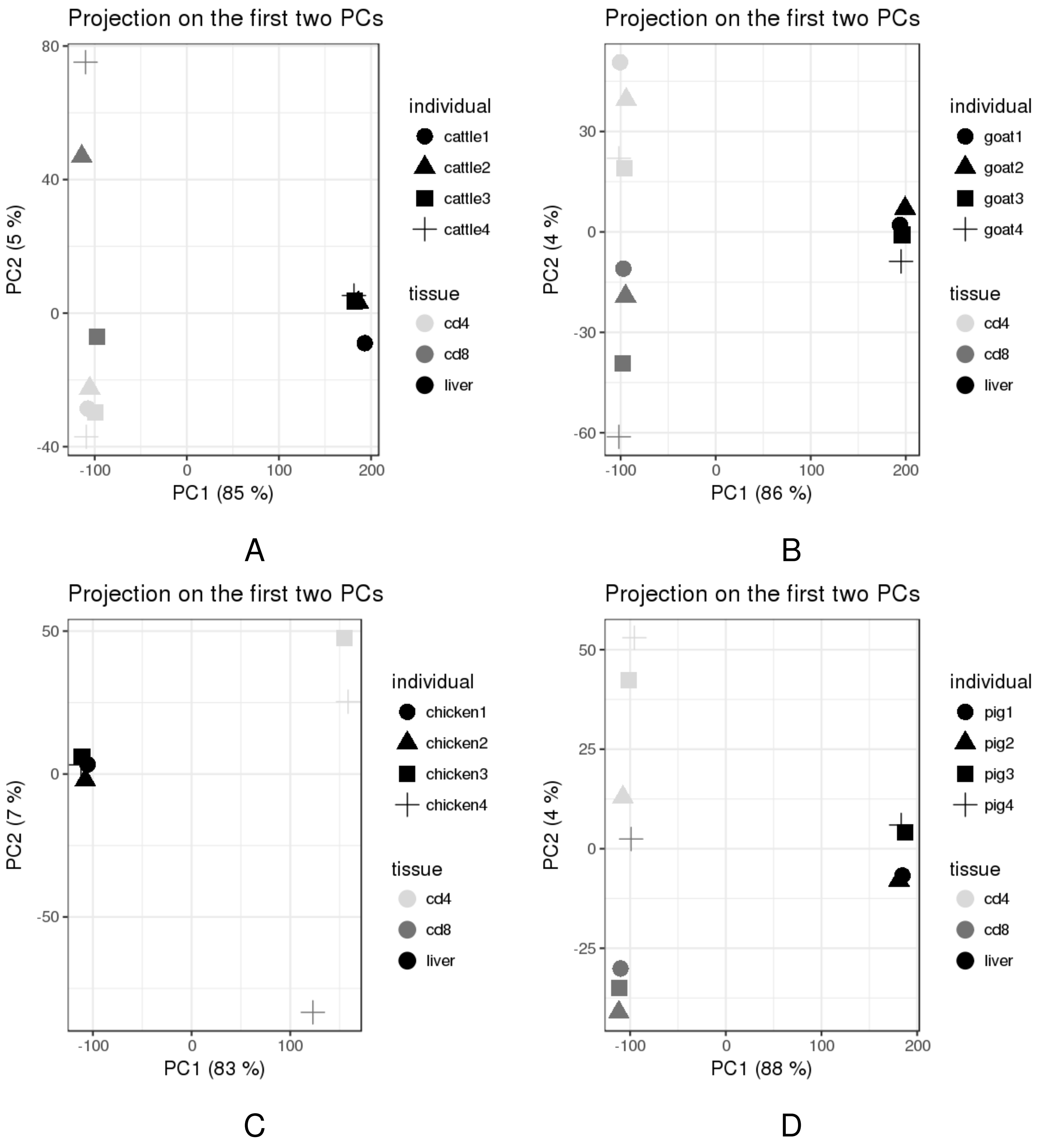


Figure S11: **RNA-seq sample PCA based on FR-AgENCODE gene expression.**

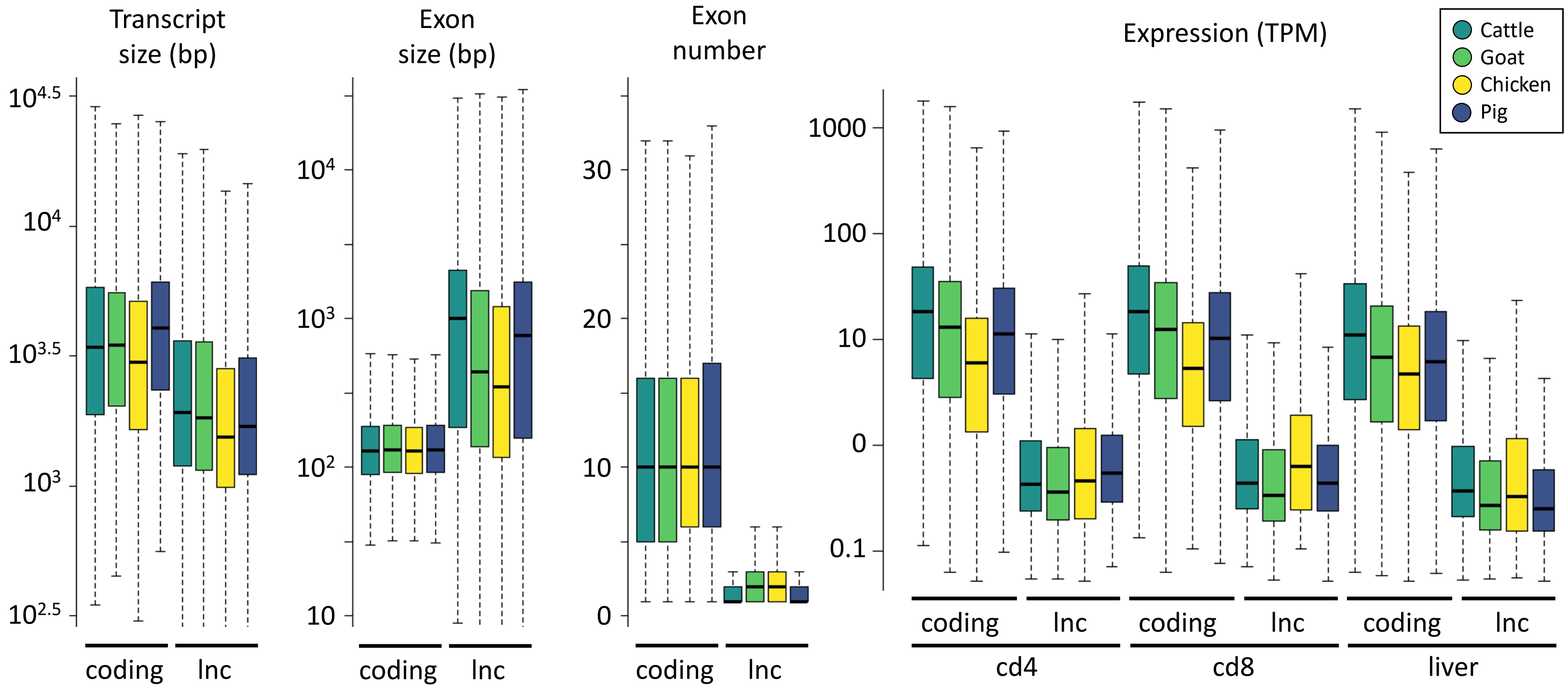


Figure S12: **LncRNA gene features.** Genomic structure features (left) and expression (right) are indicated for protein coding genes (coding) and lncRNA genes (lnc). The expression (TPM) is given for the three tissues and the four species; lncRNA genes are between 8- and 43-fold less expressed than protein coding genes (expression median: 0.4 vs. 10 TPM respectively).

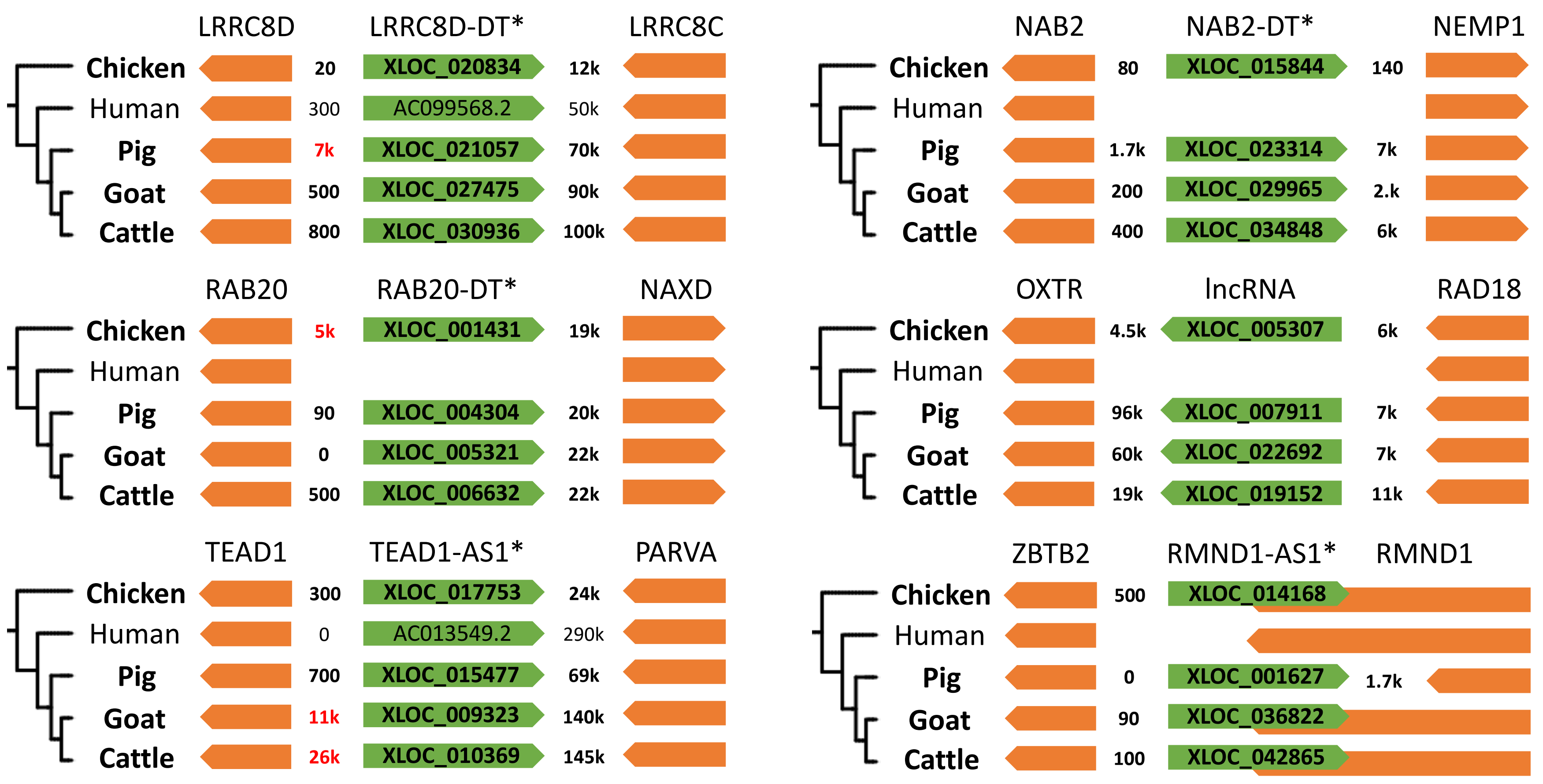


Figure S13: **Syntenic lncRNAs conserved between the 4 livestock species and human.** The 6 syntenic lncRNAs conserved in the 4 livestock species and human are represented in green and their surrounding protein coding genes in orange. Distances between the genes are indicated either in base pair or in kilo base pair (k). A distance of 0 means the genes are overlapping. A distance in red means a lower confidence in the orthology relationship for this species, after inspection of the distances found in the other species. When the lncRNA was not known before, a new gene name is proposed (asterisk).

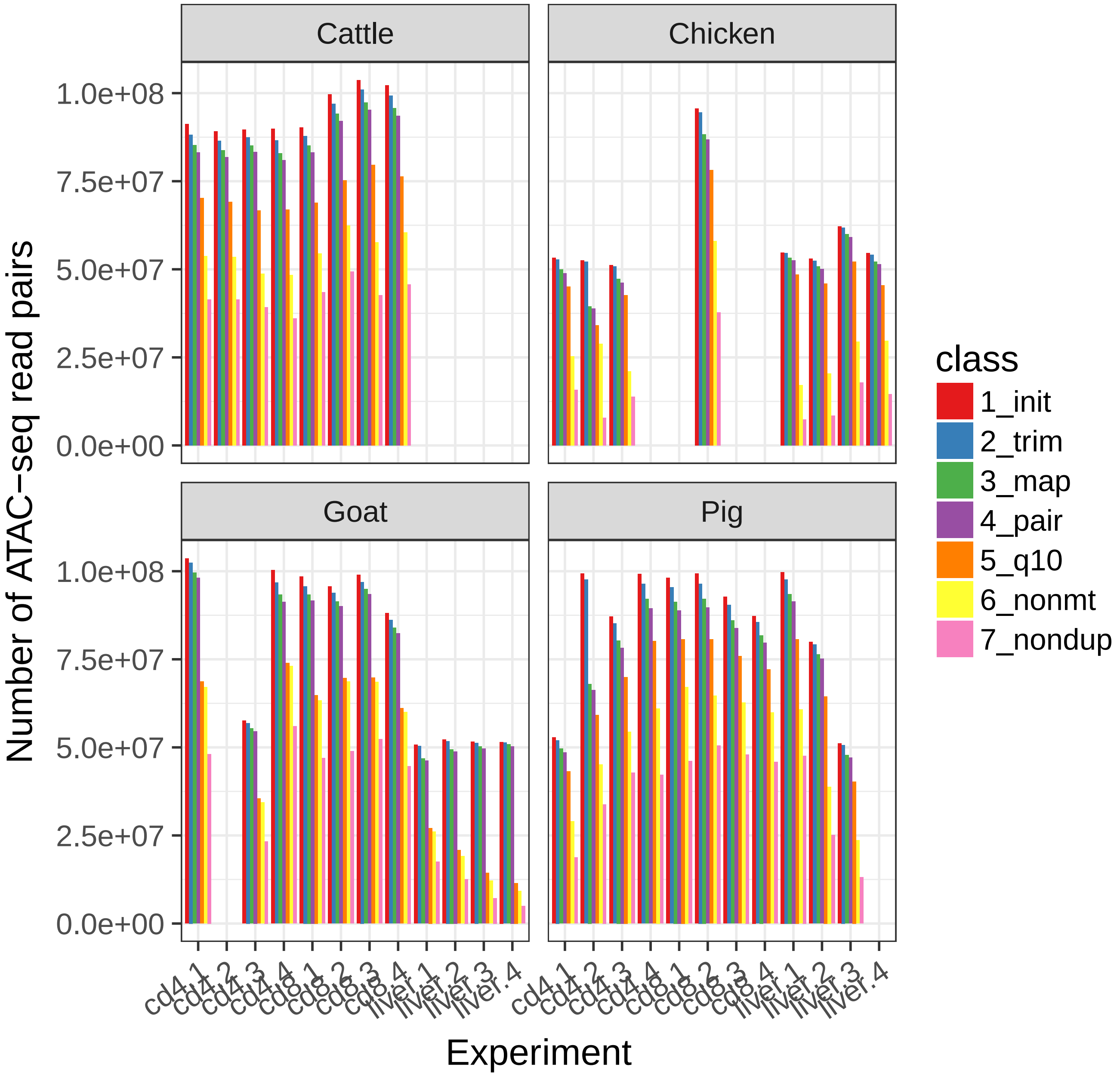


Figure S14: **ATAC-seq read pair summary statistics.** For each species and sample (labelled by its tissue and animal number), the number of initial read pairs (1_init), of read pairs obtained after trimming (2_trim), mapping (3_map), proper pairing (4_pair), q10 filtering (5_q10), mitochondrial read removal (6_nonmt), and PCR duplicate read removal (7_nondup) are shown.

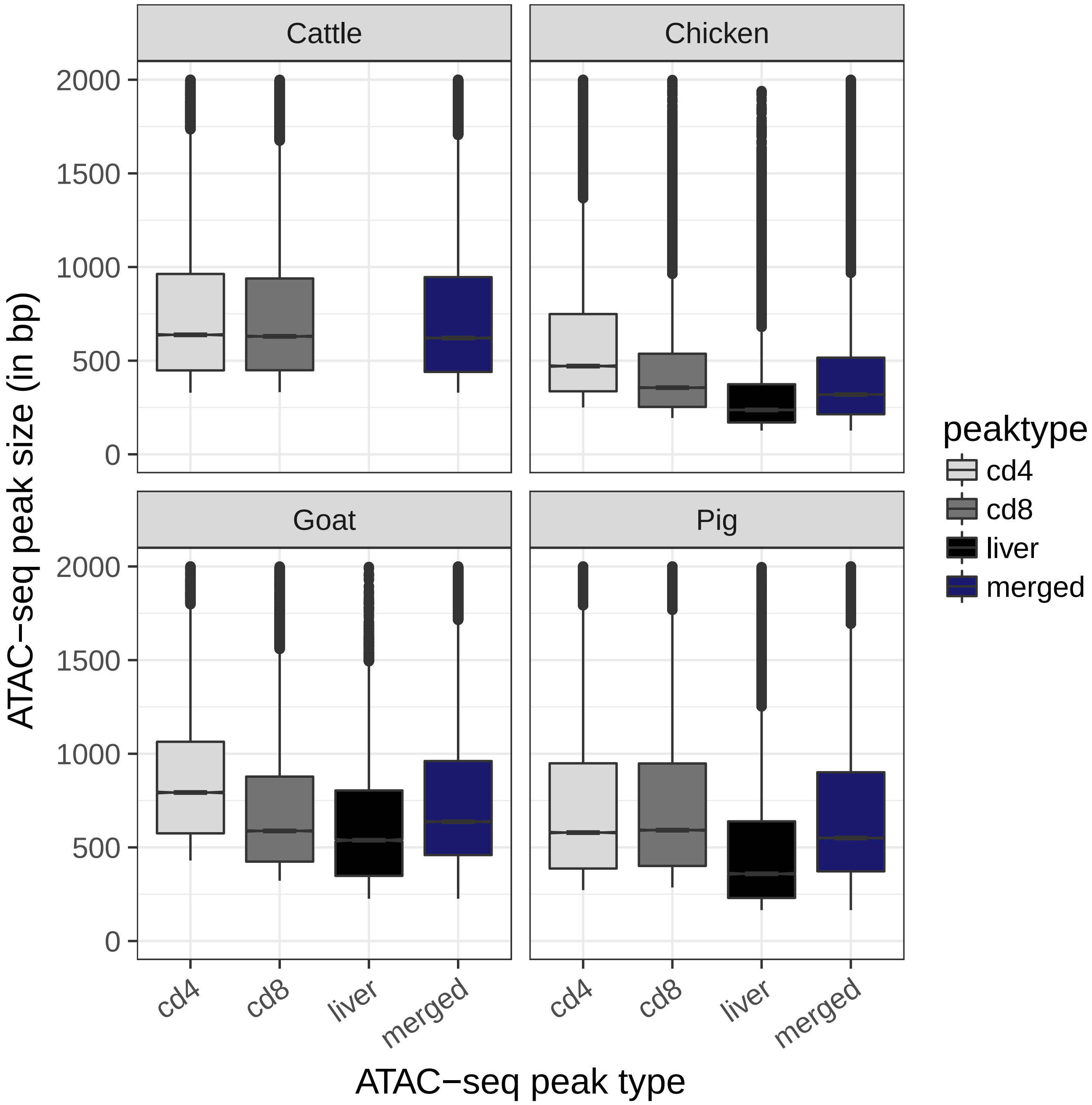


Figure S15: **ATAC-seq peak size distribution.** For each species, the size distribution of tissue and merged (i.e. across all tissues) ATAC-seq peaks are provided.

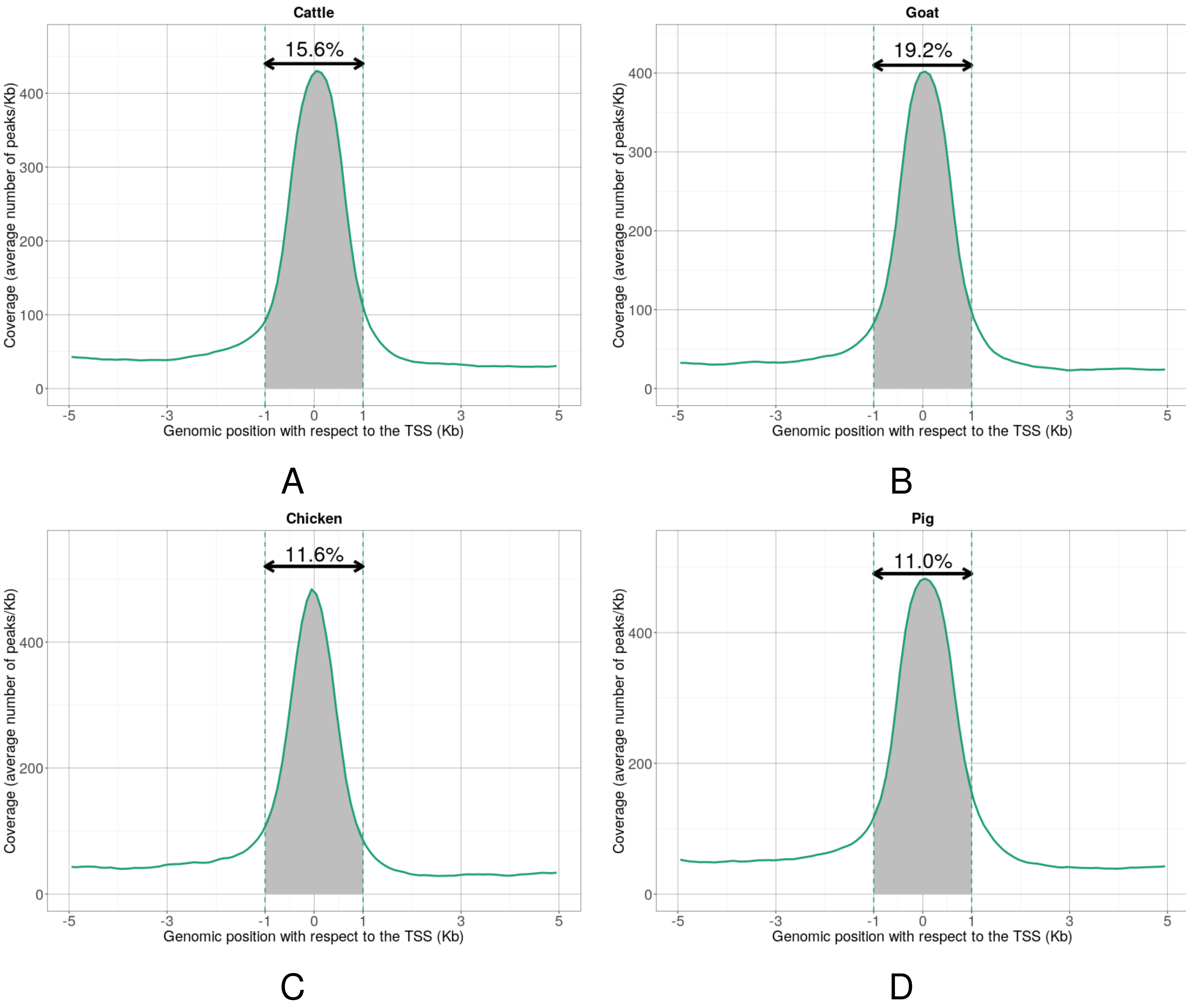


Figure S16: **Density of ATAC-seq peaks around starts of novel (i.e. not known) transcripts for cattle (A), goat (B), chicken (C) and pig (D)**

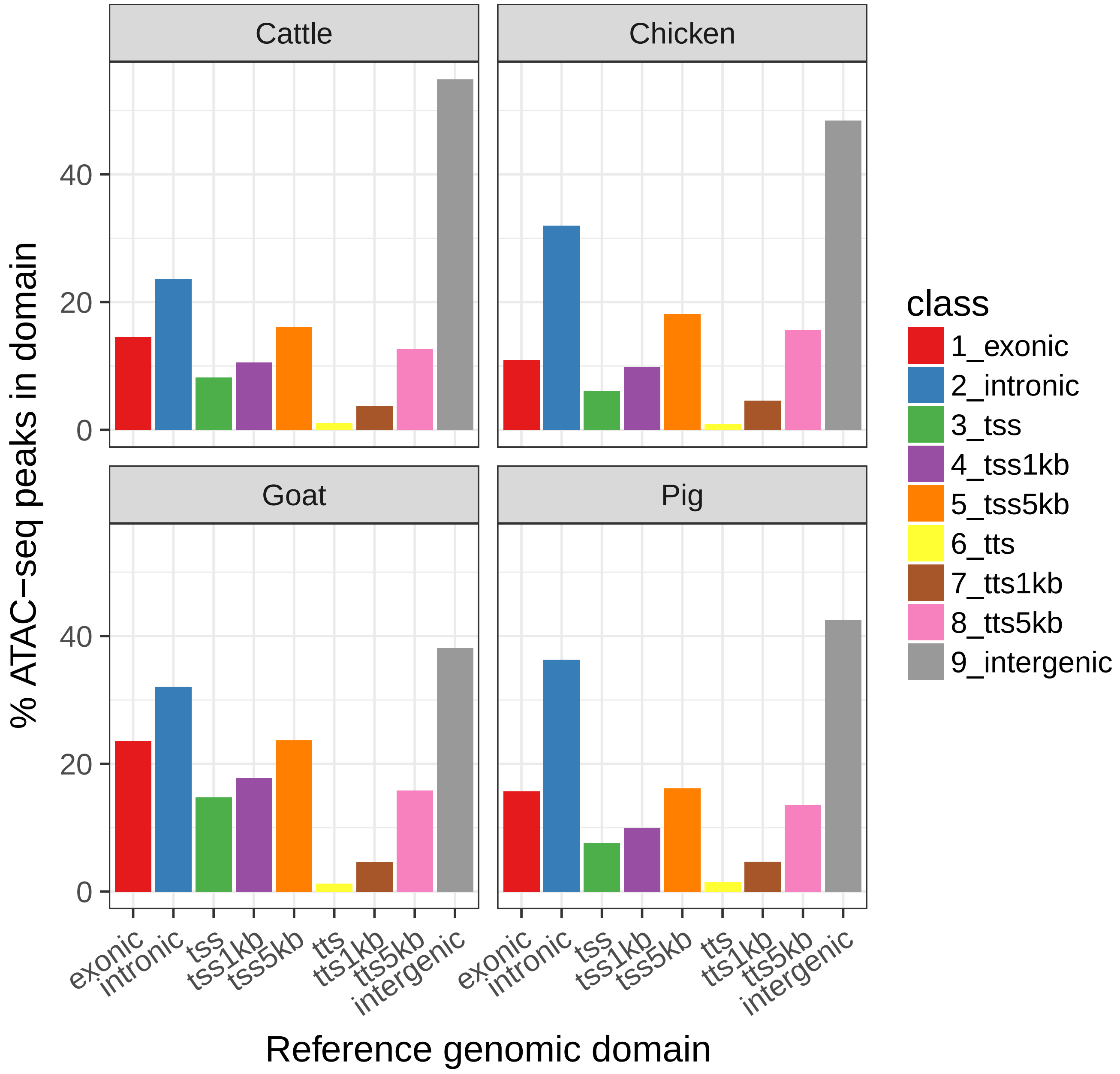


Figure S17: **ATAC-seq peak distribution into genomic domains.** For each species, are provided: the percentage of ATAC-seq peaks that are exonic, intronic, overlapping the TSS of a reference gene extended by 0, 1 or 5 Kb on each side, overlapping the TTS of a reference gene extended by 0, 1 or 5 Kb on each side and intergenic.

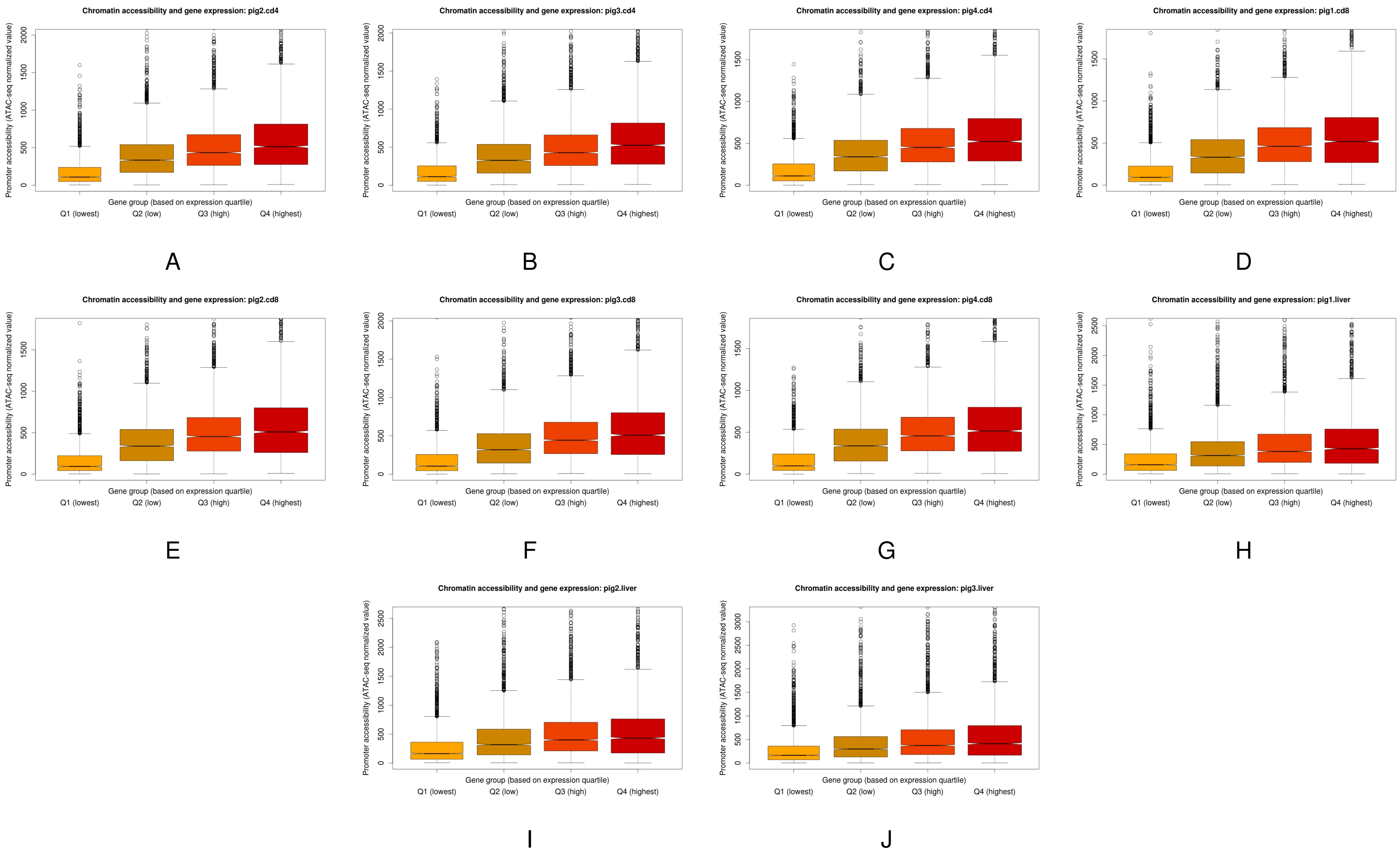


Figure S18: **Per sample promoter accessibility for four reference gene expression quartiles in pig.**

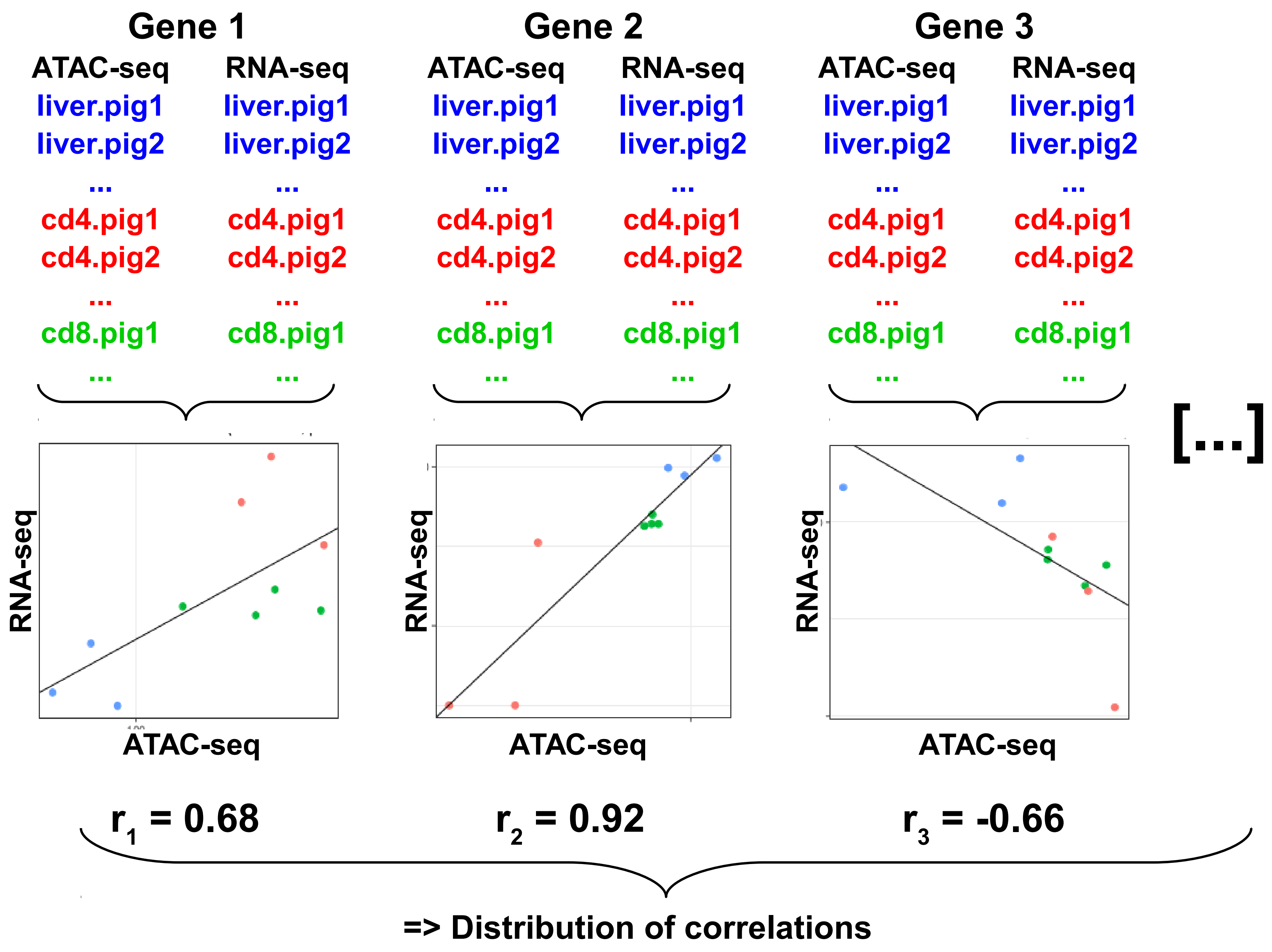


Figure S19: **Schematic illustration of the per-gene and cross-sample RNA-seq versus ATAC-seq correlation analysis.**

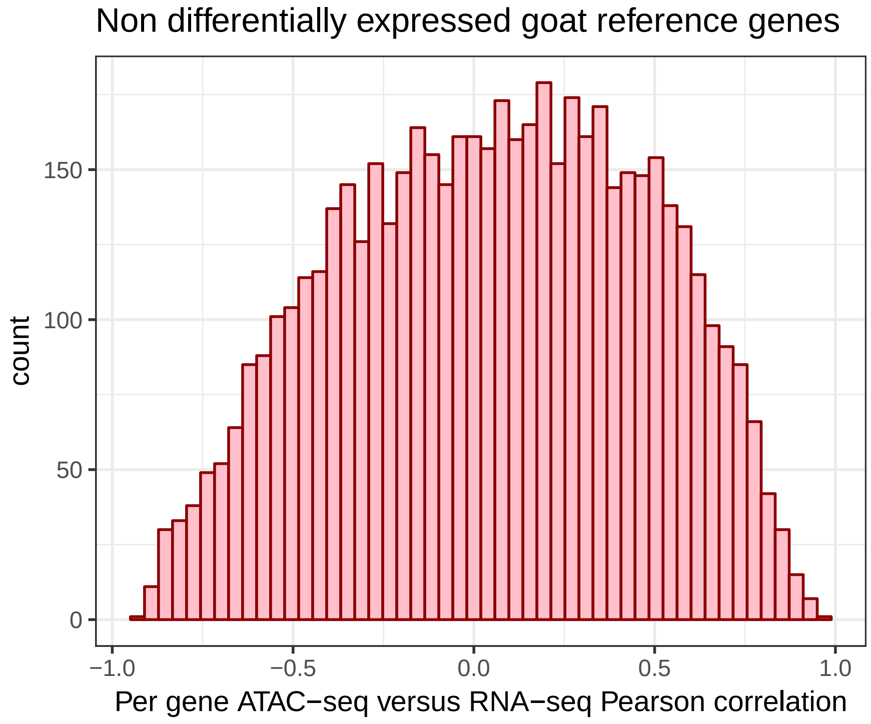


A

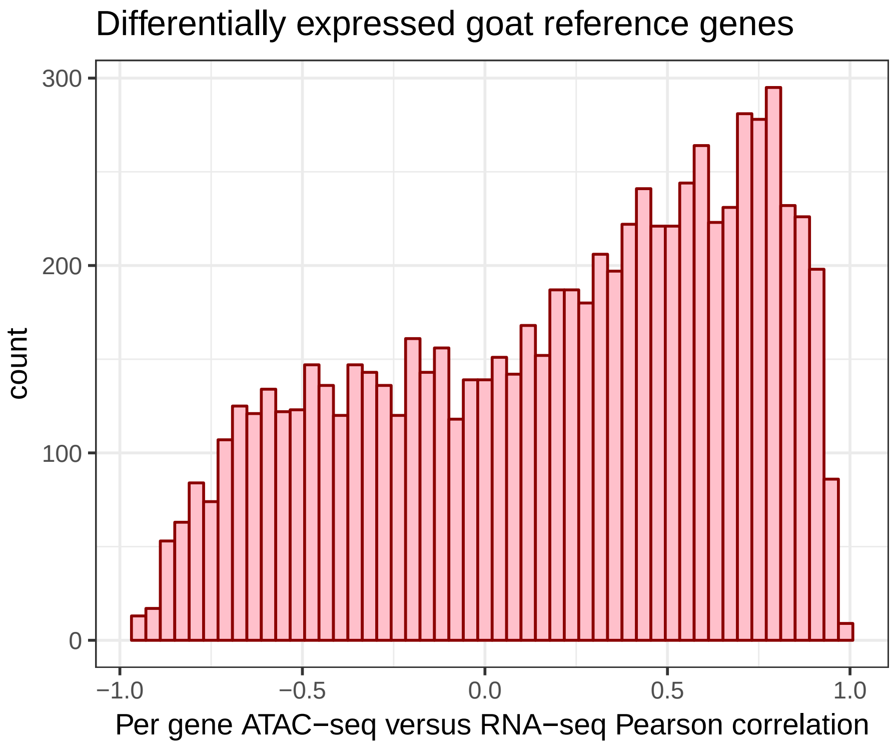


B

Figure S20: **Correlation between reference gene expression and chromatin accessibility at the TSS for goat.** For each pair of reference gene and ATAC-seq peak overlapping the reference gene TSS extended by 1 Kb on each side, the Pearson correlation was computed between the base 10 logarithm of the gene TMM and the base 10 logarithm of the normalized ATAC-seq signal at the peak. The distribution was then plotted for non DE genes **(A, top)** and for DE genes **(B, bottom)**.

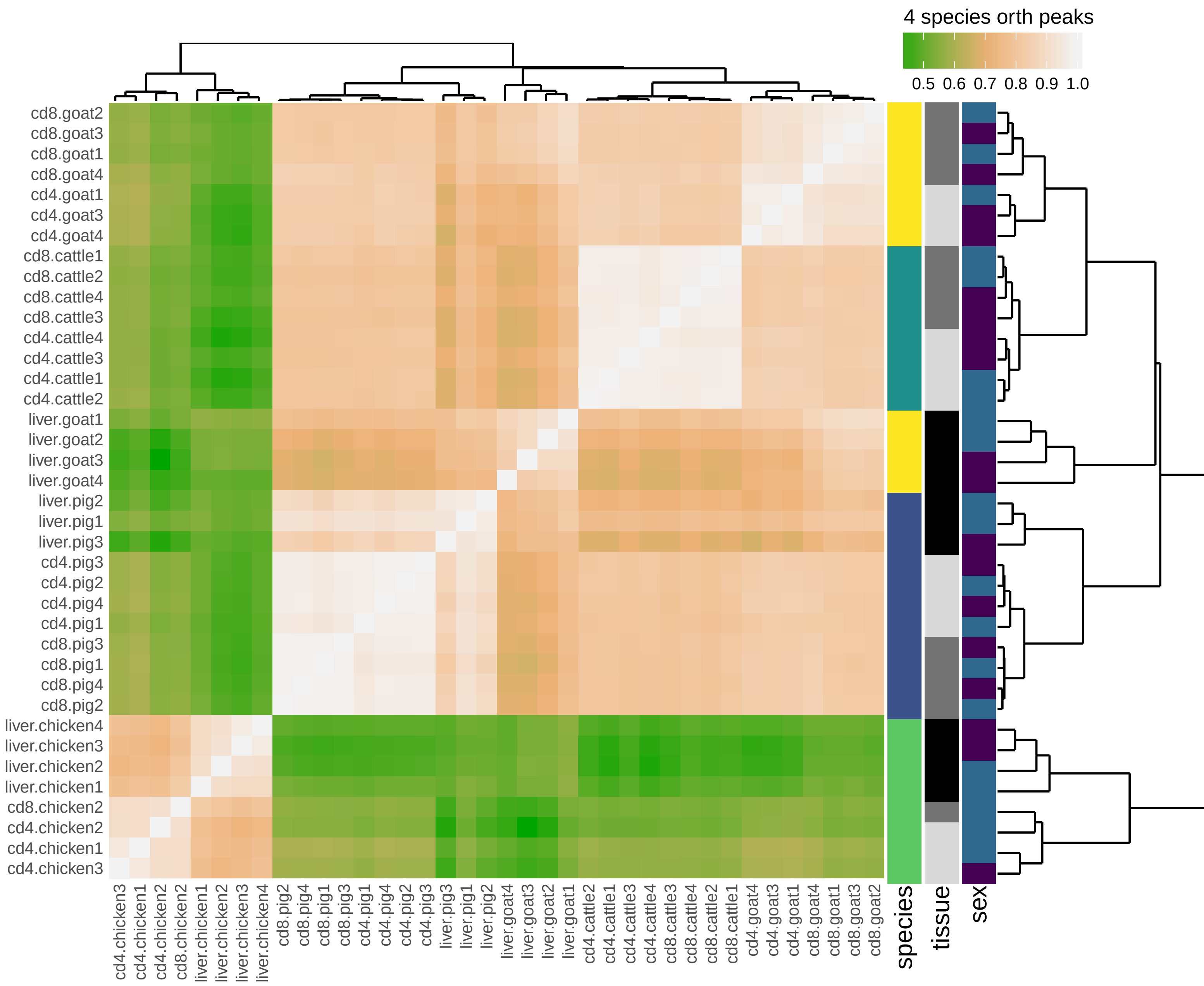


Figure S21: **ATAC-seq sample heatmap and hierarchical clustering based on the 1083 level 4 ATAC-seq peaks.** Pairwise similarity between samples is computed as the Pearson correlation between the base 10 logarithm of the normalized reads of the 1083 ATAC-seq peaks common to 4 species. These similarities are plotted as a heatmap, where samples appear both as rows and columns and are labelled by their species, tissue and the sex of the animal. The color of each heatmap cell also reflects the similarity (Pearson correlation) between each sample pair (the lighter, the higher). Hierarchical clustering is performed using one minus the squared Pearson correlation as a distance and the complete linkage aggregation method.

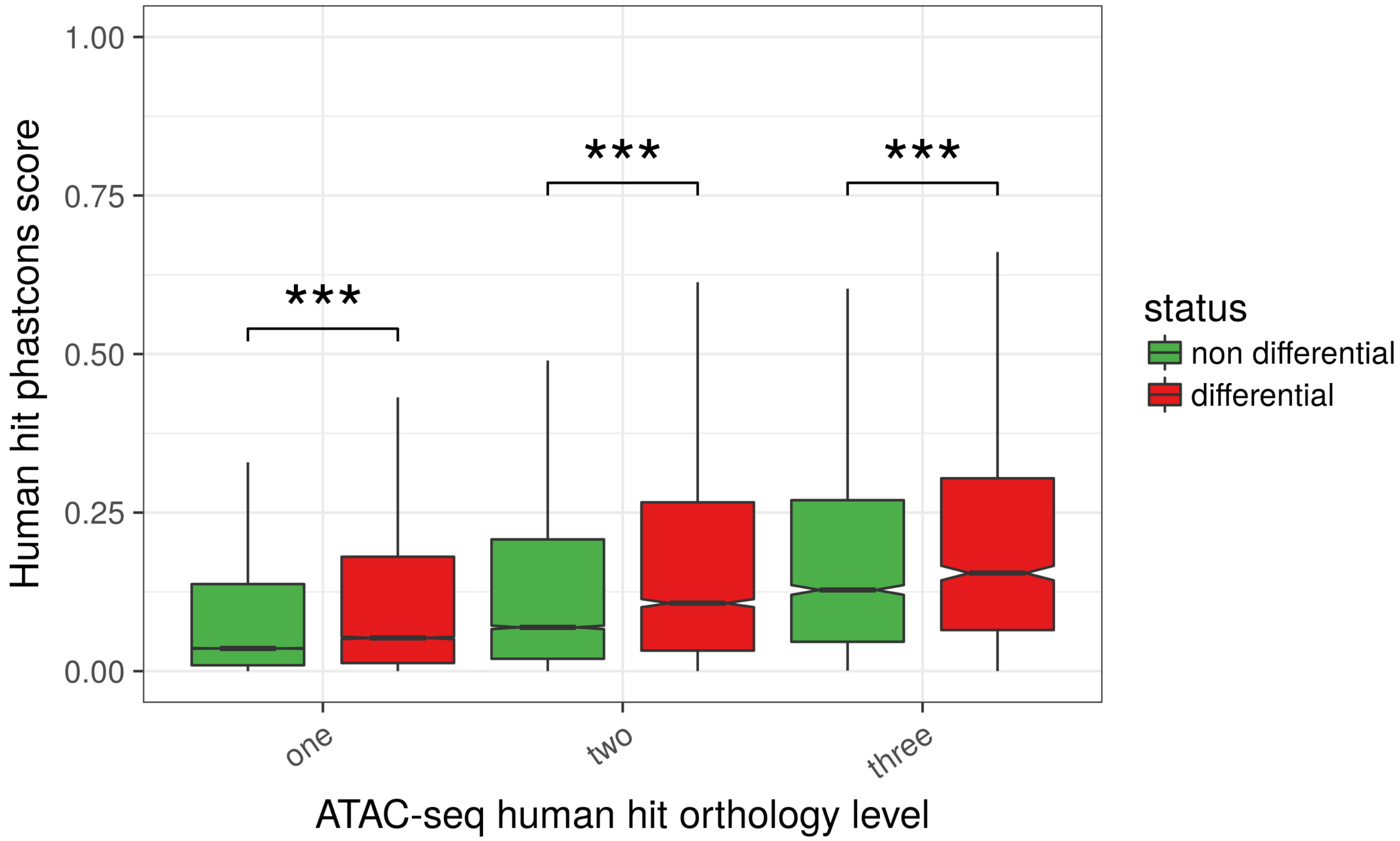


Figure S22: **Relation between non promoter proximal chromatin accessibility conservation and differential accessibility.** This figure is the same as main Figure 6 but is done restricting to ATAC-seq peaks not overlapping the TSS+-1kb in any of the species where it is presen. Phastcons scores of these ATAC-seq peaks were plotted after dividing the human hits according to both their orthology level (between 1 and 3, x-axis, level 4 removed because there were less than 50 peaks in a category) and their differential accessibility (DA) status (DA in at least one species or DA in none of the 4 species, boxplot color). Although the phastcons score obviously increases with the orthology level, it is clear that, for a given orthology level, the phastcons score is higher for DA human hits than for non DA human hits (all orthology levels, p-values $< 1.2e-6$ overall, Wilcoxon tests) (number of elements the boxplots from left to right: 151,783, 19,732, 11,922, 3,317, 2,100, 1,082).

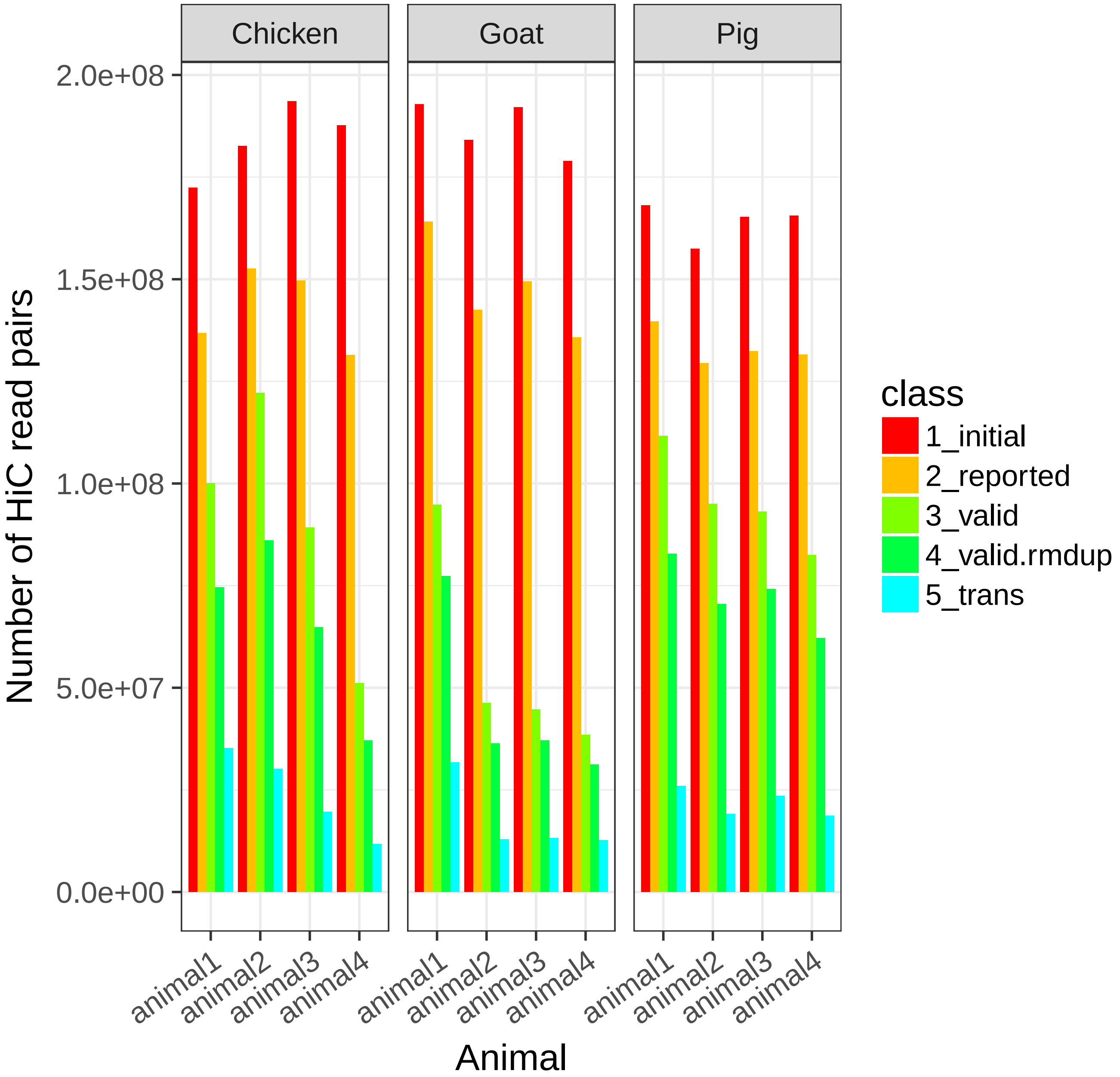


Figure S23: **Hi-C read summary statistics.** For each species and animal, are plotted the number of: initial read pairs after sequencing (1_initial), pairs with both reads mapped on the genome (2_reported), pairs in a valid configuration, that is when the sum of the distances from the reads to their next HindIII restriction sites downstream is comprised between 20bp and 1Kb (3_valid), pairs after removing PCR duplicates (4_valid_rmdup) and the number of read pairs supporting proximity between two different chromosomes (5_trans).

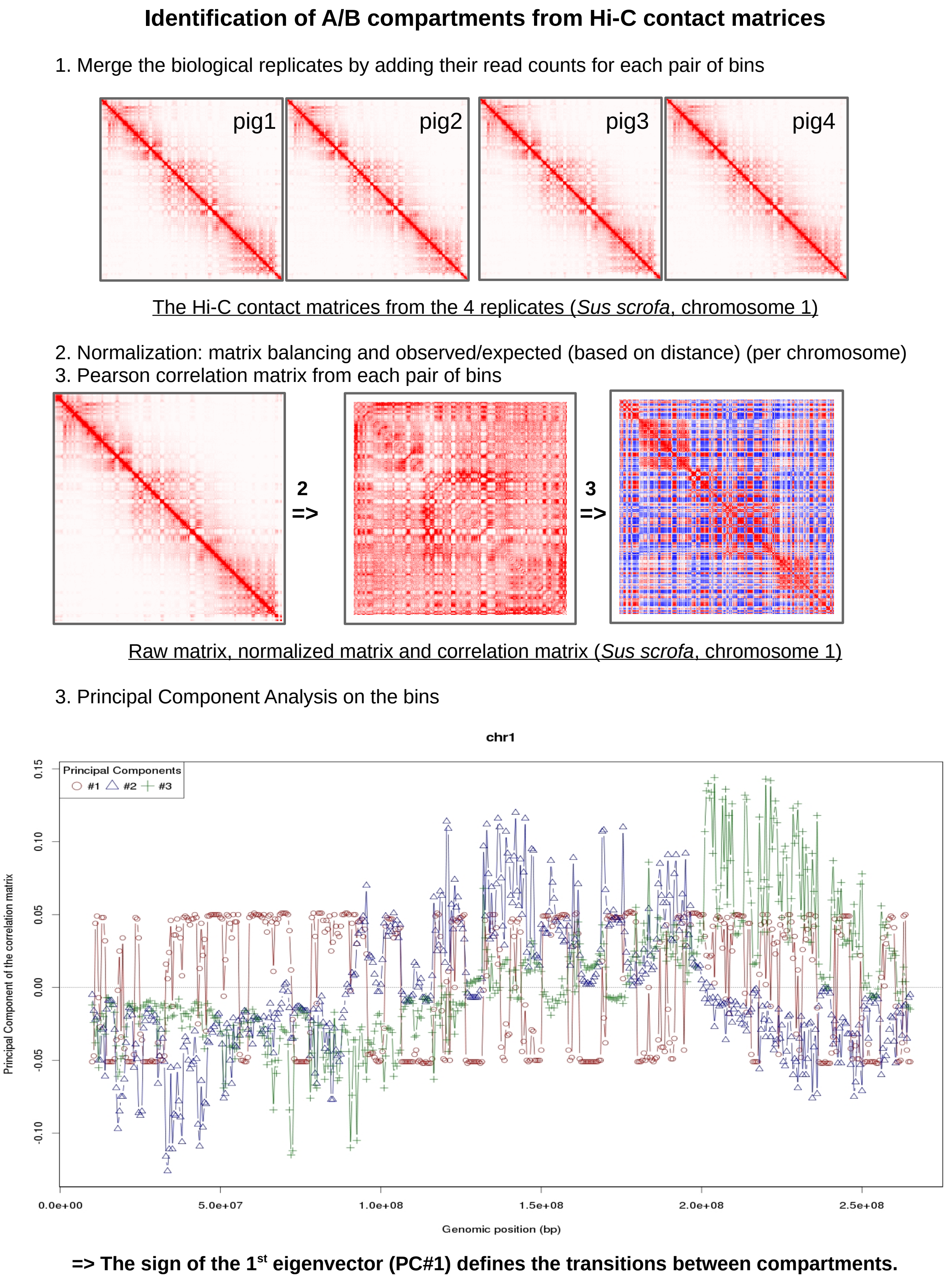


Figure S24: **Method for predicting Hi-C A and B compartments.** Illustration of the A/B compartment calling workflow using the interaction matrices of the first chromosome in pig. Upper panels: interaction matrices at different steps of the workflow. Lower panel: the first three eigenvectors are shown along the chromosome to illustrate the relevance of PC#1 as the discriminative value to segregate bins between A and B compartments.

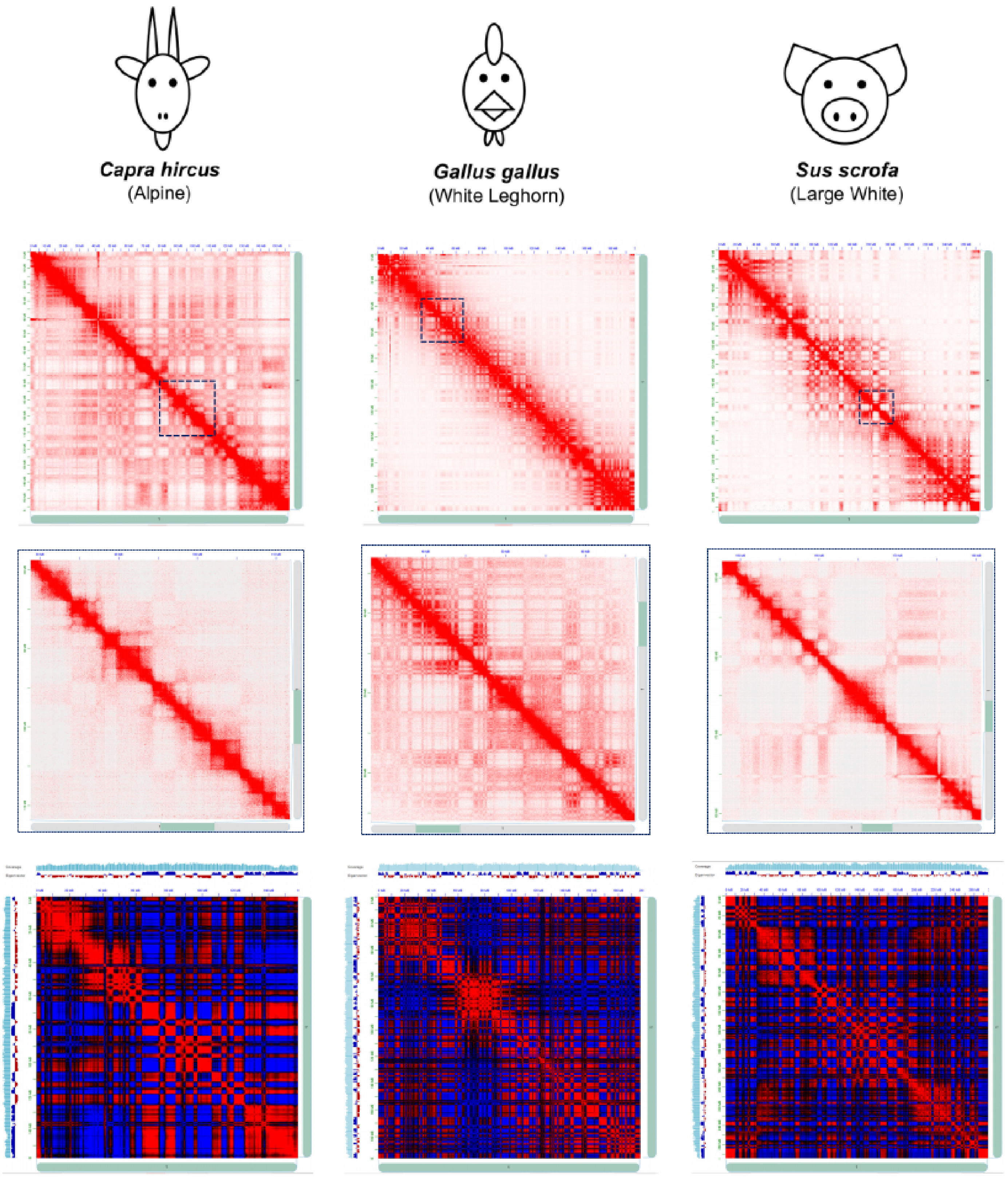


Figure S25: **Juicebox Hi-C maps of three livestock species' chromosome 1.** Each column represents a different species (goat, chicken, pig). For a given column, the first map represents the whole chromosome 1 Hi-C heatmap at 500Kb resolution. On this map the blue square represents a zone that is further enlargeg on the second map, itself representing part of chromosome 1 Hi-C heatmap at 40Kb resolution. The third map represents the chromosome 1 bin pair Pearson correlation matrix computed from the 500Kb resolution Hi-C matrix, and that allows to see the A and B compartments.

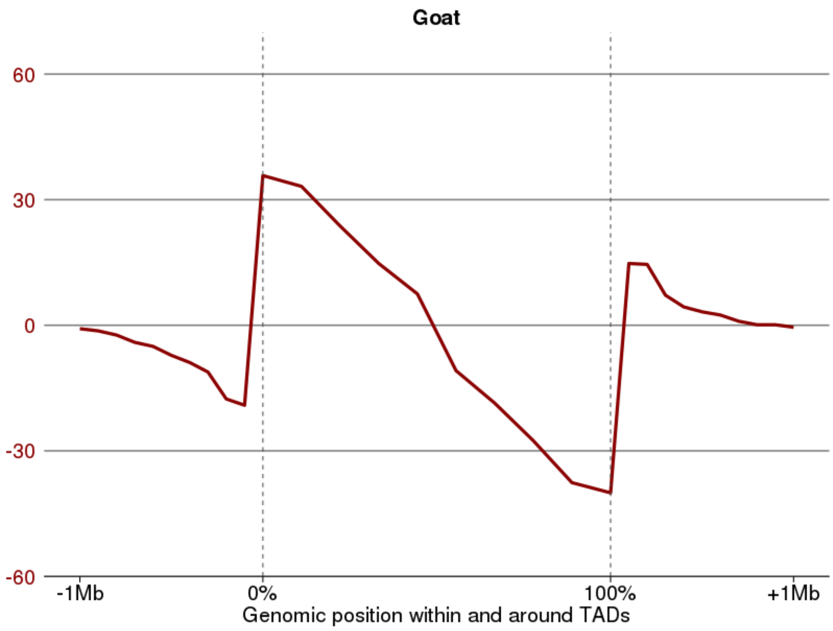


A

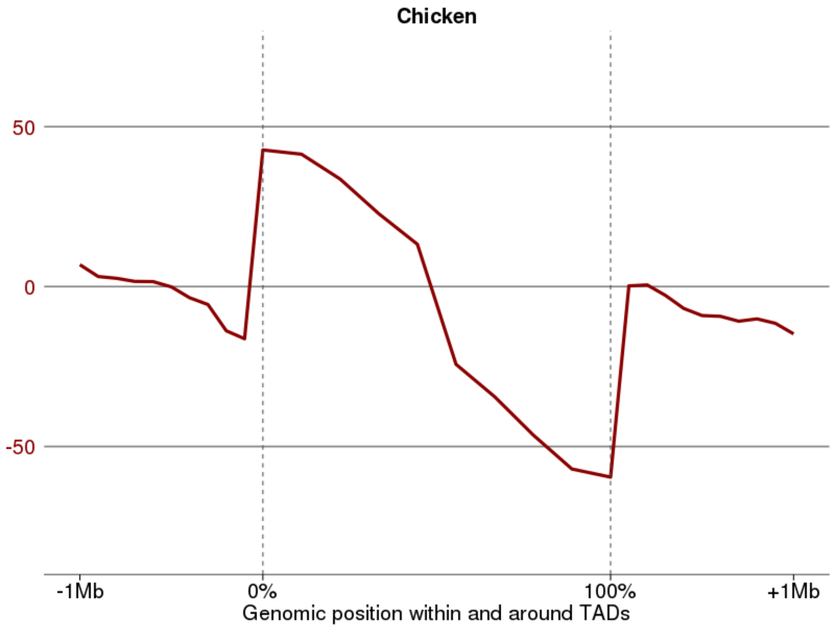


B

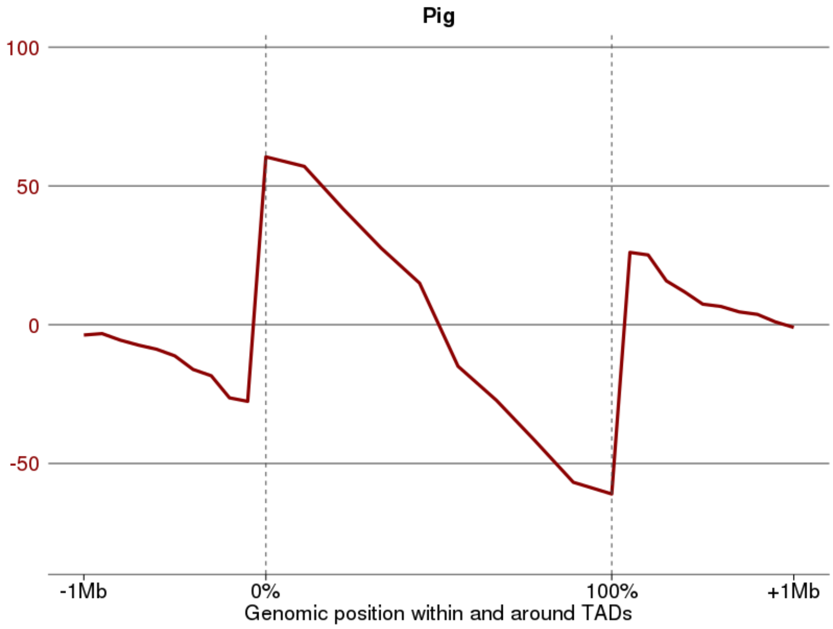


C

Figure S26: **Directionality Index (DI) around Hi-C TADs.**

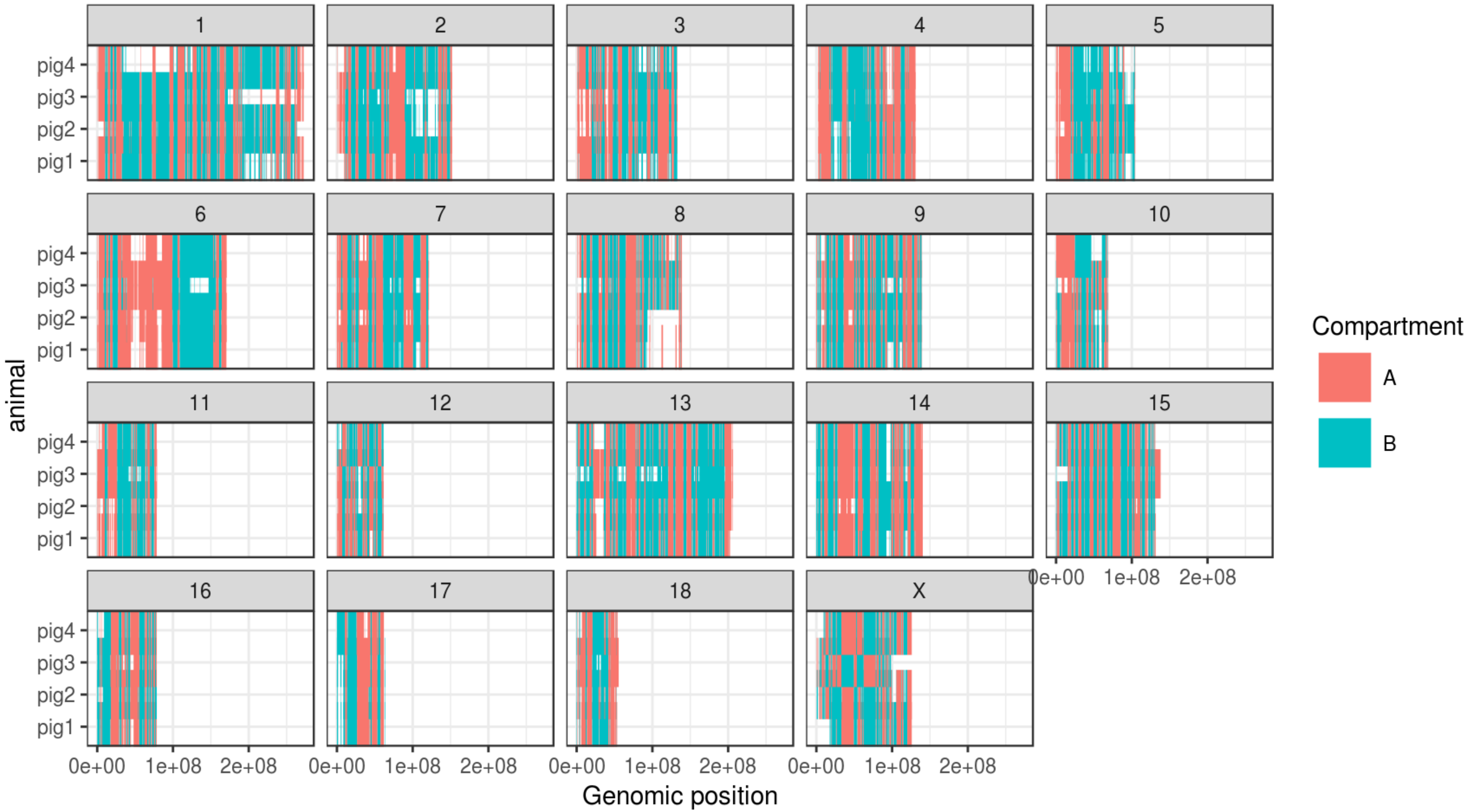


Figure S27: **Distribution of Hi-C A and B compartments along each chromosome and for each animal.** Genome-wide overview of compartment labels per 500Kb bin in pig for each animal. A general coherence can be observed across replicates. White regions are devoid of any called compartment.

Table S1: **Completed experiments.** Available data per animal, sample and experimental assay. Green: completed experiment. Red (NA): not available. Hi-C was attempted on liver samples only, and succeeded on all species but cattle. Consequently, ATAC-seq was not attempted on the cattle liver samples neither.

**RNA-seq**

| **Cattle** | cattle1 | cattle2 | cattle3 | cattle4 | **Goat** | goat1 | goat2 | goat3 | goat4 |
|---|---|---|---|---|---|---|---|---|---|
| cd4 | | | | | cd4 | | | | |
| cd8 | NA | | | | cd8 | | | | |
| liver | | | | | liver | | | | |
| **Chicken** | chicken1 | chicken2 | chicken3 | chicken4 | **Pig** | pig1 | pig2 | pig3 | pig4 |
| cd4 | NA | NA | | | cd4 | NA | | | |
| cd8 | NA | NA | NA | | cd8 | | | | |
| liver | | | | | liver | | | | |

**ATAC-seq**

| **Cattle** | cattle1 | cattle2 | cattle3 | cattle4 | **Goat** | goat1 | goat2 | goat3 | goat4 |
|---|---|---|---|---|---|---|---|---|---|
| cd4 | | | | | cd4 | | NA | | |
| cd8 | | | | | cd8 | | | | |
| liver | NA | NA | NA | NA | liver | | | | |
| **Chicken** | chicken1 | chicken2 | chicken3 | chicken4 | **Pig** | pig1 | pig2 | pig3 | pig4 |
| cd4 | | | | NA | cd4 | | | | |
| cd8 | NA | | NA | NA | cd8 | | | | |
| liver | | | | | liver | | | | NA |

**Hi-C**

| **Cattle** | cattle1 | cattle2 | cattle3 | cattle4 | **Goat** | goat1 | goat2 | goat3 | goat4 |
|---|---|---|---|---|---|---|---|---|---|
| liver | NA | NA | NA | NA | liver | | | | |
| **Chicken** | chicken1 | chicken2 | chicken3 | chicken4 | **Pig** | pig1 | pig2 | pig3 | pig4 |
| liver | | | | | liver | | | | |

Table S2: **Genome and reference gene annotation used for each species.**

| Species | Reference genome assembly | | | | Reference gene annotation | | |
| --- | --- | --- | --- | --- | --- | --- | --- |
| | Version | # fasta sequences | Size (in Gb) | Contig N50 (in Mb) | Source | # genes | # transcripts |
| Cattle | UMD3.1 | 3,317 | 2.67 | 0.1 | Ensembl v90 | 24,616 | 26,740 |
| Goat | CHIR_ARS1 | 29,907 | 2.92 | 26.2 | NCBI vCHIR_ARS1 | 28,931 | 53,266 |
| Chicken | GalGal5 | 23,475 | 1.23 | 2.9 | Ensembl v90 | 24,881 | 38,118 |
| Pig | Sscrofa11.1 | 613 | 2.5 | 48.2 | Ensembl v90 | 25,880 | 49,448 |

Table S3: **Software used in the FR-AgENCODE project.**

| Software name | Software version | Software URL |
|---|---|---|
| R | 3.3.3 | https://www.r-project.org/about.html |
| python | 2.7.2 | https://www.python.org/ |
| samtools | 1.3.1 | http://www.htslib.org/ |
| bedtools | 2.26.0 | http://bedtools.readthedocs.io/en/latest/ |
| kentUtils | 302.1.0 | https://github.com/ENCODE-DCC/kentUtils |
| bwtool | Nov 2015 version | https://github.com/CRG-Barcelona/bwtool/wiki |
| emboss | 6.4.0.0 | http://emboss.sourceforge.net/ |
| fastqc | 0.11.2 | https://www.bioinformatics.babraham.ac.uk/projects/fastqc/ |
| cutadapt | 1.8.3 | http://cutadapt.readthedocs.io/en/stable/guide.html |
| trim_galore | 0.4.0 | https://www.bioinformatics.babraham.ac.uk/projects/trim_galore/ |
| picardtools | 2.1.1 | https://broadinstitute.github.io/picard/ |
| STAR | 2.5.1b | https://github.com/alexdobin/STAR |
| cufflinks | 2.2.1 | http://cole-trapnell-lab.github.io/cufflinks/ |
| RSEM | 1.3.0 | http://deweylab.biostat.wisc.edu/rsem/README.html |
| FEELnc | 0.1.0 | https://github.com/tderrien/FEELnc |
| Bowtie2 | 2.3.3.1 | http://bowtie-bio.sourceforge.net/bowtie2/index.shtml |
| Macs2 | 2.1.1.20160309 | https://chipster.csc.fi/manual/macs2.html |
| fimo | 4.11.1 | http://meme-suite.org/doc/fimo.html |
| HiC-Pro | 2.9.0 | http://nservant.github.io/HiC-Pro/ |
| Armatus | 2.1 | https://www.cs.cmu.edu/~ckingsf/software/armatus/ |
| HiTC | 1.18.1 | https://bioconductor.org/packages/release/bioc/html/HiTC.html |
| Juicer tools | 0.7.5 | https://github.com/theaidenlab/juicer/wiki/Juicer-Tools-Quick-Start |
| Last | 956 | http://last.cbrc.jp/ |

Table S4: **RNA-seq read mapping statistics.** Number and proportion of mapped and uniquely mapped RNA-seq read pairs for each species, replicate and tissue.

| Species | Tissue | Animal | # read pairs | # read pairs mapped | % read pairs mapped | # uniquely mapped read pairs | % uniquely mapped read pairs (out of mapped) | % uniquely mapped read pairs (out of total) |
|---|---|---|---|---|---|---|---|---|
| Cattle | cd4 | cattle1 | 115 398 649 | 111 206 516 | 96,4 | 108 830 546 | 97,9 | 94,3 |
| | cd4 | cattle2 | 118 918 186 | 114 474 132 | 96,3 | 112 127 763 | 98,0 | 94,3 |
| | cd4 | cattle3 | 121 389 996 | 113 765 363 | 93,7 | 108 717 357 | 95,6 | 89,6 |
| | cd4 | cattle4 | 116 083 510 | 112 298 028 | 96,7 | 110 019 926 | 98,0 | 94,8 |
| | cd8 | cattle2 | 117 869 348 | 113 617 513 | 96,4 | 111 375 385 | 98,0 | 94,5 |
| | cd8 | cattle3 | 118 910 270 | 114 458 706 | 96,3 | 111 966 825 | 97,8 | 94,2 |
| | cd8 | cattle4 | 121 883 485 | 117 905 219 | 96,7 | 115 423 917 | 97,9 | 94,7 |
| | liver | cattle1 | 115 371 212 | 111 956 767 | 97,0 | 110 499 020 | 98,7 | 95,8 |
| | liver | cattle2 | 119 870 689 | 116 728 118 | 97,4 | 115 092 798 | 98,6 | 96,0 |
| | liver | cattle3 | 136 710 805 | 132 763 513 | 97,1 | 130 973 598 | 98,7 | 95,8 |
| | liver | cattle4 | 118 889 544 | 115 841 946 | 97,4 | 114 125 584 | 98,5 | 96,0 |
| Goat | cd4 | goat1 | 133 365 348 | 130 269 121 | 97,7 | 123 559 681 | 94,8 | 92,6 |
| | cd4 | goat2 | 132 983 117 | 128 521 637 | 96,6 | 121 198 446 | 94,3 | 91,1 |
| | cd4 | goat3 | 133 831 464 | 129 566 423 | 96,8 | 122 990 885 | 94,9 | 91,9 |
| | cd4 | goat4 | 130 150 700 | 126 874 881 | 97,5 | 121 461 169 | 95,7 | 93,3 |
| | cd8 | goat1 | 130 970 787 | 127 546 631 | 97,4 | 120 904 297 | 94,8 | 92,3 |
| | cd8 | goat2 | 131 654 878 | 128 355 710 | 97,5 | 121 468 794 | 94,6 | 92,3 |
| | cd8 | goat3 | 132 923 610 | 129 276 688 | 97,3 | 123 235 599 | 95,3 | 92,7 |
| | cd8 | goat4 | 131 413 248 | 127 239 980 | 96,8 | 121 762 207 | 95,7 | 92,7 |
| | liver | goat1 | 112 623 152 | 110 432 315 | 98,1 | 105 943 818 | 95,9 | 94,1 |
| | liver | goat2 | 120 492 856 | 117 992 580 | 97,9 | 112 453 956 | 95,3 | 93,3 |
| | liver | goat3 | 133 931 176 | 131 150 414 | 97,9 | 126 458 023 | 96,4 | 94,4 |
| | liver | goat4 | 125 330 385 | 122 259 872 | 97,6 | 118 111 570 | 96,6 | 94,2 |
| Chicken | cd4 | chicken3 | 91 907 306 | 86 645 024 | 94,3 | 83 374 510 | 96,2 | 90,7 |
| | cd4 | chicken4 | 90 843 387 | 79 772 579 | 87,8 | 76 778 803 | 96,2 | 84,5 |
| | cd8 | chicken4 | 90 476 514 | 82 431 332 | 91,1 | 79 685 312 | 96,7 | 88,1 |
| | liver | chicken1 | 119 681 527 | 114 944 560 | 96,0 | 111 122 525 | 96,7 | 92,8 |
| | liver | chicken2 | 107 777 068 | 103 770 521 | 96,3 | 100 787 680 | 97,1 | 93,5 |
| | liver | chicken3 | 121 616 481 | 116 940 968 | 96,2 | 113 315 177 | 96,9 | 93,2 |
| | liver | chicken4 | 124 089 440 | 119 195 187 | 96,1 | 115 442 821 | 96,9 | 93,0 |
| Pig | cd4 | pig2 | 116 273 646 | 113 997 546 | 98,0 | 109 051 735 | 95,7 | 93,8 |
| | cd4 | pig3 | 118 260 498 | 115 042 891 | 97,3 | 110 755 493 | 96,3 | 93,7 |
| | cd4 | pig4 | 114 159 672 | 111 934 574 | 98,1 | 108 379 228 | 96,8 | 94,9 |
| | cd8 | pig1 | 231 339 734 | 225 402 022 | 97,4 | 219 199 568 | 97,2 | 94,8 |
| | cd8 | pig2 | 116 383 341 | 114 513 826 | 98,4 | 110 146 147 | 96,2 | 94,6 |
| | cd8 | pig3 | 116 696 412 | 113 825 826 | 97,5 | 110 613 640 | 97,2 | 94,8 |
| | cd8 | pig4 | 117 449 531 | 114 751 230 | 97,7 | 108 598 741 | 94,6 | 92,5 |
| | liver | pig1 | 116 242 745 | 113 660 238 | 97,8 | 110 849 470 | 97,5 | 95,4 |
| | liver | pig2 | 117 818 284 | 115 364 464 | 97,9 | 111 627 222 | 96,8 | 94,7 |
| | liver | pig3 | 110 567 402 | 108 214 816 | 97,9 | 105 590 955 | 97,6 | 95,5 |
| | liver | pig4 | 114 769 769 | 112 137 997 | 97,7 | 109 296 648 | 97,5 | 95,2 |

Table S5: **Differentially Expressed (DE) reference genes.** Number of differentially expressed reference genes obtained by the two statistical models (see main text and Methods)

| Differential analysis model | Tissue 1 | Tissue 2 | log(Tissue2 / Tissue1) | Number of DE reference genes | | | |
|---|---|---|---|---|---|---|---|
| | | | | Cattle | Goat | Chicken | Pig |
| 1 (tissue pairs) | cd4 | cd8 | >0 | 751 | 1 098 | 1 504 | 867 |
| | cd4 | cd8 | <0 | 463 | 748 | 592 | 604 |
| | cd4 | liver | >0 | 5 048 | 6 303 | 5 112 | 5 725 |
| | cd4 | liver | <0 | 3 887 | 4 304 | 3 351 | 3 653 |
| | cd8 | liver | >0 | 4 890 | 6 134 | 4 146 | 5 665 |
| | cd8 | liver | <0 | 3 911 | 4 397 | 3 087 | 3 792 |
| 2 (tcell vs liver) | cd | liver | >0 | 4 992 | 6 188 | 4 307 | 5 666 |
| | cd | liver | <0 | 3 943 | 4 384 | 2 640 | 3 772 |

Table S6: **Genes consistently over-expressed in CD4+ compared to CD8+ or reciprocally, in four livestock species.** For the 39 genes consistently seen as over-expressed in CD4+ with respect to CD8+ (10 genes) or reciprocally (29 genes), are indicated: the human gene ID, the gene name in a column that indicates the cell type in which the gene is over-expressed, the TPM in human CD4+ and CD8+ cells from the Blueprint project (CD4-positive, alpha-beta T cell and CD8-positive, alpha-beta T cell respectively), whether or not these expression levels are consistent with the differential behavior observed in livestock, the expression in livestock CD4+ and CD8+ (average across samples) and a reference describing the role of the gene in blood cells when available.

| Human gene id | Gene name (in column corresponding to over-expression) | | TPM in human | | Consistency between livestock and human (FC>1) | TPM in cattle | | TPM in goat | | TPM in chicken | | TPM in pig | |
|---|---|---|---|---|---|---|---|---|---|---|---|---|---|
| | CD4+ | CD8+ | CD4+ | CD8+ | | CD4+ | CD8+ | CD4+ | CD8+ | CD4+ | CD8+ | CD4+ | CD8+ |
| ENSG00000102245 [1] | CD40LG | . | 27.0 | 0.5 | yes | 97.9 | 7.6 | 51.6 | 10.5 | 164.8 | 7.5 | 153.4 | 28.7 |
| ENSG00000106537 | TSPAN13 | . | 0.4 | 0.5 | no | 118.8 | 62.6 | 93.9 | 32.6 | 4.0 | 0.5 | 3.4 | 1.0 |
| ENSG00000107485 [2] | GATA3 | . | 6.0 | 4.0 | yes | 37.7 | 11.5 | 43.3 | 18.8 | 113.5 | 7.6 | 51.1 | 30.0 |
| ENSG00000115602 | IL1RL1 | . | NA | 0.1 | NA | 4.7 | 0.3 | 5.2 | 2.2 | 5.5 | 0.2 | 4.2 | 1.0 |
| ENSG00000126353 [3] | CCR7 | . | 312.0 | 102.0 | yes | 33.7 | 11.3 | 164.5 | 44.6 | 96.2 | 21.7 | 87.1 | 33.2 |
| ENSG00000130396 | AFDN | . | 0.9 | 0.3 | yes | 6.6 | 2.9 | 17.8 | 2.8 | 3.3 | 0.8 | 2.4 | 0.1 |
| ENSG00000163599 [4] | CTLA4 | . | 39.0 | 2.0 | yes | 6.1 | 0.6 | 19.8 | 1.4 | 635.6 | 16.9 | 15.0 | 2.2 |
| ENSG00000178199 | ZC3H12D | . | 17.0 | 6.0 | yes | 30.5 | 13.6 | 45.7 | 9.4 | 1,384.7 | 134.7 | 26.3 | 4.5 |
| ENSG00000178562 [5] | CD28 | . | 161.0 | 74.0 | yes | 34.4 | 9.3 | 91.5 | 11.0 | 208.3 | 14.8 | 107.4 | 27.6 |
| ENSG00000178573 [6] | MAF | . | 3.0 | 1.0 | yes | 13.2 | 1.5 | 11.8 | 5.4 | 6.8 | 0.3 | 15.0 | 2.5 |
| ENSG00000139567 | . | ACVRL1 | NA | NA | NA | 5.9 | 48.0 | 5.0 | 18.9 | 5.2 | 10.1 | 0.0 | 0.2 |
| ENSG00000169252 [7] | . | ADRB2 | 0.6 | 2.0 | yes | 3.9 | 17.9 | 0.6 | 7.3 | 57.8 | 219.3 | 2.7 | 15.2 |
| ENSG00000156966 | . | B3GNT7 | 0.1 | 0.2 | yes | 1.7 | 7.2 | 0.2 | 0.7 | 0.0 | 2.8 | 0.0 | 1.4 |
| ENSG00000136573 | . | BLK | 1.0 | 4.0 | yes | 15.9 | 27.0 | 3.8 | 53.2 | 3.9 | 15.4 | 1.5 | 32.8 |
| ENSG00000182985 | . | CADM1 | 0.2 | NA | NA | 4.1 | 21.4 | 1.1 | 3.5 | 1.6 | 3.9 | 0.0 | 0.2 |
| ENSG00000271503 [8] | . | CCL5 | 8.0 | 38.0 | yes | 33.0 | 1,205.8 | 16.2 | 2,009.6 | 1.2 | 102.1 | 190.2 | 2,465.9 |
| ENSG00000172116 [9] | . | CD8B | 2.0 | 210.0 | yes | 9.7 | 260.7 | 17.6 | 514.5 | 0.8 | 173.0 | 1.1 | 115.4 |
| ENSG00000109943 [10] | . | CRTAM | 3.0 | 54.0 | yes | 2.3 | 16.3 | 4.0 | 16.2 | 0.8 | 52.0 | 3.1 | 37.6 |
| ENSG00000100592 | . | DAAM1 | 4.0 | 3.0 | no | 5.3 | 5.0 | 2.1 | 2.8 | 0.3 | 0.3 | 5.5 | 5.7 |
| ENSG00000035664 [11] | . | DAPK2 | 0.3 | 0.9 | yes | 1.8 | 8.2 | 0.9 | 10.1 | 3.0 | 11.7 | 14.9 | 60.1 |
| ENSG00000213853 | . | EMP2 | 0.6 | 0.4 | no | 0.1 | 0.7 | 1.9 | 6.9 | 0.1 | 0.5 | 0.1 | 0.9 |
| ENSG00000163508 [12] | . | EOMES | 0.9 | 1.0 | yes | 1.0 | 11.3 | 1.0 | 17.5 | 2.9 | 133.4 | 1.0 | 12.5 |
| ENSG00000139132 | . | FGD4 | 0.2 | 2.0 | yes | 3.3 | 8.5 | 0.9 | 2.3 | 0.9 | 6.8 | 0.2 | 5.3 |
| ENSG00000160219 | . | GAB3 | 4.0 | 4.0 | no | 3.3 | 8.6 | 9.2 | 25.1 | 4.4 | 38.4 | 8.9 | 27.2 |
| ENSG00000125245 | . | GPR18 | 17.0 | 26.0 | yes | 7.8 | 22.8 | 9.0 | 22.8 | 13.5 | 104.8 | 39.0 | 297.6 |
| ENSG00000100385 [13] | . | IL2RB | 7.0 | 9.0 | yes | 42.7 | 131.6 | 134.9 | 573.8 | 30.2 | 190.7 | 62.6 | 205.2 |
| ENSG00000157404 | . | KIT | 0.9 | 2.0 | yes | 2.4 | 7.9 | 2.4 | 6.4 | 0.1 | 0.3 | 0.0 | 1.2 |
| ENSG00000086730 [14] | . | LAT2 | 0.8 | 4.0 | yes | 46.1 | 156.3 | 20.0 | 46.2 | 2.0 | 11.9 | 22.1 | 123.7 |
| ENSG00000189067 | . | LITAF | 68.0 | 58.0 | no | 24.0 | 98.1 | 26.0 | 82.3 | 12.3 | 41.5 | 4.6 | 21.4 |
| ENSG00000158186 | . | MRAS | 0.1 | NA | NA | 0.9 | 5.9 | 2.1 | 6.4 | 16.4 | 32.9 | 0.0 | 0.0 |
| ENSG00000100311 | . | PDGFB | 0.9 | NA | NA | 6.1 | 25.8 | 2.2 | 8.6 | 0.2 | 0.9 | 0.7 | 5.6 |
| ENSG00000115956 | . | PLEK | 2.0 | 22.0 | yes | 10.4 | 32.1 | 9.1 | 52.4 | 36.8 | 43.0 | 5.2 | 55.9 |
| ENSG00000141956 | . | PRDM15 | 5.0 | 2.0 | no | 1.8 | 1.8 | 3.4 | 3.0 | 7.6 | 3.3 | 3.4 | 3.4 |
| ENSG00000198915 | . | RASGEF1A | 0.1 | 0.1 | no | 3.3 | 8.3 | 2.5 | 10.8 | 27.6 | 61.8 | 0.3 | 3.0 |
| ENSG00000119729 | . | RHOQ | 6.0 | 4.0 | no | 18.1 | 74.3 | 14.5 | 45.9 | 0.6 | 8.9 | 2.9 | 16.5 |
| ENSG00000136158 [15] | . | SPRY2 | NA | 1.0 | yes | 1.0 | 23.5 | 0.6 | 11.3 | 2.0 | 66.0 | 0.1 | 40.5 |
| ENSG00000204634 | . | TBC1D8 | 0.6 | 2.0 | yes | 12.6 | 62.9 | 5.8 | 17.0 | 0.3 | 0.6 | 0.2 | 0.5 |
| ENSG00000070759 | . | TESK2 | 6.0 | 6.0 | no | 4.8 | 15.2 | 6.9 | 29.1 | 3.3 | 9.5 | 5.6 | 39.0 |
| ENSG00000165914 | . | TTC7B | 0.4 | 0.5 | yes | 0.4 | 2.0 | 0.2 | 0.7 | 1.0 | 2.5 | 0.1 | 0.6 |

### Table S7: **FR-AgENCODE transcript positional classification.**

| Species | Total # | known | | extension | | alternative | | novel | |
|---|---|---|---|---|---|---|---|---|---|
| | | # | % of total | # | % of total | # | % of total | # | % of total |
| Cattle | 84,971 | 11,736 | **13.8** | 2,500 | 2.9 | 40,813 | **48.0** | 29,922 | **35.2** |
| Goat | 78,091 | 29,520 | **37.8** | 2,583 | 3.3 | 28,891 | 37.0 | 17,097 | 21.9 |
| Chicken | 57,817 | 15,896 | 27.5 | 3,018 | 5.2 | 28,065 | **48.5** | 10,838 | 18.7 |
| Pig | 77,540 | 23,921 | 30.8 | 2,702 | 3.5 | 35,062 | **45.2** | 15,855 | 20.4 |

| Species | known.mRNA | | known.lncRNA | | known.otherRNA | |
|---|---|---|---|---|---|---|
| | # | % of known | # | % of known | # | % of known |
| Cattle | 11,576 | **98.6** | 13 | 0.1 | 147 | 1.3 |
| Goat | 26,973 | **91.4** | 1,163 | 3.9 | 1,384 | **4.7** |
| Chicken | 14,765 | **92.9** | 882 | **5.5** | 249 | 1.6 |
| Pig | 23,701 | **99.1** | 96 | 0.4 | 124 | 0.5 |

| Species | extension.mRNA | | extension.lncRNA | | extension.otherRNA | |
|---|---|---|---|---|---|---|
| | # | % of extension | # | % of extension | # | % of extension |
| Cattle | 2,497 | **99.9** | 0 | 0.0 | 3 | 0.1 |
| Goat | 2,351 | **91.0** | 100 | 3.9 | 132 | **5.1** |
| Chicken | 2,982 | **98.8** | 33 | 1.1 | 3 | 0.1 |
| Pig | 2,694 | **99.7** | 5 | 0.2 | 3 | 0.1 |

| Species | alternative.mRNA | | alternative.lncRNA | | alternative.otherRNA | |
|---|---|---|---|---|---|---|
| | # | % of alternative | # | % of alternative | # | % of alternative |
| Cattle | 40,770 | **99.9** | 0 | 0.0 | 43 | 0.1 |
| Goat | 26,554 | **91.9** | 984 | 3.4 | 1,353 | **4.7** |
| Chicken | 27,470 | **97.9** | 399 | 1.4 | 196 | 0.7 |
| Pig | 34,822 | **99.3** | 226 | 0.6 | 14 | 0.0 |

| Species | novel.mRNA | | novel.lncRNA | | novel.otherRNA | |
|---|---|---|---|---|---|---|
| | # | % of novel | # | % of novel | # | % of novel |
| Cattle | 4,958 | 16.6 | 22,711 | 75.9 | 2,253 | **7.5** |
| Goat | 2,949 | **17.2** | 11,617 | 67.9 | 2,531 | **14.8** |
| Chicken | 2,350 | 21.7 | 6,797 | 62.7 | 1,691 | **15.6** |
| Pig | 2,504 | 15.8 | 12,284 | 77.5 | 1,067 | **6.7** |

Table S8: **FR-AgENCODE novel coding genes and their orthology with human**

| Species | # FR-AgENCODE genes | # novel FR-AgENCODE genes | # novel coding FR-AgENCODE genes | that can be projected to a human gene | |
|---|---|---|---|---|---|
| | | | | # | % |
| Cattle | 34,296 | 19,324 | 15,611 | 166 | 12.3 |
| Goat | 28,537 | 11,545 | 16,437 | 85 | 6.4 |
| Chicken | 20,408 | 6,597 | 13,149 | 78 | 15.5 |
| Pig | 24,570 | 10,238 | 14,054 | 38 | 9.5 |

Table S9: **Differentially Expressed (DE) FR-AgENCODE genes.** Number of differentially expressed FR-AgENCODE genes obtained by the two statistical models (see main text and Methods).

| Differential analysis model | Tissue 1 | Tissue 2 | log(Tissue2 / Tissue1) | Number of DE FR-AgENCODE genes | | | |
|---|---|---|---|---|---|---|---|
| | | | | Cattle | Goat | Chicken | Pig |
| 1 (tissue pairs) | cd4 | cd8 | >0 | 1 450 | 1 614 | 1 940 | 1 211 |
| | cd4 | cd8 | <0 | 923 | 1 056 | 974 | 792 |
| | cd4 | liver | >0 | 12 231 | 11 782 | 7 426 | 9 709 |
| | cd4 | liver | <0 | 10 311 | 7 152 | 5 134 | 6 386 |
| | cd8 | liver | >0 | 11 810 | 11 542 | 5 899 | 9 880 |
| | cd8 | liver | <0 | 10 573 | 7 408 | 4 331 | 6 782 |
| 2 (tcell vs liver) | cd | liver | >0 | 12 218 | 11 773 | 6 246 | 10 040 |
| | cd | liver | <0 | 10 818 | 7 448 | 3 929 | 6 910 |

Table S10: **lncRNA classification.** Number of classified expressed lncRNAs per species. This table includes monoexonic lncRNAs that represent 57-68% of these transcripts. Loci are bracketed. Fields with an asterisk (*) indicate the existence of lncRNAs unclassified by FEELnc because they are on unassembled contigs; they represent 217, 717, 2,718 and 83 lncRNAs in cattle, goat, chicken and pig respectively and are not listed here.

| | | Intergenic lncRNAs (loci) – 88-94% | | | | | Genic lncRNAs (loci) – 6-12% | | |
|---|---|---|---|---|---|---|---|---|---|
| | Total (transcript) | Same strand | | Divergent | | Convergent | Exonic antisense | Intronic antisense | Intronic sense |
| | | upstr. | downstr. | ≤1kb | >1kb | | | | |
| Cattle | 22724 | 5522 (4671) | 6438 (5460) | 1450 (1070) | 4494 (3588) | 3328 (2617) | 1235 (851) | 241 (173) | 16 (16) |
| Goat | 13864 | 2431 (1940) | 4050 (3312) | 1744 (1241) | 2908 (2233) | 1966 (1496) | 437 (220) | 219 (124) | 109 (89) |
| Chicken | 7502* | 1618 (1390) | 2045 (1728) | 687 (539) | 1194 (983) | 1093 (903) | 646 (510) | 157 (124) | 62 (56) |
| Pig | 12587* | 2508 (2100) | 3539 (3075) | 1117 (810) | 2466 (2060) | 1746 (1436) | 935 (669) | 216 (178) | 60 (58) |

Table S11: **FR-AgENCODE transcript coding classification.**

| Species | Total # | mRNA | | lncRNA | | otherRNA | |
|---|---|---|---|---|---|---|---|
| | | # | % ot total | # | % ot total | # | % ot total |
| Cattle | 84,971 | 59,801 | **70.4** | 22,724 | **26.7** | 2,446 | 2.9 |
| Goat | 78,091 | 58,827 | **75.3** | 13,864 | 17.8 | 5,400 | 6.9 |
| Chicken | 57,817 | 47,567 | **82.3** | 8,111 | 14.0 | 2,139 | 3.7 |
| Pig | 77,540 | 63,721 | **82.2** | 12,611 | 16.3 | 1,208 | 1.6 |

| Species | mRNA.known | | mRNA.extension | | mRNA.alternative | | mRNA.novel | |
|---|---|---|---|---|---|---|---|---|
| | # | % of mRNA | # | % of mRNA | # | % of mRNA | # | % of mRNA |
| Cattle | 11,576 | 19.4 | 2,497 | 4.2 | 40,770 | **68.2** | 4,958 | 8.3 |
| Goat | 26,973 | **45.9** | 2,351 | 4.0 | 26,554 | **45.1** | 2,949 | 5.0 |
| Chicken | 14,765 | 31.0 | 2,982 | 6.3 | 27,470 | **57.8** | 2,350 | 4.9 |
| Pig | 23,701 | 37.2 | 2,694 | 4.2 | 34,822 | **54.6** | 2,504 | 3.9 |

| Species | lncRNA.known | | lncRNA.extension | | lncRNA.alternative | | lncRNA.novel | |
|---|---|---|---|---|---|---|---|---|
| | # | % of lncRNA | # | % of lncRNA | # | % of lncRNA | # | % of lncRNA |
| Cattle | 13 | 0.1 | 0 | 0.0 | 0 | 0.0 | 22,711 | **99.9** |
| Goat | 1,163 | **8.4** | 100 | 0.7 | 984 | **7.1** | 11,617 | **83.8** |
| Chicken | 882 | **10.9** | 33 | 0.4 | 399 | **4.9** | 6,797 | **83.8** |
| Pig | 96 | 0.8 | 5 | 0.0 | 226 | 1.8 | 12,284 | **97.4** |

| Species | otherRNA.known | | otherRNA.extension | | otherRNA.alternative | | otherRNA.novel | |
|---|---|---|---|---|---|---|---|---|
| | # | % of otherRNA | # | % of otherRNA | # | % of otherRNA | # | % of otherRNA |
| Cattle | 147 | 6.0 | 3 | 0.1 | 43 | 1.8 | 2,253 | **92.1** |
| Goat | 1,384 | **25.6** | 132 | 2.4 | 1,353 | **25.1** | 2,531 | 46.9 |
| Chicken | 249 | 11.6 | 3 | 0.1 | 196 | 9.2 | 1,691 | **79.1** |
| Pig | 124 | 10.3 | 3 | 0.2 | 14 | 1.2 | 1,067 | **88.3** |

Table S12: **Number of ATAC-seq peaks per species.**

| Species | Tissue | # ATAC-seq peaks |
|---|---|---|
| Cattle | cd4 | 69,661 |
| | cd8 | 75,295 |
| | merged | 104,985 |
| Goat | cd4 | 39,526 |
| | cd8 | 57,084 |
| | liver | 14,137 |
| | merged | 74,805 |
| Chicken | cd4 | 38,594 |
| | cd8 | 49,962 |
| | liver | 75,305 |
| | merged | 119,894 |
| Pig | cd4 | 80,745 |
| | cd8 | 111,457 |
| | liver | 25,885 |
| | merged | 149,333 |

Table S13: **Number of differentially accessible (DA) ATAC-seq peaks per species.**

| Tissue 1 | Tissue 2 | log(Tissue2/Tissue1) | Species | Number of DA ATAC-seq peaks |
|---|---|---|---|---|
| T cell | Liver | >0 | Goat | 2,780 |
| | | <0 | | 2,042 |
| | | >0 | Chicken | 6,663 |
| | | <0 | | 6,991 |
| | | >0 | Pig | 5,467 |
| | | <0 | | 3,678 |

Table S14: **Hi-C read pair mapping statistics.** Number of read pairs of different categories. *Initial*: total number of sequenced read pairs. *Reported*: pairs with both reads mapped on the genome. *Valid*: uniquely mapped pairs with an estimated insert size (sum of the distances from the reads to their next downstream HindIII genomic sites) between 20bp and 1Kb. *Valid.rmdup*: valid read pairs after duplication removal that were used to build the interaction matrices. *Trans*: pairs with reads on different chromosomes.

| Species | Animal | initial | reported | valid | valid.rmdup | trans |
|---|---|---|---|---|---|---|
| Goat | goat1 | 192,807,889 | 164,130,417 | 94,760,148 | 77,324,040 | 31,728,784 |
| | goat2 | 184,098,994 | 142,479,273 | 46,280,100 | 36,346,806 | 12,858,516 |
| | goat3 | 192,081,174 | 149,494,649 | 44,712,829 | 37,115,258 | 13,227,700 |
| | goat4 | 178,929,203 | 135,758,984 | 38,435,525 | 31,232,835 | 12,691,582 |
| Chicken | chicken1 | 172,356,821 | 136,854,885 | 100,067,668 | 74,551,182 | 35,227,545 |
| | chicken2 | 182,654,001 | 152,646,931 | 122,151,951 | 86,044,085 | 30,151,392 |
| | chicken3 | 193,520,830 | 149,648,213 | 89,183,496 | 64,824,211 | 19,603,078 |
| | chicken4 | 187,696,586 | 131,460,531 | 51,115,059 | 37,069,637 | 11,768,987 |
| Pig | pig1 | 168,050,522 | 139,712,050 | 111,642,484 | 82,782,023 | 25,956,367 |
| | pig2 | 157,480,346 | 129,447,277 | 95,004,214 | 70,483,985 | 19,076,359 |
| | pig3 | 165,285,596 | 132,403,825 | 93,115,687 | 74,144,797 | 23,560,745 |
| | pig4 | 165,529,922 | 131,590,998 | 82,504,226 | 62,212,375 | 18,705,743 |

Table S15: **Statistics of Hi-C TADs and A/B compartments.**

| Feature | Species | Number | Min size | Mean size | Max size | Genomic coverage (Mb) | Genomic coverage (%) |
|---|---|---|---|---|---|---|---|
| TADs | Pig | 10,982 | 80,000 | 184,600 | 4,520,000 | 2,027.28 | 83.25 |
| | Chicken | 5,362 | 80,000 | 148,100 | 3,520,000 | 794.28 | 79.4 |
| | Goat | 8,990 | 80,000 | 219,800 | 6,680,000 | 1,975.96 | 85.92 |
| A/B comp. | Pig | 698 | 343,300 | 3,175,000 | 44,000,000 | 2,215.84 | 90.51 |
| | Chicken | 578 | 95,050 | 1,596,000 | 10,000,000 | 922.348 | 92.21 |
| | Goat | 616 | 426,000 | 3,412,000 | 21,000,000 | 2,101.87 | 91.4 |
